# Dynamics of cortical contrast adaptation predict perception of signals in noise

**DOI:** 10.1101/2021.08.11.455845

**Authors:** Christopher F. Angeloni, Wiktor Młynarski, Eugenio Piasini, Aaron M. Williams, Katherine C. Wood, Linda Garami, Ann M. Hermundstad, Maria N. Geffen

## Abstract

Neurons throughout the sensory pathway adapt their responses depending on the statistical structure of the sensory environment. Contrast gain control is a form of adaptation in the auditory cortex, but it is unclear whether the dynamics of gain control reflect efficient adaptation, and whether they shape behavioral perception. Here, we trained mice to detect a target presented in background noise shortly after a change in the contrast of the background. The observed changes in cortical gain and behavioral detection followed the dynamics of a normative model of efficient contrast gain control; specifically, target detection and sensitivity improved slowly in low contrast, but degraded rapidly in high contrast. Auditory cortex was required for this task, and cortical responses were not only similarly affected by contrast but predicted variability in behavioral performance. Combined, our results demonstrate that dynamic gain adaptation supports efficient coding in auditory cortex and predicts the perception of sounds in noise.

## Introduction

As we navigate the world around us, the statistics of the environment can change dramatically. In order to maintain stable percepts, the nervous system adapts to persistent statistical properties of sensory inputs. The efficient coding hypothesis postulates that the nervous system accomplishes this by matching the limited dynamic range of individual neurons to the statistics of incoming sensory signals^1^, thereby efficiently encoding information within many types of environments^2–4^. Indeed, adaptation to environmental statistics has been found in many sensory modalities and species^5–13^. In the auditory system, neurons exhibit contrast gain control, adapting the gain of their response function to match the variability in level (contrast) of the incoming sounds^14–19^. Yet, it is unknown whether and how the dynamics of contrast gain control in the auditory system inform perception, as the dynamics of neuronal adaptation have not previously been measured simultaneously with behavior. The goal of our study was to test if the dynamics of contrast gain control in auditory cortex predict changes in the perception of targets embedded in noise.

Contrast gain control has been proposed as a mechanism for creating sensory representations of sounds that are invariant to background noise^20^. In the ferret auditory pathway, representations of sounds become more noise-tolerant in higher auditory areas, and the amount of noise-tolerance correlates with the strength of adaptation to the stimulus statistics, suggesting that level adaptation and gain control enhance the encoding of stimuli embedded in stationary noise^16^. Additionally, recent psychophysical studies found that perception in noise is altered by efficient adaptation to stimulus statistics. In humans, target level discriminability is greater in low contrast than in high contrast, an effect consistent with gain control observed in primary auditory cortex^19^. Similar relationships between gain control and behavioral percepts of sound location have also been found in ferrets^10^ and guinea pigs^21^. However, in these previous studies, the measurement of psychophysical performance and neuronal gain control were performed separately in human and animal subjects, respectively, so it remains unclear how variability in adaptation over sessions or subjects relates to behavioral performance. Additionally, these previous studies focused primarily on behavioral performance after complete adaptation to stimulus contrast, whereas recent work has highlighted the dynamical nature of efficient adaptation^22, 23^. In the current study, we aimed to assess whether and how the dynamics of contrast gain control reflected efficient adaptation and predicted behavioral performance across subjects.

First, we established a normative framework to model the neuronal dynamics of gain control and predict how efficient contrast adaptation should affect the detection of signals in noise. We then derived a procedure for estimating moment-to-moment changes in neuronal gain based on generalized linear models (GLM) and found that the dynamics of gain control in auditory cortex were asymmetric, as observed in the normative model. Next, to test the role of contrast gain control in auditory behavior, we trained mice to detect targets after a change in background contrast. We found that contrast-induced changes in behavioral target detection threshold, sensitivity, and behavioral adaptation dynamics followed the normative model predictions. Furthermore, we found that auditory cortex was necessary for detection in the presence of a background, but not for detection in silence, suggesting a distinct role of auditory cortex in separating targets from the background. Building on this finding, we found that the dynamics of cortical encoding of targets resembled the normative model predictions and observed behavioral adaptation, and that population activity in the auditory cortex predicted individual variability in behavioral performance. Finally, we estimated cortical gain during the task, finding that variability in neuronal gain predicted variability in task performance. Combined, our results identify a predictive relationship between efficient adaptation via gain control and detection of signals in noise, and provide a normative framework to predict the dynamics of behavioral performance in changing sensory environments.

## Results

### Target-in-background detection task and normative model for task predictions

To assess how behavioral performance is impacted by adaptation to stimulus contrast, we first devised a GO/NO-GO task in which mice were trained to detect targets embedded in switching low and high contrast backgrounds. During each trial, the mouse was presented with dynamic random chords (DRCs) of one contrast, which switched after 3 s to the other contrast. At variable delays after the contrast switch, broad- band target chords were superimposed on the background chords, and mice were trained to lick for a water reward upon hearing the target (henceforth, we refer to high-to-low contrast trials as “low contrast” and low-to-high contrast trials as “high contrast”, referring to the contrast where mice detected targets). Target trials were interleaved with background-only trials, in which the mouse was trained to withhold licking, but would receive a 7s timeout for licking after the contrast switch (Figure 1a,b). To assess behavioral sensitivity to targets, we parametrically varied target level in each contrast, and to assess behavioral adaptation, we parametrically varied target timing (Figure 1c). This stimulus design allowed us to quantitatively test how the dynamics of adaptation to background contrast affect perception.

**Figure 1.**
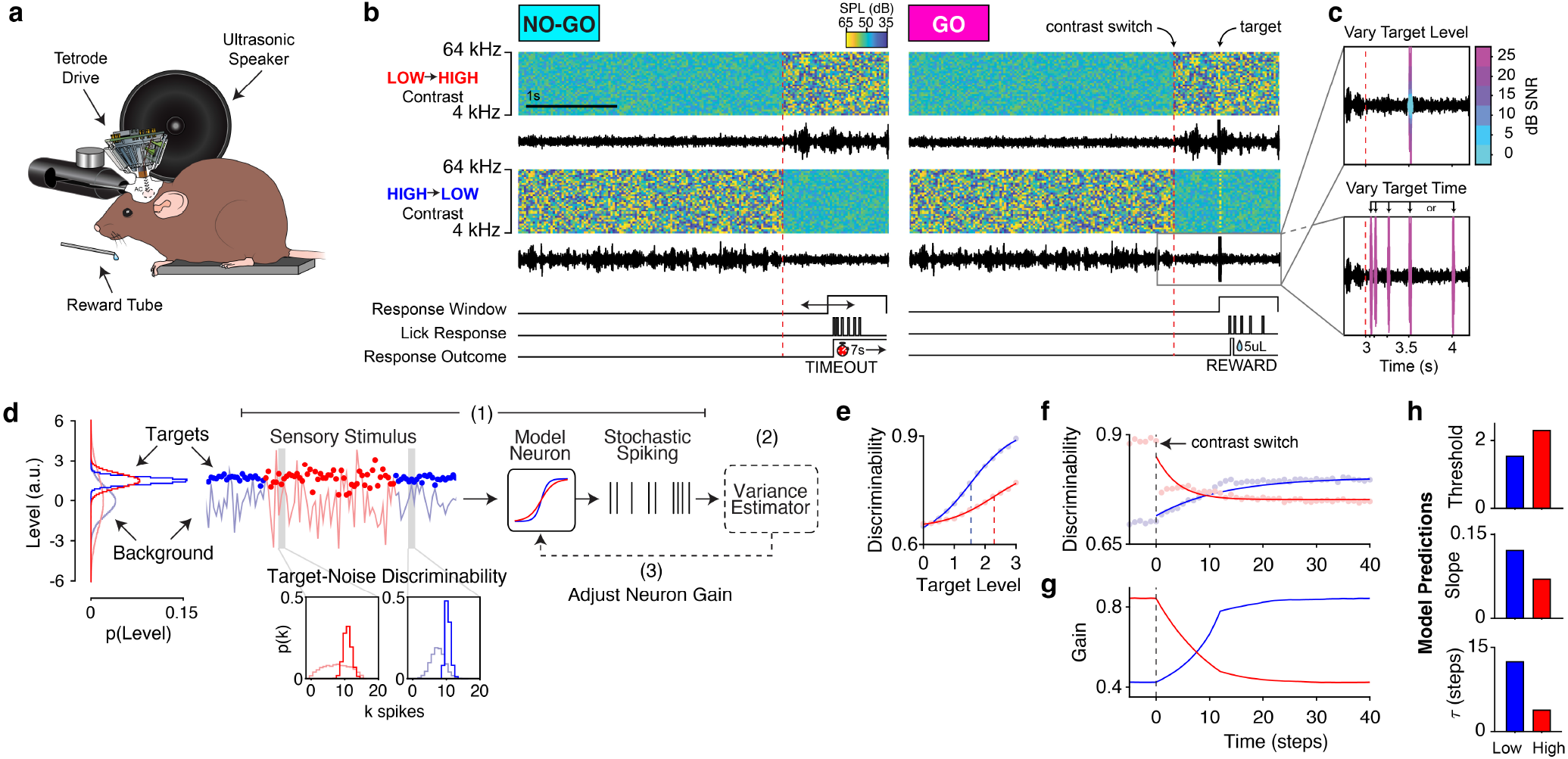
Target in background detection task and normative model predictions. **a,** Experimental setup. **b,** GO/NO-GO task design. Spectrograms are plotted for example NO-GO and GO trials with transitions from low to high contrast (top row) and high to low contrast (bottom row), with waveforms plotted below each spectrogram (color bar indicates the sound level). Below the example trials, the timing of the response window, schematic licks, and responses to licks are plotted. For NO-GO trials, licks in the response window received a timeout. For GO trials, licks in the response window were rewarded with 5µL of water. **c,** Example target parameters. *Top*: varied target levels, with level in dB SNR indicated by the color bar. *Bottom*: varied target times, where each arrow indicates a potential target delay. **d,** Normative model of efficient gain control. Target and background distributions for each contrast are indicated in the left panel. (1) Target stimuli are indicated with circles, while the background stimulus is indicated by a line. The stimulus response in a given time window is transformed by an adapting nonlinearity to generate spikes. (2) The spiking responses are decoded to update an estimate of the stimulus variance. (3) The gain of the nonlinearity is adjusted to optimally predict the variance of the next timestep. *Inset*: Example spike distributions of the model neuron in low and high contrast for targets (dark histograms), and background (light histograms). **e,** Model target-from-background discriminability as a function of contrast and target level. Circles indicate model performance overlaid with logistic function fits (solid lines) and thresholds (dashed lines). **f,** Model discriminability over time in low and high contrast. Circles indicate model performance overlaid with exponential function fits. **g,** Model gain dynamics over time in each contrast. **h,** From top to bottom: model predictions for target detection thresholds, slopes, and adaptation time constants in each contrast. Source data are provided as a Source Data file.

To estimate the optimal time course of contrast gain control and predict its impact on target detection, we developed a normative model of behavioral performance constrained by efficient neuronal coding. In this model, we simulated a neuron designed to encode stimuli with minimal error. To efficiently exploit its finite dynamic range, the model neuron estimated the contrast of the recent stimuli, and adjusted the gain of its nonlinearity to minimize the error in estimated contrast (Figure 1d, panels 1-3; *Methods*)^22, 23^. Adding targets at different levels and times relative to contrast transitions allowed us to probe the sensitivity of the model neuron to targets of varying strength over the time course of adaptation (Supplementary Figure 1c,d). When varying target strength and measuring the ”psychometric” performance of the model (Equation 1, *Methods*), we found decreased detection thresholds and steeper slopes in low contrast relative to high contrast (Figure 1e). When varying target timing, two factors affected target discriminability: 1) The change in the stimulus distribution after the contrast switch; 2) The effect of gain adaptation on responses to the background (Figure 1f,g; Supplementary Figure 1c,d). Recapitulating previous results^14^, the dynamics of the model responses to targets were asymmetric across each contrast (Figure 1f). These dynamics were well characterized by a single effective timescale, which we quantified by fitting an exponential (Equation 2) to each transition. The normative model presented three primary predictions for target detection behavior: When adapted to low contrast, 1) target detection thresholds will be lower and 2) model psychometric functions will have steeper slopes, and, 3) target detectability during the adaptation period will be asymmetric: rapidly decreasing after a switch to high contrast, and slowly increasing after a switch to low contrast (Figure 1h).

### Estimated cortical gain dynamics follow a normative model of gain control

Previous work on contrast gain control used static models, measuring steady-state gain after the neuron fully adapted to the new stimulus^14, 16, 17, 19^, but see^15, 24^. We developed a model to estimate the gain of neurons in auditory cortex over time following a contrast transition. This generalized linear model (GLM) was fit to data recorded from the auditory cortex of a naive mouse (n = 97 neurons) presented with 3 s alternations of low and high contrast DRCs (Figure 2a,b).

**Figure 2.**
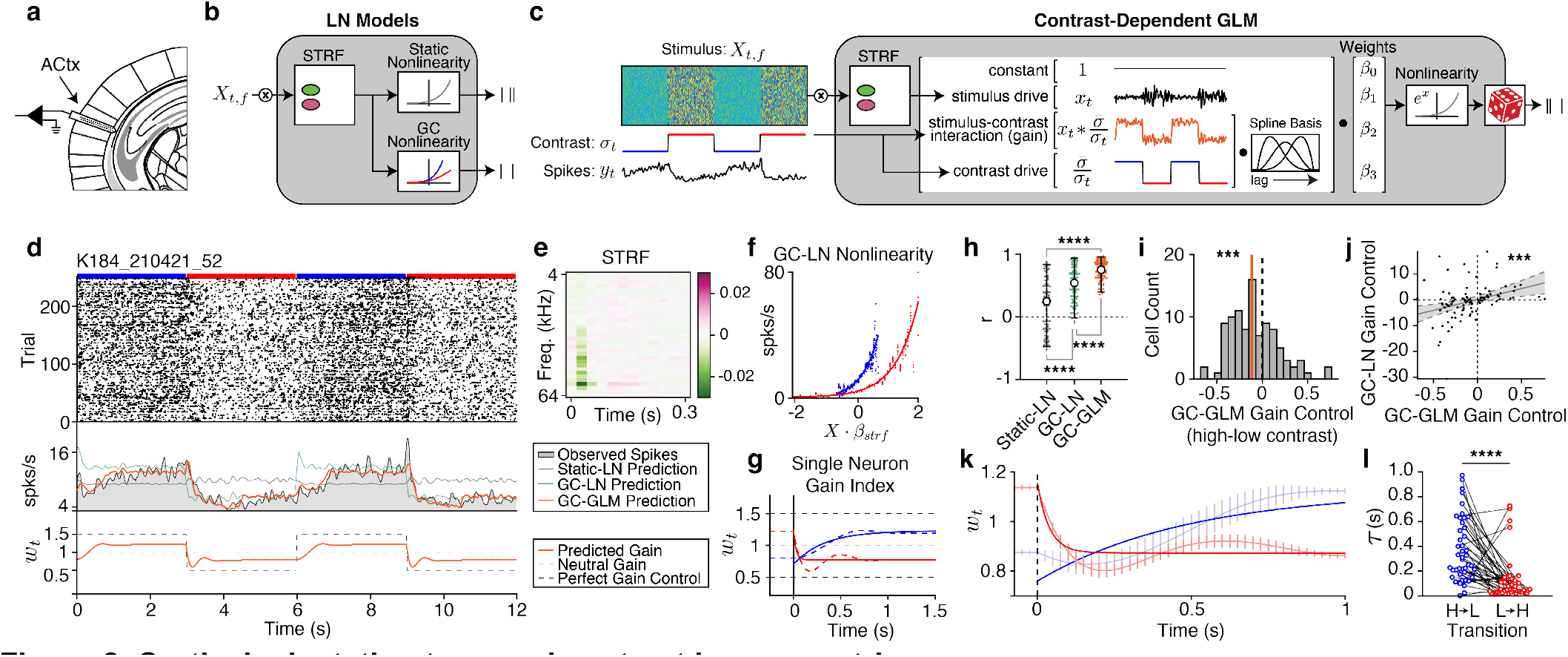
Cortical adaptation to sound contrast is asymmetric. **a,** Schematic^26^ of acute recordings from auditory cortex. **b,** Schematic of the linear-nonlinear (LN) model, with a static (grey) or gain-controlled (blue, red) nonlinearity. **c,** A Poisson GLM for estimating gain dynamics. Note that the model estimates gain as the interaction between the stimulus and contrast. **d**, *Top*: Spike raster for a representative unit. Blue and red horizontal bars indicate low and high contrast periods of the trial, respectively. *Middle*: the spike rate of the neuron is overlaid with the predictions from a static LN model, LN model with gain control, or GLM with gain control. *Bottom*: gain index, *w_t_*, estimated from the GLM parameters (*Methods*). Dashed lines indicate optimal and no gain control (*Methods*). Orange trace indicates the gain dynamics of the neuron. **e,** Spectro-temporal receptive field (STRF) fit to this neuron. Color bar indicates the strength of the filter response. **f**, Nonlinearities fit to the STRF prediction in low and high contrast. **g**, Dashed blue and red lines indicate the gain index of the example cell in low and high contrast. The overlaid solid lines are exponential fits to the data. **h,** Cross-validated Pearson correlation coefficients between the trial averaged model predictions and spike rates for each model (n=97 neurons; colors as in d). Error bars indicate 95^th^ percentiles around the median. Asterisks indicate the results of two-way sign-rank tests. **i,** Distribution of gain control estimated by the GLM. Orange line indicates the median. Asterisks indicate the results of a two-way sign-rank test (*p* = 0.0040). **j,** Gain control estimates for each neuron from the GC-GLM and the GC-LN model (black dots) overlaid with the best linear fit (black line) and 95% confidence interval of the fit (grey area). Asterisks indicate whether the linear model was significantly different from a constant model (*p* = 7.33e-4). **k,** Gain index for all of the neurons with gain control (n=45). Light lines are the average ±SEM, while the dark lines are exponential fits to the average. **l,** Adaptation time constants from gain-controlled neurons after a switch to low (blue dots) and high contrast (red dots). Asterisks indicate the results of a two-way sign-rank test (*p* = 6.16e-6). Source data are provided as a Source Data file.

The inference model is a GLM with dynamic gain control (GC-GLM) that decomposes the relationship between spiking activity (*y_t_*) and the presented sounds into a stimulus component (*x_t_*), contrast component (σ/σ_t_), and an interaction between the stimulus and the contrast (*x_t_*\*σ/σ_t_, where σ is an arbitrary constant, defined as the contrast at which the gain is 1; *Methods*). We calculated a gain index (*w_t_*) from the fitted model parameters (Figure 2b), which quantified whether gain control estimated by the model was optimal given the background contrast levels. For the contrast levels used in this study, *w_t_* = 1.5 indicates an optimal increase in gain during low contrast, *w_t_* = 0.5 indicates an optimal decrease in gain during high contrast, and *w_t_* = 1.0 indicates no gain control (*Methods*). To validate the model and gain index we simulated neurons with defined temporal trajectories of gain control and found that the model accurately recovered the ground-truth simulated gain dynamics (Supplementary Figure 2; *Supplementary Information*). For comparison, we also fit previously described linear-nonlinear (LN) models to each neuron^14, 16, 17, 19^, one with a static output nonlinearity (static- LN), and one with a contrast-dependent, or gain-controlled output nonlinearity (GC-LN, Figure 2c; representative neuron: Figure 2d-g). In this neuron, the fits of both the GC-LN model and GC-GLM exhibited contrast gain control, as characterized by high gain in low contrast and low gain in high contrast (Figure 2f and g, respectively), suggesting that both models capture similar contrast-driven changes in cortical gain.

Qualitatively, GC-GLM outperformed standard LN models, primarily by capturing the adaptation dynamics after the transition (Figure 2d, middle panel), allowing us to analyze the gain control index (*w_t_*) as a function of time (Figure 2d, bottom panel; Figure 2g). To test whether the GC-GLM could better account for the data than standard models, we compared cross-validated correlations of the model predictions with the trial-averaged PSTH for each neuron, finding a significant effect of model type on the correlations (Kruskal-Wallis test: *H*(2) = 93.61, p = 6.70e-21). GC-GLM correlation was significantly higher (Median (*Mdn*) = 0.75, Inter-Quartile Range (*IQR*) = 0.24) than the GC-LN model (*Mdn* = 0.54, *IQR* = 0.49, *p* = 4.41e-6, post-hoc sign-rank test) and the static-LN model (*Mdn* = 0.25, *IQR* = 0.73, *p* = 9.56e-10, post-hoc sign-rank test). Consistent with previous studies^14, 15^, we also found that the GC-LN model outperformed the static-LN model (*p* = 3.50e-6, Figure 2h). To compare the GC-GLM to an existing dynamical model of contrast gain control, we also fit a recent model of dynamic contrast adaptation^19^ to our data (Supplementary Figure 3a). Across the neurons in this dataset, we found that the GC-GLM was a better predictor of neuronal activity than the existing model (Supplementary Figure 3b).

We then quantified whether the GC-GLM reliably detected gain control in the neuronal population. Here, we defined steady-state gain control by calculating the change in *w_t_* between high (*w_H_*) and low (*w_L_*) contrast after the gain has stabilized (1 s after the contrast switch). Based on our definition of *w_t_*, *w_H_* – *w_L_* = –1 if gain control is optimal (*Methods*). Across all neurons, we found significant gain control (*Mdn*: -0.10, *IQR*: 0.35, sign-rank test: *Z* = -2.90, *p* = 0.004; Figure 2i). To further validate the GLM estimates of gain, we compared the GC-GLM gain control indices at steady-state to those of the GC-LN model and found a significant relationship (linear regression: *F*(1,95) = 12.20, *p* = 7.33e-4, *R*^2^ = 0.11; Figure 2j). Together, these results demonstrate that the GC-GLM model better accounts for the neuronal data by incorporating the dynamics of gain control and conclude that this method captures a similar estimate of steady-state gain control when compared to standard models.

Finally, we analyzed the dynamics of gain control by fitting *w_t_* after each contrast switch with an exponential function (Figure 2g). In neurons with gain control (*w_t_* < 0 at steady state), the average time course of *w_t_* was asymmetric across contrast transition types, rapidly decreasing after a switch to high contrast, and slowly increasing after a switch to low contrast (n = 45 neurons; Figure 2k). Within this same population, we quantified the timescale of adaptation to each contrast using the time constant (τ) of each exponential fit, finding significantly longer time constants in low contrast (*Mdn* = 0.29, *IQR* = .39) relative to high contrast (*Mdn* = 0.048, *IQR* = 0.094; sign-rank test: *Z* = 4.52, *p* = 6.16e-6; Figure 2l). This asymmetry in gain adaptation reflected the dynamics of the normative model (Figure 1g), prior electrophysiological studies^14^, and the behavior of previously described optimal variance estimators^25^.

### Mouse behavioral detection is modulated by background contrast

We next tested whether the asymmetry in gain control in cortex was reflected in behavioral sensitivity to targets in background noise. Mice were initially trained in a simple version of the GO/NO-GO task where they were required to lick in response to a target and withhold licks on trials without a target (Figure 1b, 3a). Out of the 25 mice trained, 24 mice learned this task reliably, typically reaching criterion performance of 80% correct within 2-3 weeks in either contrast (Figure 3b). False alarm rates were significantly higher in high contrast than in low contrast (Supplementary Figure 4a), suggesting that detection is more difficult in high contrast, which we discuss next.

**Figure 3.**
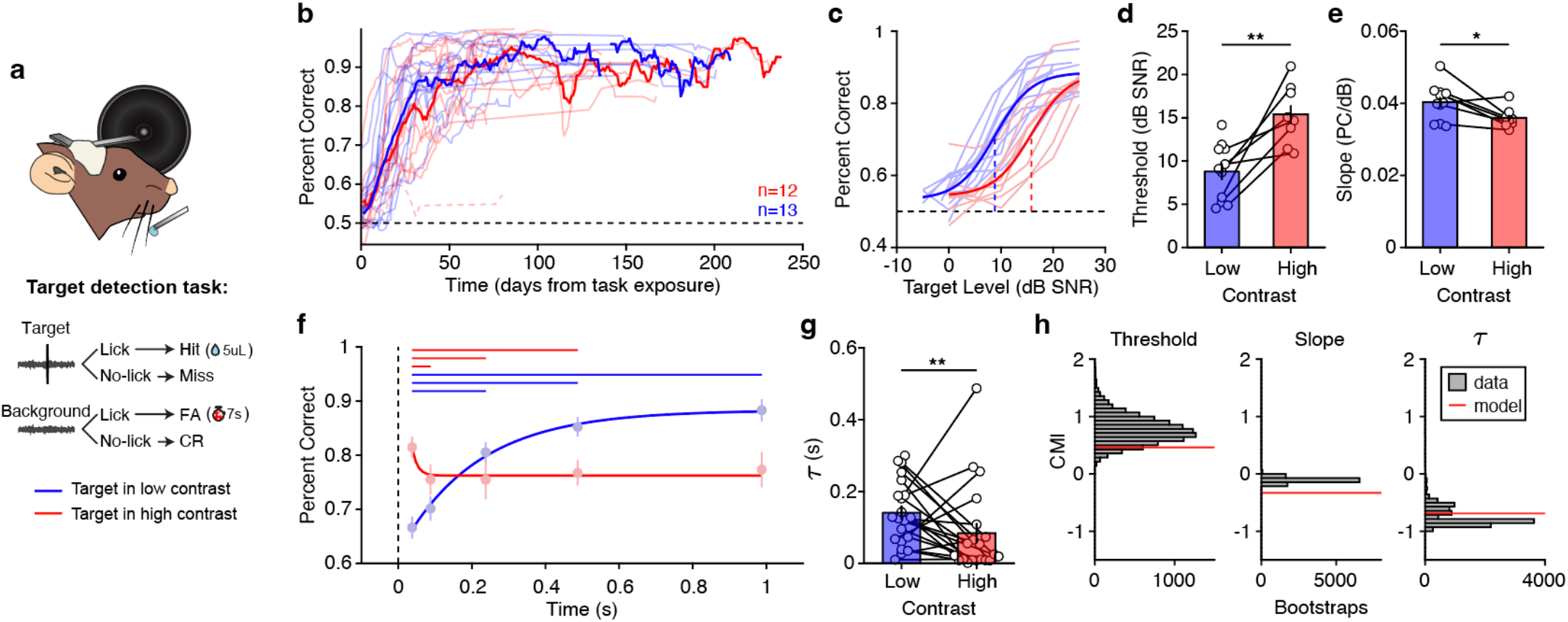
Target detection performance is consistent with the normative model predictions. **a,** Schematic of the behavioral setup and task outcomes. **b,** Performance as a function of task exposure. Individual traces indicate performance of mice first trained to detect targets in low contrast (blue) or high contrast (red). Thick lines are a 7-day running average across mice in each training group. The dashed red trace near 0.5 was the performance of a mouse who failed to learn the task in high contrast and was excluded from further analysis. **c,** Individual psychometric curves in low and high contrast (n=12 mice), overlaid with psychometric fits to the average. Dashed vertical lines indicate detection thresholds. **d,** Detection thresholds from the data plotted in c (n=12 mice; two- way paired t-test: *p* = 0.0057). **e,** Slopes of the psychometric functions plotted in c (n=12 mice; two-way paired t-test: *p* = 0.023). Error bars indicate ±SEM over mice. **f,** Detection performance of threshold level targets presented at different delays from the contrast switch (circles indicate mean ±SEM). Performance is overlaid with exponential fits in each contrast. Horizontal lines above the plot indicate significant sign-rank tests after false-discovery correction. **g,** Adaptation time constants from exponential functions fit to individual mice (n=21). Asterisks indicate the result of a two-way sign-rank test (*p* = 0.0060). Error bars indicate ±SEM over mice. **h,** Comparison of average normative model predictions (red lines) to the data (grey distributions) for psychometric thresholds, slopes and adaptation time. The y- axis is the contrast modulation index (CMI, *Methods*) of the values in each plot. Data distributions were estimated using bootstrapping of the mean (threshold, slope) or median (τ) CMI (10,000 samples with replacement). Source data are provided as a Source Data file.

By varying the level of presented targets, we collected psychometric curves for each mouse in each contrast. To isolate the effect of stimulus contrast on behavioral performance, we included sessions from mice that were exposed to similar target levels in low and high contrast (Figure 3c; see Supplementary Figure 4 and Supplementary Results for results using different target ranges). In this cohort, targets were easier to detect in low contrast: we observed significantly lower detection thresholds in low contrast (Mean (*M*) = 8.79, standard deviation (*std*) *=* 3.13) as compared to high contrast (*M =* 15.39, *std =* 3.27; paired t-test: *t(10)* = - 4.20, *p* = 0.0057, Figure 3d). Furthermore, psychometric slopes were significantly steeper in low contrast (*M* = 0.040, *std =* 0.0048) compared to high contrast (*M =* 0.036, *std =* 0.0026; paired t-test: *t(10)* = 3.037, *p* = 0.023; Figure 3e). Combined, these results demonstrate that targets were easier to detect in low contrast, as predicted from the normative model (Figure 1h).

To assess behavioral adaptation to the background contrast, we presented targets at threshold level at variable delays following the contrast transition. We observed behavioral time courses consistent with the normative model and with gain measured in auditory cortex: after a switch of the stimulus to high contrast, detection rates decreased quickly over time, but after a switch to low contrast detection, rates increased slowly over time (Figure 3f). In high contrast, the first significant drop in performance occurred between the first two time points, while in low contrast the first significant increase in performance occurred between the first and third time points (Figure 3f, Supplementary Table 1). Indeed, fitting exponential functions to performance over time revealed that behavioral adaptation was significantly faster in high contrast (median and interquartile range of time constant: *Mdn* = 0.023, *IQR* = 0.082) compared to low contrast (*Mdn =* 0.13, *IQR* = 0.13; sign-rank test: *Z* = 2.75, *p* = 0.0060; Figure 3g).

To directly compare the predictions of the normative model presented in Figure 1 to behavioral performance, we computed a contrast modulation index to measure the percent change in behavioral and model parameters between high and low contrast (CMI: Equation 3, *Methods*). To assess whether the model prediction was within the range of expected behavioral values, we computed the 95% confidence intervals of the behavioral CMI values using a bootstrap procedure. We found that the CMI values of the normative predictions fell within the range of expected CMI values for behavioral thresholds and adaptation times. As observed in behavior, the model predicted a decrease in slope in high contrast; however, the magnitude of the predicted decrease was larger than the range of observed slope CMI values (Figure 3h). Taken together, the behavioral results were qualitatively consistent with all three predictions from the normative model (Figure 1h): 1) Detection thresholds are lower in low contrast; 2) Psychometric slopes are steeper in low contrast; 3) Performance decreases rapidly in high contrast and increases gradually in low contrast.

### Auditory cortex is necessary for detection in background noise

Whereas gain control is present in many areas along the auditory pathway, it is strongest in auditory cortex^16, 19^. As such, we hypothesized that auditory cortex supports the detection of sounds in the presence of background noise. To test whether auditory cortex is required for task performance, we inactivated auditory cortex using the GABA-A receptor agonist muscimol. We validated that muscimol disrupts cortical coding of target sounds by applying muscimol topically to the cortical surface during passive playback of the behavioral stimuli, finding near complete suppression of target responses (Supplementary Figure 5a-f, *Supplementary Information*).

To test if inactivation of auditory cortex affects behavioral performance, we repeated the same experiments in behaving mice, administering muscimol or saline bilaterally through chronically implanted cannulae (4 mice; Figure 4a). We found a profound decrease in the response rates to targets and background in both contrasts (Figure 4b). We quantified these effects on the psychometric curve using a three-way ANOVA with cortical intervention (muscimol or saline), contrast, and target level as factors. We found significant main effects of cortical intervention (*F*(1,307) = 278.63, *p* = 3.83e-44), contrast (*F*(1,307) = 4.39, *p* = 0.037) and level (*F*(6,307) = 40.90, *p =* 7.54e-36). Post-hoc tests showed that muscimol application significantly decreased hit rates in both contrasts by 31.45% (95% CI: [27.76, 35.14], *p* = 1.060e-10), whereas an increase in background contrast significantly decreased hit rates in both intervention conditions by 3.95% (95% CI: [2.57, 7.64], *p* = 0.036). Furthermore, we observed significant interactions between target level and cortical intervention (*F*(6,307) = 14.11, *p* = 4.47e-14), and between target level and contrast (*F*(6,307) = 2.97, *p* = 7.87e-3), but we did not observe a significant interaction between contrast and cortical intervention, suggesting that muscimol has the same effect in low and high contrast. To quantify the effects of muscimol on behavioral performance, we extracted response rates to the maximum target level, false alarm rates, thresholds, and slopes of psychometric functions fit to each session, and found that muscimol significantly reduced every measure of behavioral performance, with the exception of behavioral threshold (Figure 4c, Supplementary Table 1). From these results, we can conclude that auditory cortex is necessary for detecting targets in background, regardless of background contrast.

**Figure 4.**
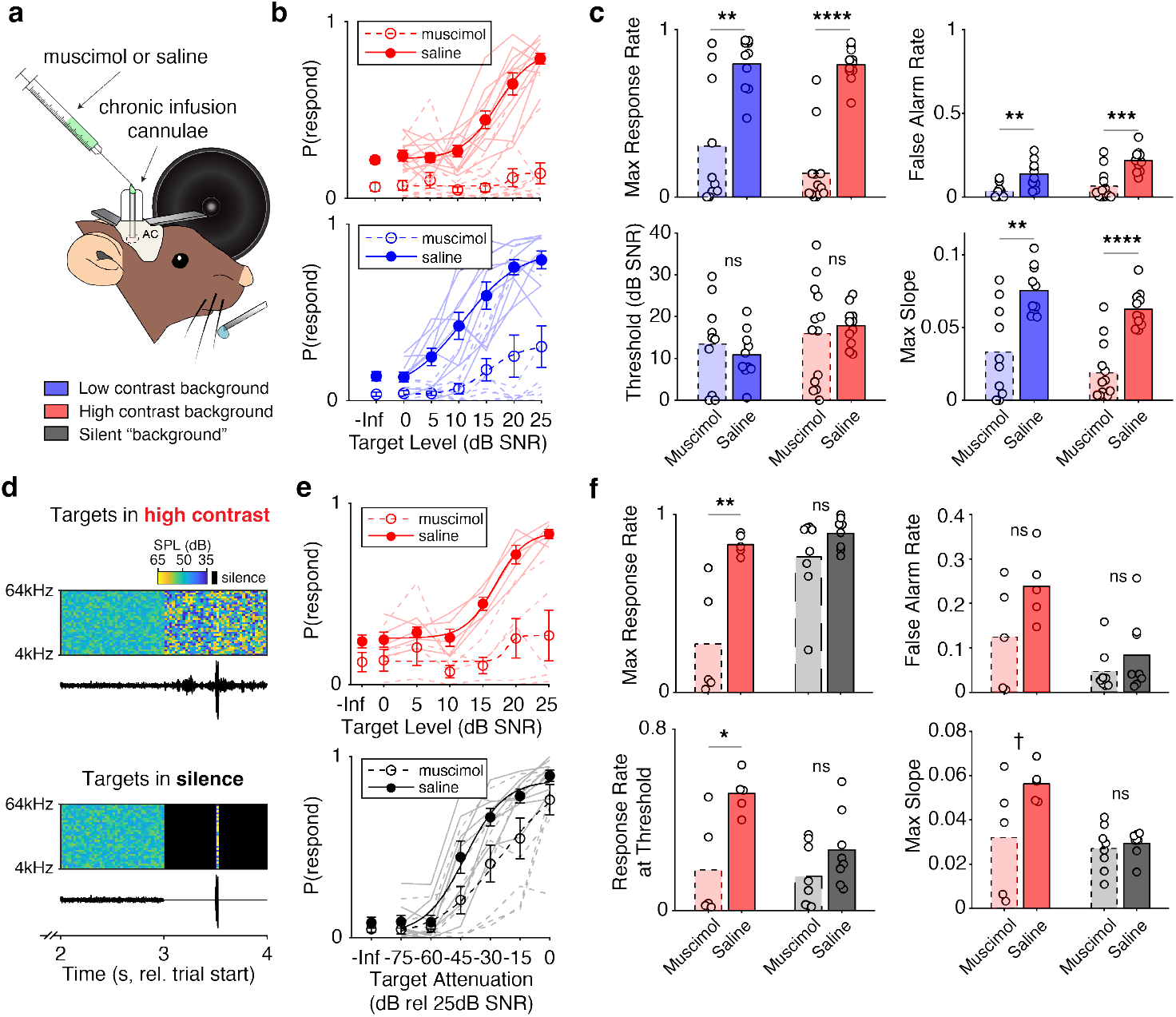
Inactivation of auditory cortex selectively disrupts detection of targets in background sounds. a, Schematic of chronic muscimol and saline application in behaving mice. Legend indicates the three potential background conditions in the task. **b,** Behavioral performance on individual sessions (light traces) as a function of contrast (red, blue) and muscimol or saline application (dashed and solid lines, respectively) for n=44 sessions. -Inf indicates performance on background-only trials. Dots indicate the average performance across sessions ±SEM, overlaid with logistic fits to the data. **c,** Effects of muscimol and contrast on multiple behavioral measures. Bars indicate the mean performance across sessions ±SEM, while dots indicate performance on individual sessions (n=23 muscimol sessions and 21 saline sessions). Asterisks indicate the significance of two-way rank-sum tests (Supplementary Table 1). **d,** Example spectrograms and waveforms with a target presented in high contrast (top) and the same stimulus when the target was presented in silence (bottom). Color bar indicates the sound level, with black indicating silence. **e,** Behavioral performance in high contrast (top) and in silence (bottom) for n=26 sessions. Error bars are ±SEM across sessions. Formatting as in b. **f,** The effect of muscimol on multiple behavioral measures when targets were presented in high contrast (red bars; n=10 sessions) or in silence (black bars; n=16 sessions). Error bars are ±SEM across sessions. Asterisks indicate the significance of two-way rank-sum tests. Formatting as in c. In all plots: ns, not significant; ^†^p<0.1; *p<0.05; **p<0.01; ***p<0.001; ****p<0.0001. Detailed statistical results for c and f are in Supplementary Table 1. Source data are provided as a Source Data file.

A potential alternative effect of muscimol is a general loss of function that is not specific to hearing target sounds. To control for this, we devised an alternative to the detection in background task where mice detected targets in silence (Figure 4d). To ensure equivalency between the two tasks, we took the highest- level target trials in the target-in-background task (25dB SNR in high contrast) and removed the background noise during the target detection period (Figure 4e, bottom). In this new task, mice heard the exact same targets as in the target-in-background task except that the background noise pre- and post-target presentation was removed. This allowed us to test whether auditory cortex is specifically required for target detection in the presence of a background (*Methods*).

To assess behavioral performance in this new task, we modulated detection difficulty by attenuating the level of each target. As observed previously, inactivation of auditory cortex impaired detection in high contrast (Figure 4e, top). However, cortical inactivation had little effect on behavioral performance in silence (Figure 4e, bottom). We quantified these effects on behavior using a three-way ANOVA with cortical intervention (muscimol or saline), task (detection in background or silence), and target level as factors (n = 26 sessions from 2 mice). We found significant main effects of intervention (*F*(1,181) = 62.83, *p* = 3.62e-13), task (*F*(1,181) = 6.82, *p* = 9.86e-3), and level (*F*(6,181) = 46.16, *p* = 1.69e-32). Muscimol significantly reduced hit rates by 20.2% (95% CI: [15.19, 25.17], *p* = 1.060e-10, post-hoc). Hit rates for targets presented in silence were significantly elevated by 6.65% relative to targets presented in background (95% CI: [1.65, 11.64], *p* = 0.0090). Furthermore, we found significant interactions between cortical intervention and task type (*F*(1,181) = 6.36, *p* = 0.013), intervention and level (*F*(6,181) = 3.47, *p* = 2.98e-3), and level and task type (*F*(6,181) = 8.47, *p* = 5.43e-8). As before, we parameterized behavioral performance by fitting the performance on each session with a psychometric curve, and we extracted the response rates to the maximum target level, false alarm rates, response rates at threshold level, and slopes of psychometric functions. During the target-in-background task, we found significant effects of muscimol on the response rates at maximum level and threshold, a moderate effect on psychometric slope, and no effect on false alarm rate. However, muscimol application had no significant effect on any of these measures in the target-in-silence task (Figure 4f, Supplementary Table 1). Taken together, these results show that while both cortical inactivation and the presence or absence of background noise affected behavioral performance, these effects interacted: muscimol had a larger effect on performance when background noise was present.

Combined, our findings demonstrate that the auditory cortex is specifically required for detection in the presence of background noise, but not in silence. Our next goal was to test whether neuronal activity in AC is predictive of behavioral performance.

### Cortical codes predict individual behavioral performance

To better understand how representations in the auditory cortex could give rise to behavior, we chronically recorded from populations of neurons in auditory cortex of 12 mice while they performed the psychometric task (Figure 5a, Supplementary Figure 6). When using microdrives with drivable tetrodes, we lowered the tetrodes a small distance at the end of each session to record from new populations of cells. In the 242 sessions analyzed, we recorded from 18±15 neurons simultaneously (mean±standard deviation), with a maximum of 73 and minimum of 3 neurons simultaneously recorded. For the following analyses, we only included neurons with spike rates greater than 1 Hz, realistic spike waveforms (*Methods*), and with significant responses to targets (AUC value significantly greater than 0.5 at two or more levels, Figure 5b, inset; *Methods*). Following these selection criteria, 12±9 neurons were included in analysis for each session.

**Figure 5.**
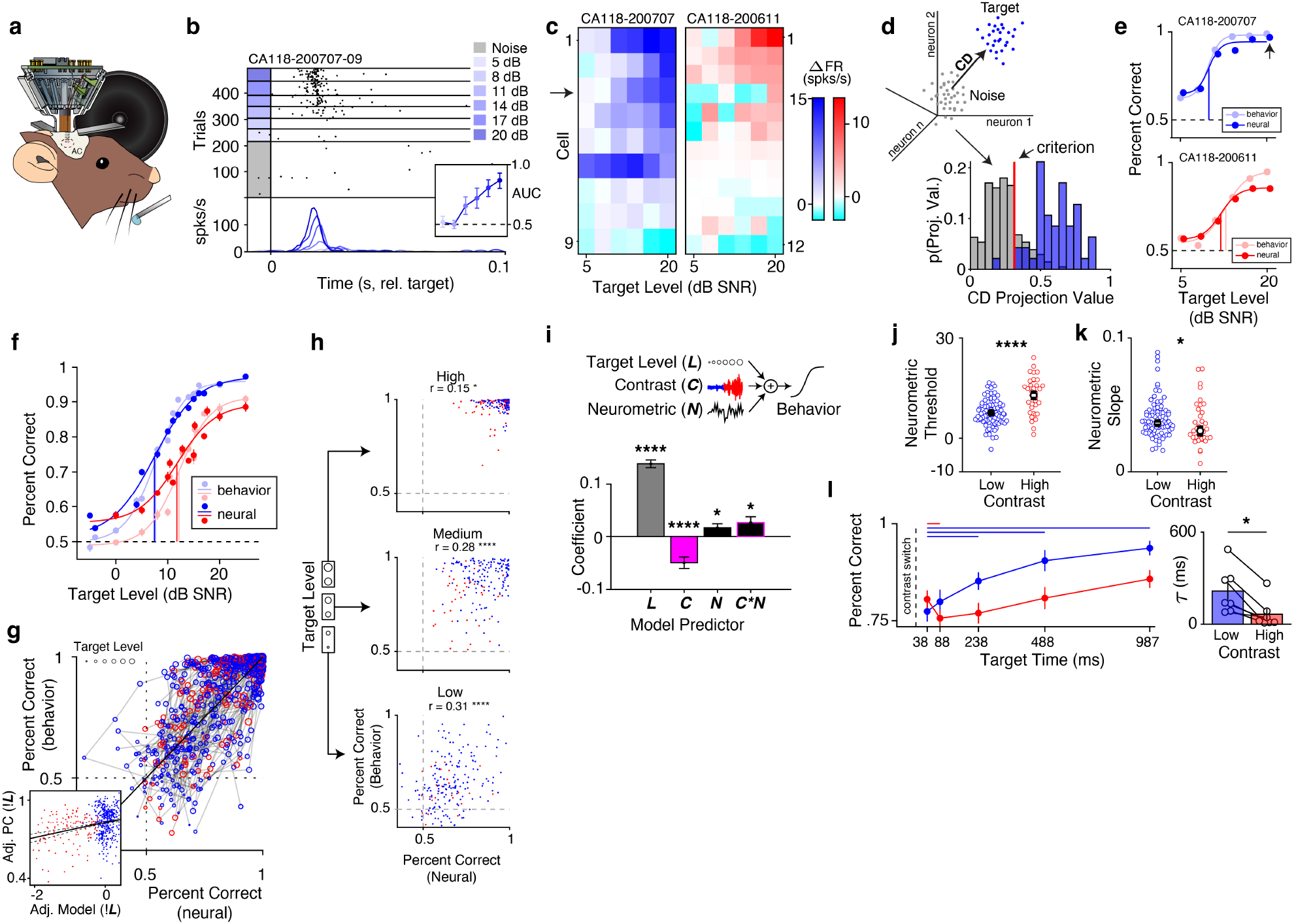
Cortical responses to targets predict behavioral performance and exhibit contrast adaptation. **a,** Schematic of chronic recordings during behavior. **b,** Spike raster from a neuron recorded during the task with average PSTHs. *Inset*: area under the ROC curve (AUC) for each level ±bootstrapped 95% CI. **c,** Neurograms of populations of simultaneously recorded neurons in low (left) and high contrast (right) sessions. Color bars indicate the difference in firing rate between the noise-only condition and each target level. The arrow indicates the cell in b. **d,** *Upper left*: schematic for estimating the population coding direction (CD). *Lower right*: probability distributions of CD projections for noise-only trials (grey) and the highest-level target trials (blue). The vertical red line indicates the criterion for the decoder. **e,** Neurometric performance and behavioral performance in representative low contrast (top) and high contrast (bottom) sessions. Dark dots are decoder performance, while light dots indicate behavioral performance, overlaid by logistic function fits. The arrow indicates the decoder performance for the distributions in d. **f,** Average psychometric and neurometric performance ±SEM over sessions. Formatting as in e. **g,** Behavioral performance as a function of neurometric performance for n=102 sessions. Each point is performance at a single target level in a single session. Point color indicates contrast, while point size indicates target level. Dots joined by grey lines were collected in the same session. *Inset*: Same data as in g, with the effect of target level regressed out (***!L***). The black line indicates the model fit and 95% CI. **h,** Behavioral performance as a function of neural performance split by target levels. Text indicates the correlation coefficient and its significance value. **i,** *Top:* the linear model used to predict behavior. *Bottom:* model coefficients ±standard error for target level (***L***), contrast (***C***), neurometric performance (***N***) terms, and the interaction between the latter two terms (***C*N***). Asterisks indicate significant predictors (n=102 sessions). **j,** Neurometric thresholds ±SEM for sessions in low (n=82) and high contrast (n=35). Asterisks indicate significant two-way rank-sum test (*p* = 1.34e-6). **k,** Same as j, but for neurometric slopes (two-way rank-sum test: *p* = 0.029). **l,** *Left*: Neurometric performance ±SEM as a function of target delay over sessions (n=87). Formatting and statistical tests are the same as Figure 3f. *Right*: Adaptation time constants ±SEM fit to the neurometric response from n=12 mice. Asterisk indicates a significant two-way sign-rank test (*p* = 0.016). Source data are provided as a Source Data file.

To quantify the representations of targets and background in the neuronal population (example responses in Figure 5b,c), we adapted a population vector approach^27^ to generate a target-from-noise discriminability metric using population activity (*Methods*). This method allowed us to project trial distributions in n-dimensional neuronal space along a single dimension that separated target and background trials (Figure 5d, left panel). We then estimated the criterion projection value that best predicted whether each trial contained a target or just background^28^ (Figure 5d, right).

This population decoding method allowed us to estimate neurometric functions to directly compare to psychometric functions for each mouse (Figure 5e). On average, neurometric and psychometric functions were qualitatively similar (Figure 5f) and highly correlated across sessions (Figure 5g). To assess the relationship between neurometric and behavioral performance, we first considered the potential variables that were likely involved to modulate behavior, namely: the level of the presented targets, the contrast of the background, and neuronal variability that could fluctuate from session to session. To isolate the effects of each of these variables, we used a linear model to predict behavioral performance on each session using target level, contrast, and the neurometric performance as predictors (*F*(4,794) = 593.0, *p* = 3.15e-202, *R*^2^ = 0.69, Figure 5i, top).

As expected, we found that basic stimulus features (target level and background contrast) were strong predictors of behavioral performance, such that performance was higher for increased target levels (*b* = 0.14, 95% CI [0.12, 0.15], *p* = 4.15e-65), while performance was lower for increased contrast (*b* = -0.050, 95% CI [-0.071, -0.028], *p* = 5.67e-6). These results confirm our previous results on the behavioral impacts of stimulus contrast (Figure 3). Crucially, we also observed that population activity was predictive of behavioral performance at a session level, such that an increase in neurometric performance was accompanied by an increase in behavioral performance (*b* = 0.016, 95% CI [0.0017, -0.031], *p* = 0.029). Finally, we found that the relationship between behavioral and neuronal performance was mediated by stimulus contrast (*b* = 0.026, 95% CI [0.0042, 0.048], *p* = 0.020; Figure 5i, bottom). Taken together, these results suggest that session-to-session fluctuations in cortical population activity resulted in corresponding fluctuations in behavior, and that this relationship was affected by stimulus contrast.

To control for the effect of target level on the relationship between behavioral and neuronal performance, we first visualized the data by regressing out the effect of level on the model terms and behavioral performance: this visualization demonstrates the positive relationship between the two terms (Figure 5g, inset), as captured by the significant neurometric coefficient in Figure 5i. To further control for the effect of level, we split the data into low, medium and high target levels (Figure 5h). In all level-splits, the relationship between neurometric and behavioral performance was significant (Supplementary Table 1). These findings suggest that regardless of the target level, neuronal performance predicted behavioral performance.

Next, we assessed whether neurometric performance was affected by contrast similarly to behavioral performance. To isolate the effect of contrast on neurometric curves and best compare to psychometric curves presented in Figure 3, we selected only recording sessions with matched target ranges (n = 117) and fit each neurometric curve with a logistic function. We found that neurometric function thresholds increased in high contrast (*Mdn[IQR]* of low contrast sessions (n=82): 7.19[4.83], high contrast sessions (n = 35): 13.47[8.88]; rank-sum: *Z* = 4.69, *p* = 1.34e-6, Figure 5j), and that neurometric slopes decreased in high contrast (low contrast: 0.036[0.014], high contrast: 0.030[0.021]; *Z* = 1.88, *p* = 0.029, Figure 5k). These findings corroborated the observed changes in psychometric functions (Figure 3) and the predictions of the normative model (Figure 1).

Combined, these results demonstrate that parameters of neurometric and psychometric functions are affected by contrast as predicted by a normative model of gain control. We also find that individual variation in behavioral performance is predicted by population activity in auditory cortex, independently of the effect of contrast and target level, which further supports the role of auditory cortex in the detection of signals in noise.

### Dynamics of cortical target detection during adaptation

We next measured how cortical discriminability evolved as a function of time and contrast in sessions where mice were presented with targets at threshold level at different offsets relative to the contrast switch. In line with our behavioral results (Figure 3f), we found that in high contrast the first significant drop in cortical discriminability occurred between the first two target times, while in low contrast the first significant drop occurred between the first and third target times (n = 43 recording sessions; Supplementary Table 1; Figure 5l, left). To quantify the speed of neuronal adaptation, we fit the average neuronal discrimination time course for each mouse with an exponential function (n = 12 mice). Consistent with the normative model (Figure 1f- h), and with gain control dynamics estimated from cortical activity (Figure 2k,l) and behavior (Figure 3f,g), we found asymmetric adaptation in the neuronal responses, with larger adaptation time constants in low contrast (*Mdn* = 0.14, *IQR* = 0.21) relative to high contrast (*Mdn* = 0.033, *IQR =* 0.16; sign-rank test: *p* = 0.016; Figure 5l, right).

### Cortical gain predicts behavioral performance

Our results so far demonstrate that population codes in auditory cortex are not only shaped by stimulus features, but also predict variation in behavioral performance. To assess the role of cortical gain in behavior, we leveraged the design of the background sounds to estimate spectrotemporal receptive fields (STRFs) and nonlinearities of neurons recorded during task performance. For each neuron, we fit a model with a static nonlinearity (static-LN) or a model with gain control (GC-LN; Figure 6a-d). We then pooled the neurons recorded across sessions including only neurons with strong stimulus responses in both contrasts (*Methods*). To ensure that observed changes in gain were not due to changes in STRF shape during the task, we first compared STRFs estimated from each task epoch (ie. low and high contrast) and found that STRF properties were stable across the trial (Supplementary Figure 7, *Supplemental Methods*).

**Figure 6.**
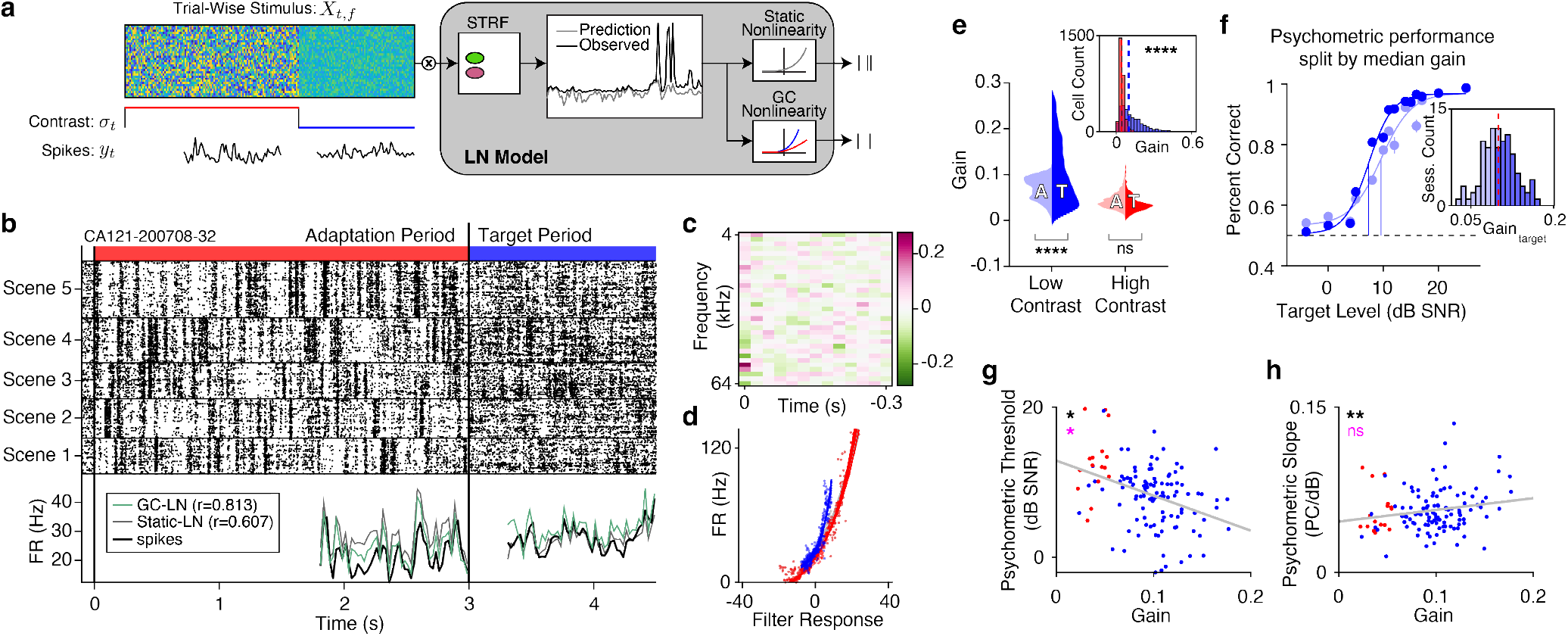
Cortical gain predicts session-to-session variability in behavioral performance. **a,** Schematic of the LN model fit to neuronal responses during the behavioral task. **b,** Spike raster of a representative unit recorded during the behavioral task, sorted by the background scene presented in each trial. Below is the average spike rate (black trace) overlaid with the LN model fit with a static (grey) or gain-controlled nonlinearity (green). **c,** STRF for the representative cell. **d,** The fitted nonlinearities in low and high contrast periods of the trials. **e,** Gain distributions across all recorded neurons as a function of contrast and trial period (A: adaptation, T: target). Gain was significantly larger in the target period during low contrast (two-way ANOVA: *p* = 3.77e-9) but not in high contrast (*p* = 0.18) Inset: the average gain in low and high contrast for all cells, dashed lines indicate the median of each distribution. Gain was higher in low contrast (two-way ranksum: *p* = 1.60e-91. **f,** Psychometric curves split by the median gain (n=107 sessions). Light colored dots indicate performance across sessions with low gain, dark colored dots indicate performance on sessions with high gain. Error bars indicate ±SEM. *Inset*: the distribution of gain values on the same sessions. The dashed red line indicates the median used to split the data. **g,** Relationship between session-to-session changes in gain and behavioral thresholds (n=124 sessions). Each dot is a session, with the color indicating the contrast in which targets were presented. The grey line is the linear best fit. Black asterisks indicate whether gain is a significant predictor of the psychometric threshold, while magenta asterisks indicate whether contrast was a significant predictor of behavioral thresholds (Supplementary Table 1). **h,** Same formatting as g, but plotting the relationship between gain and psychometric slope (n=124 sessions). Source data are provided as a Source Data file.

First, we compared the cross-validated performance of the static-LN model versus the GC-LN model. We found higher correlations using the GC-LN model (*Mdn* = 0.82, *IQR* = 0.17) relative to the static-LN model (*Mdn* = 0.67, *IQR* = 0.12; n = 2,792 neurons; sign-rank test: *Z* = -36.74, *p* = 1.88e-295; Supplementary Figure 8a). We also found significantly higher gain in low contrast (*Mdn* = 0.10, *IQR* = 0.13) than in high contrast (*Mdn* = 0.041, *IQR* = 0.023; sign-rank test: *Z* = 37.92, *p* = 1.070e-314; Figure 6e, inset). These results demonstrate that LN models can more accurately predict cortical activity when incorporating contrast gain control, and confirm previous reports of robust gain control in mouse auditory cortex^17–19^.

Based on our previous results, we expected that the strength of gain control in auditory cortex would predict target detectability. When fitting the GC-LN model, we separately estimated neuronal gain during the adaptation period of the trial and the target period of the trial (defined as the time periods before and after the contrast switch, respectively; Figure 6b). To quantify the effects of contrast and trial period on gain, we performed a two-way ANOVA, with gain as the dependent variable, and contrast, trial period, and their interaction as factors (n = 2,262 neurons, after excluding outliers, see *Methods*). As expected, we found a significant main effect of contrast (*F*(1,4523) = 431.03, *p* = 1.60e-91). Furthermore, there was a significant main effect of trial period (*F*(1,4523) = 35.79, *p* = 2.36e-9) and a significant interaction between contrast and trial period (*F*(1,4523) = 77.91, *p* = 1.51e-18). Post-hoc tests revealed that, in low contrast, gain during the target period significantly increased (0.032, 95% CI: [0.024, 0.040], *p* = 3.77e-9), but did not significantly change in high contrast (0.0062, 95% CI: [-0.017, 0.014], *p* = 0.18; Figure 6e). These findings indicate that neuronal gain is not only sensitive to stimulus contrast, but also increases during the target period of the trial, specifically in low contrast.

To visualize the gross relationship between gain and behavioral performance, we first averaged the gain of stimulus-responsive neurons during the target period of the trial in each session (n = 168 sessions across 13 mice). We then selected only low contrast sessions and split the data by the median gain in the target period, computing the average psychometric curves for sessions in the bottom versus the top 50^th^ percentile (Figure 6f, inset). We observed that sessions with high gain had steeper slopes and lower thresholds than sessions with low gain (Figure 6f). To quantify the relationship between gain and task performance, we fit a mixed-effects model using contrast and gain during the target period as fixed effects, mouse identity as a random effect, and either psychometric slopes or thresholds as the dependent variable. This approach allowed us to separate the neuronal and behavioral impact of contrast gain control from the effect of session-to-session fluctuations in gain. We tested whether gain and contrast were significant predictors of behavioral performance by comparing the full model to null models excluding either gain or contrast. We found that the model including gain was a better predictor of behavioral threshold than was the null model (Likelihood Ratio Test: χ^2^(1) = 5.82, *p* = 0.016), indicating that thresholds decreased by 3.046 dB SNR ±1.24 (SEM) for every 10% increase in gain. Using a similar procedure, we found that contrast was also a significant predictor of behavioral threshold (Likelihood Ratio Test: χ^2^(1) = 5.84, *p* = 0.038), with the step from high to low contrast inducing a decrease in behavioral thresholds of 3.27 dB SNR ±1.33 (Figure 6g).

We applied the same analysis to test the effects of contrast and gain on psychometric slope (Figure 6f), again finding that gain significantly predicted psychometric slopes (Likelihood Ratio Test: χ^2^(1) = 6.96, *p* = 0.0083), such that the psychometric slope increased by 0.16 dB/PC ±0.060 for every 100% increase in gain. However, contrast did not significantly improve the fit of this model (Likelihood Ratio Test: χ^2^(1) = 2.28, *p* = 0.13; Figure 6h). This result is not entirely unexpected, given that we observed no effect of contrast on psychometric slopes when comparing across sessions with different target distributions (Supplementary Figure 2b), which is true of the sessions used in this analysis.

Our findings suggest that the relationship between gain and behavioral performance is shaped by two sources: contrast-induced gain control and fluctuations in gain from session to session. To further disentangle the relationship between these two sources of behavioral modulation, we repeated the mixed effects models, this time using gain during the adaptation period as the predictor of interest. We hypothesized that gain in this period should not be predictive of behavioral performance, as there were no targets presented during this portion of the trial. We found that this was the case; we did not observe any predictive relationship between gain estimated in this period and behavioral performance (Supplementary Figure 8b-d; Supplementary Table 1). In summary, we found that cortical gain was modulated by both stimulus contrast and trial period, increasing when contrast is low and when mice were detecting targets. Furthermore, we found that behavioral performance was predicted by both the stimulus contrast and by session-to-session changes in cortical gain during target detection.

## Discussion

Auditory environments exhibit complex statistical properties that change over time. Changes in the dynamic range, or contrast, of acoustic inputs poses a challenge to the auditory system, which is composed of neurons with limited dynamic range. The efficient coding hypothesis predicts that as stimulus contrast changes, neurons should adjust their gain in order to match their limited dynamic range to that of the stimulus distribution^1^. Multiple studies have demonstrated that neurons throughout the auditory pathway exhibit such contrast gain control^14, 16, 19^. Additionally, contrast gain control is theoretically beneficial for separating signals from background noise^16, 20^, implicating that this phenomenon should underlie perception in noisy environments. Whereas recent work has demonstrated a link between contrast gain control and changes in behavioral discrimination in humans^19, 21^, it is unclear how cortical gain directly relates to behavior, as neuronal responses and behavior were not observed simultaneously. Finally, recent work has highlighted the dynamic nature of efficient neuronal codes^22, 23^, and it is unclear whether the observed neuronal dynamics reflect what is predicted to be theoretically efficient, and how the observed dynamics affect behavior.

Our study addressed these open questions by developing a normative framework to predict how efficient gain adaptation should affect behavior, and then testing those predictions using a combination of behavior, chronic recordings, and cortical manipulations. First, we developed a normative model based on efficient coding^22, 23^, which predicted that: 1) Detection thresholds of targets should be lower in low contrast than in high contrast; 2) Sensitivity to changes in target level should be greater in low contrast relative to high contrast; and 3) Detection should change asymmetrically over time: increasing slowly after a switch to low contrast, but decreasing rapidly after a switch to high contrast (Figure 1). Then, we derived a form of Poisson GLM to confirm that gain control dynamics in auditory cortex are indeed asymmetric (Figure 2). To behaviorally test the predictions of the normative model and GLM, we trained mice to detect a target embedded in dynamic random chords while manipulating the contrast of the background between high and low contrast. As predicted by the normative model, mice had lower detection thresholds and were more sensitive to changes in target level in low contrast. Behavioral adaptation was also asymmetric, decreasing rapidly after a switch to high contrast, and increasing slowly after a switch to low contrast, in agreement with our model (Figure 3). Furthermore, we found that AC is necessary for this detection-in-background task (Figure 4). When recording in AC, we found that the variability in neuronal population responses significantly predicted session-by-session variability in behavior above and beyond variability induced by stimulus features, and we demonstrated that target discriminability adapted asymmetrically, as predicted (Figure 5). Finally, we found that the amount of cortical gain also predicted behavioral performance on a session-to-session basis, independently of the effect of contrast (Figure 6). Taken together, these results support our hypothesis that the dynamics of gain control support efficient coding in auditory cortex and predict the perception of targets embedded in background sounds.

### The role of cortex in behavior

The role of auditory cortex in behavior has been subject of debate. A number of prior studies found that auditory cortex was not required for relatively simple behavioral tasks such as frequency discrimination or detection^29, 30^. Rather, many studies found that auditory cortex is primarily involved in more complex behaviors, such those requiring temporal expectation^31^, localization^32^, or discrimination of more complex sounds^33–35^. Consistent with previous findings^36^, we found that AC inactivation selectively impaired detection of targets in a noisy background, but did not impair detection of targets in silence (Figure 4). Furthermore, neuronal activity in AC predicted variability in behavioral performance (Figures 5, 6). This set of results establishes that AC is necessary for the detection of targets in background noise and supports the more general notion that AC is required for more difficult auditory tasks.

While the previous work demonstrates the necessity of auditory cortex in behavioral performance, the brain areas and mechanisms supporting the transformation from stimulus to decision are an active field of study^37, 38^. By recording neuronal activity during the task, we were able to leverage behavioral variability to show that task performance covaried with representations of targets within small neuronal populations (Figure 5), and with cortical gain (Figure 6). There is a large body of literature relating cortical codes to behavioral variability: early studies in the visual system suggested that information from relatively small numbers of neurons was sufficient to match or outperform animal behavior in psychophysical tasks^39–41^ and that behavioral choice can be predicted from activity in sensory areas^28, 41^. These accounts suggested that variability in bottom-up sensory encoding drives the variability in behavioral output. However, more recent work suggests that variability in sensory areas is driven by top-down influences^42–45^, which are modulated by attention and learning^46–49^. A recent study imaging tens of thousands of neurons in the visual cortex supports this notion, finding that cortical representations had higher acuity than behaving mice, yet did not correlate with behavioral performance, suggesting that perceptual discrimination depends on post-sensory brain regions^50^.

Our results suggest that bottom-up adaptation to stimulus statistics shapes behavioral output: We observed asymmetric time courses of target discrimination following a change in contrast (Figure 3). The asymmetric adaptation in behavioral performance was consistent with predictions derived from a normative account of contrast gain control (Figure 1), resembled contrast gain adaptation in auditory cortex in the absence of behavior (Figure 2), and was evident in patterns of target-driven activity in auditory cortex during task performance (Figure 5). Indeed, there have been other studies demonstrating that individual differences in sensory-guided behaviors are reflected in cortical activity^51, 52^, are bidirectionally modulated by cortical manipulation^53, 54^, and can be predicted from tuning properties in auditory cortex^55, 56^. While our results cannot rule out top-down input as the causal driver of sensory decisions, they do support the notion that the sensory information upon which decisions are made is shaped by neuronal adaptation, which thereby affects behavioral outcomes.

### Roles of gain in the auditory system

Neurons throughout the auditory system adapt to the statistics of the acoustic environment, including the frequency of stimuli over time^57, 58^, more complex sound patterns^24, 59^, and task-related or rewarded stimuli^60–65^. In this study, we focused on contrast gain control as a fundamental statistical adaptation that relates to efficient coding^14, 17–19^. Previous work identified that responses to targets presented after opposing changes in contrast were asymmetric^14^, a result consistent with optimal estimation of stimulus variance^25^. We observed similar asymmetric dynamics in a normative model of contrast gain control (Figure 1), which suggests that previous observations of asymmetric adaptation are a result of efficient encoding of the background stimulus.

Inspired by this and other previous work^60^, we intentionally designed our stimuli using unbiased white-noise backgrounds, which allowed us to fit encoding models to our data. In this study, we developed an application of Poisson GLM that allowed us to quantify gain dynamics directly from responses to continuous DRC stimuli. This approach allowed us to verify that gain adaptation in auditory cortex is asymmetric (Figure 2), and we found that this model better predicted neuronal activity when compared to previous models (Figure 2, Supplementary Figure 3). While we focused on the multiplicative influence of contrast in this study, the GLM presented here could in theory be applied to any other time-varying signal that modulates neuronal gain, such as movement^66, 67^, arousal^68, 69^, or experimental interventions such as optogenetics^70–73^

Additionally, previous studies found that contrast gain control was predictive of behavioral performance^10, 19^. Here, we extended these findings by examining the dynamics of this process and found that behavioral detection of targets also adapted asymmetrically (Figure 3). In addition to confirming previously reported effects of contrast on psychometric curves, these results suggest that the dynamics of contrast gain control influenced task performance. Pursuing this further, we estimated cortical gain as mice performed the task, and discovered a predictive relationship between stimulus contrast, gain in auditory cortex, and behavioral performance (Figure 6). These results suggest two sources of cortical gain modulation: 1) Bottom-up adaptation to stimulus contrast (ie. contrast gain control), and 2) session-to-session modulation of gain. Previous studies have demonstrated this latter phenomenon, suggesting that top-down gain modulation underlies attention^42, 43, 74^ and the maintenance of optimal behavioral states^68, 69^. Our results suggest that both contrast gain control and session-to-session fluctuations in gain modulate behavior, providing a starting point for dissecting the neuronal mechanisms underlying these two forms of gain modulation.

### Cellular mechanisms of gain control

While this work and other studies have established contrast gain control as a fundamental property of the auditory system, the neuronal mechanisms driving gain adaptation at a cellular level remain unclear. Previous work theorized that cortical feedback was responsible for contrast gain control in early areas^16, 20^, but recent experiments disproved this hypothesis^19^. In the current study, we have likely recorded from a mixed population of excitatory and inhibitory neurons, the latter of which exhibit specific roles in adaptation^75, 76^. While specific inhibitory neuronal subtypes facilitate divisive or subtractive control of excitatory responses in visual^70, 71^ and auditory cortex^72, 73^, the role of these interneurons in contrast gain control has been inconclusive^18^. Additionally, our results highlight the distinction between stimulus driven gain control and stimulus-invariant gain control which covaried with behavior from session to session (Figure 6). Whether these two forms of gain control share common neuronal substrates is unclear. By combining cell-specific optogenetic methods with behavioral tasks, future studies may explore and test the causal role of local circuits and top-down modulation in gain control and behavior.

### The missing link between efficient coding and behavior

Combined, our results develop a framework and provide support for the role of contrast gain control in behavior. The efficient coding hypothesis has emerged as one of the leading principles in computational neuroscience that has shaped our understanding of neuronal coding, architecture, and evolution^1, 77–80^, and prior research found that human behavior follows principles of efficiency^19, 81^. Here, we focused on a well- studied form of efficient coding, contrast gain control, and developed a framework to link the dynamics of efficient neuronal coding with behavioral performance. While the mechanisms of contrast gain control in auditory cortex remain unclear, this study highlights potential top-down and bottom-up influences on cortical gain, which may or may not share common neuronal substrates. We believe that the theoretical frameworks and modelling methods presented here will be broadly applicable in future studies of neuronal gain control, a fundamental function of the nervous system.

## Methods

### Animals

All experiments were performed in adult male (n = 19) and female (n = 19) C57BL/6 (Stock No. 000664) or B6.CAST-*Cdh23^Ahl+^* (Stock No. 002756) mice (The Jackson Laboratory; age 12-15 weeks; weight 20-30g). Some of the mice used in these experiments were crossed with other cell-type specific -cre lines, as detailed in Supplementary Table 2. All mice were housed with, at most, five mice per cage, at 28°C on a 12- h light:dark cycle with food provided ad libitum, and a restricted water schedule (see *Water restriction*). All experiments were performed during the animals’ dark cycle. All experimental procedures were in accordance with NIH guidelines and approved by the Institutional Animal Care and Use Committee at the University of Pennsylvania.

### Data availability

All data including spike times from electrophysiological recordings is available on DRYAD: https://doi.org/10.5061/dryad.6djh9w120.

### Surgery

Mice were anesthetized under isoflurane (1-3%). Prior to implantation, all mice were administered subcutaneous doses of buprenorphine (Buprenex, 0.05-0.1 mg/kg) for analgesia, dexamethasone (0.2 mg/kg) to reduce brain swelling, and bupivicane (2 mg/kg) for local anesthesia. In mice implanted with microdrives, two ground screws attached to ground wires were implanted in the left frontal lobe and right cerebellum, with an additional skull screw implanted over the left cerebellum to provide additional support. A small craniotomy was performed over the target stereotactic coordinates relative to bregma, -2.6mm anterior, -4.3mm lateral. Either custom 16-channel microdrives, 32-, or 64-channel shuttle drives (cite) holding tetrode bundles of polyimide-coated nichrome wires were chronically implanted over auditory cortex, and tetrodes were lowered 800um below the pial surface. The exposed tetrodes were covered with GelFoam (Pfizer) or sterile silicone lubricant and sealed with Kwik-Cast (World Precision Instruments). The plastic body of the microdrive and a custom stainless-steel headplate were secured to the skull using dental cement (C&B Metabond) and acrylic (Lang Dental). Mice undergoing only behavioral experiments were implanted with two skull screws in the cerebellum, and a headplate was mounted on the skull as previously described. An antibiotic (Baytril, 5mg/kg) and analgesic (Meloxicam, 5mg/kg) were administered daily (for 3 days) during recovery.

### Water restriction

Following surgical recovery (3 days post-operation), each mouse’s weight was monitored for three additional days to establish a baseline weight. Over the next seven days, mice were water deprived, beginning with a daily ration of 120µL/g and gradually decreasing their ration to 40-50µL/g. During the task, if mice did not receive their full ration, the remainder of their ration was provided in their home cage. Mouse weight relative to baseline was monitored during all stages of water restriction. Additional health signs were used to determine a health score and subsequent treatment plan if a mouse lost more than 20% of baseline weight^82^ as approved by the Institutional Animal Care and Use Committee at the University of Pennsylvania.

### Behavioral apparatus

During the GO/NO-GO task, the mouse was head-fixed in a custom-built, acoustically isolated chamber. A capacitive touch sensor (AT42QT1010, SparkFun) soldered to a lick spout monitored lick activity. Water rewards were dispensed from a gravity fed reservoir, controlled by a solenoid valve (161T011, Neptune Research) calibrated to deliver approximately 4-5µL of water per reward^83^. Low-level task logic – such as lick detection, reward and timeout delivery, and task timing intervals – was directly controlled by an Arduino Uno microprocessor running custom, low-latency software routines. High-level task logic, such as trial randomization, stimulus buffering and presentation, and online data collection and analysis were controlled by custom MATLAB (r2019a, Mathworks) software communicating with the Arduino over a USB serial port. Acoustic waveforms were generated in MATLAB and converted to analog signals via a soundcard (Lynx E44, Lynx Studio Technology, Inc.) or a National Instruments card (NI PCIe-6353) and delivered through an ultrasonic transducer (MCPCT-G5100-4139, Multicomp). The transducer was calibrated to have a flat frequency response between 3 kHz and 80 kHz using a 1/4-inch condenser microphone (Brüel & Kjær) positioned at the expected location of the mouse’s ear^84, 85^. During electrophysiological recording sessions, licks were detected using an optical interrupt sensor (EE-SX771, Omron Automation), to prevent lick-related electrical artifacts introduced by contact with a capacitive sensor.

### Behavioral timeline

Each mouse underwent four stages in the behavioral task: 1) water restriction and habituation, 2) behavioral training, 3) psychometric testing, and, 4) offset testing. During the induction of water restriction, mice were habituated to head-fixation in the behavioral chambers and received water through the lick spout, getting a drop of water for licks separated by more than 2 s. After the mouse began to receive its entire ration by licking in the booth, behavioral training was initiated (typically after 1 week). Each mouse was initially trained and tested in one contrast condition (see *Stimuli*), with the initial training condition counterbalanced across mice. Behavioral performance was monitored during training, and mice were considered trained after completing at least three consecutive sessions with over 80% percent correct. After completing training, behavioral thresholds were measured during at least three sessions in which psychometric stimuli were presented (see *Stimuli*). After estimating the behavioral threshold for each mouse, offset stimulus sets were generated using threshold-level targets. After completion of at least three sessions in the offset task, each mouse was then retrained on the remaining contrast condition. Upon reaching the training criterion of 80% in the new contrast condition, mice were then tested in the psychometric and offset tasks as previously described. For mice in electrophysiological experiments, this sequence of training and testing was continued until the recording site yielded less than three units, or until the mouse stopped performing in the task.

### Stimuli

All stimuli were created in MATLAB and sampled at 192 kHz or 200 kHz and 32-bit resolution. A set of dynamic random chords (DRCs) were created with different contrasts, similarly to those described in previous studies^14, 17, 19^. To construct a DRC, amplitude modulated pure tones were generated at multiple frequencies and then superimposed to create a chord. In some experiments, 34 frequencies were sampled between 4 and ∼40kHz in 1/10 octave steps, in the remaining experiments, 33 frequencies were sampled between 4 and 64kHz in 1/8 octave steps. The amplitude envelope of each tone was generated as follows: every 25 ms, amplitudes for each frequency were sampled from a uniform distribution with a mean of 50 dB and a width of ±5 dB in low contrast or ±15 dB in high contrast. Between each 20 ms chord, the amplitude envelope of each frequency band was linearly ramped over 5 ms to the amplitude value for the next chord, such that the total duration of each chord and its ramp was 25 ms. To synthesize the stimuli, amplitude envelopes were multiplied by a sine wave of their respective frequencies, and summed to produce the final waveform. Each time a set of DRCs was generated, 5 unique random number generator seeds were used to restrict the background noise to 5 distinct scenes (see raster in Figure 6 for an example of spike-locking to the repeated scenes).

In all stages of behavioral training and testing, stimuli created for each trial consisted of a DRC background containing a change in contrast, and the presence or lack of a target at a delay after the change in contrast. Each trial began with 3 seconds of DRC background from one contrast, followed by a switch to the other contrast. Targets consisted of a fixed chord composed of 17 frequencies pseudo-randomly sampled from the frequencies contained in the DRC background, such that the target frequencies were uniformly distributed across the frequency range of the background. To add targets to the background noise, the target amplitude at each target frequency was simply added to a single chord in the amplitude envelope of the background, and linearly ramped: this procedure ensured that target timing was perfectly aligned to changes in the background noise, removing asynchronous timing cues that could be used to detect the target. Target amplitudes are described in values of signal-to-noise ratio (SNR) relative to the average level of the background noise (ie. a 50 dB target embedded in 50 dB background would have an SNR of 0 dB). See Supplementary Table 3 for SNRs used for each mouse. In all trials, targets were embedded after a change in the background contrast, with a delay and level dependent on the current training or testing stage.

### Efficient coding model

We simulated a model neuron that encodes incoming stimuli via an adapting neuronal nonlinearity. Stimuli were drawn from a Gaussian distribution whose mean *μ* was fixed over time but whose standard deviation σ_t_ could switch over time between a low and a high value (σ_t_ = σ^L^ and σ_t_ = σ^H^, respectively). At each time t, a stimulus s_t_ was drawn from the distribution p(s_t_|σ_t_) = N(s_t_^2^; *μ*, σ_t_^2^), transformed via a saturating nonlinearity of the form 1/(1 + e^k(st−s0)^), distorted by Gaussian noise with variance σ^$^, and finally discretized into N discrete levels to generate a response r_t_. This discrete response was linearly decoded to extract an estimate *ŝ_t_* of the current stimulus: *ŝ_t_* = *p*_1_*r_t_* + *p*_0_. The recent history of *L* stimulus estimates was used to update an estimate *σ_t_* of the underlying standard deviation: 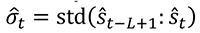. The estimate *σ̂_t_* was then used to select the parameters of the encoder (*k, s_0_*) and the decoder (*p*_1_, *p*_0_) on the next timestep. The encoding and decoding parameters were chosen to minimize the expected error in decoding stimuli given the neuron’s current estimate of the underlying standard deviation: 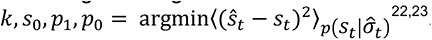.

The parameters of the encoder and decoder were adapted based on a background stimulus with a mean *μ*_B_ that was fixed over time and a standard deviation σ_B_ that switched between low and high values σ^L^ and σ^H^, respectively. We used this adapting nonlinearity to determine how well this model neuron could discriminate target stimuli from this background. Target stimuli were sampled from a Gaussian distribution with a fixed mean *μ*_T_ and with a variance σ_T_ that was scaled in proportion to the variance of the background (*σ*_T_^L^= *fσ*_B_^L^ and *σ*_T_^H^ = *fσ*_B_^H^, respectively). At each timestep, we computed the Bhattacharyya coefficient (BC) of the response distributions produced by background versus target stimuli: 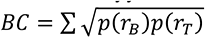. We used 1 – *BC* as our measure of discriminability.

We simulated the behavior of this model using a background “probe” stimulus whose standard deviation switched every *T* timesteps. We simulated *N_c_* cycles of this probe stimulus, where each cycle consisted of *T* timesteps in the low state, followed by *T* timesteps in the high state. This yielded timeseries of the gain *k* and offset *s*_0_ of the adapting nonlinearity, as well as distributions of the neuronal response to the background and target stimuli at each timepoint following a switch in standard deviation. We averaged the gain and offset across cycles to obtain the average properties of the encoder at each timepoint following a switch. We used the distribution of responses to target and background stimuli, measured across cycles, to compute the discriminability at each timepoint following a switch. All simulations were performed with the following values: *T* = 50, *N_c_* = 1,000, *μ_B_* = 0, *μ_T_* = 0 to 3 in 0.25 steps, *σ*_B_^L^ = 1, *σ*_B_^H^ = 3, *f* = 0.25, *σ_n_*^2^ = 0.01, N = 15, L = 12. For Figure 1g, model discriminability in each contrast was fit with a logistic function to estimate the sensitivity and threshold of the model. To approximate the stimulus conditions used in the offset task, the target thresholds for each contrast were then used to select target levels to plot discriminability over time (*μ*_T_^L^ = 1.50, *μ*_T_^H^ = 2.25; Figure 1f).

To compare behavioral performance to the model, we quantified the “psychometric performance” of the model by fitting model discriminability over time with a logistic function:

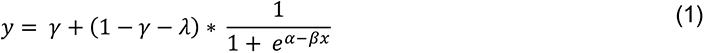

where μ is the x-offset of the function, β determined the sensitivity of the function, γ determined the guess rate (lower bound), λ determined the lapse rate (upper bound) and *x* was stimulus level. α/β determined the threshold of this function, defined as the level corresponding to the steepest part of the curve. This function was fit to behavioral or neuronal performance using constrained gradient descent (*fmincon* in MATLAB) initialized with a 10x10 grid-search of parameters α and β.

To characterize adaptation time constants, adaptation curves were fit with an exponential function

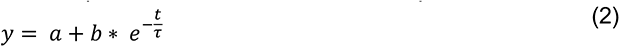

where *a* determined the y-offset of the function, *b* was a multiplicative scaling factor, and r was the time constant of the exponential in units of time t. This function was fit to behavioral or neuronal responses using constrained gradient descent initialized with a 10x10x10 grid search across all three parameters.

To measure the effect of stimulus contrast on parameters of interest, we computed a contrast modulation index (CMI), which measured the relative change in the parameter between low and high contrast:

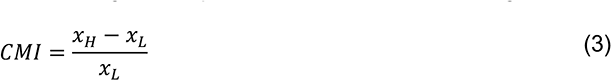

where x is the parameter in question and subscripts H and L denote the value of that parameter in high and low contrast, respectively. The resulting index is 0 if there is no change, 1 if x_H_ is two times larger than x_L_, and is -0.5 if x_L_ is two times x_H_.

### Behavioral task

We used a GO/NO-GO task to measure the detectability of targets in background. In this task, each trial consisted of a noise background with a contrast shift, along with the presence or absence of a target after the change in contrast. Mice were trained to lick when they detected a target (hit), or to withhold licking in the absence of a target (correct reject). This behavior was reinforced by providing a 4-5 µL water reward when the mouse licked correctly (hit), and by initiating a 7-10 s timeout when the mouse licked in the absence of a target (false alarm). Any licks detected during the timeout period resulted in the timeout being reset. In a subset of mice, we introduced an additional trial abort period coincident with the first part of the contrast background, before the contrast switch. Any licks detected in this abort period resulted in the trial being repeated after a 7-10 s timeout, until the mouse withheld from licking during this period. In this task, misses and correct rejects were not rewarded or punished. Trials were separated by a minimum 1.5s inter-trial-interval (ITI). To discourage spontaneous licking, licks were monitored during this period, and if any licks occurred the ITI timer was reset.

To prevent mice from predicting the target, we varied the timing of the target relative to the contrast shift. This required a method for estimating hit rates and false alarm rates at different times during each trial, and to reward and punish the animal during these times in an unbiased manner. To approach this issue, we considered licks only during a 1 s response window after a target presentation (eg. if a target was presented 500 ms post-contrast-switch, the response window persisted from 500 to 1500 ms post-contrast-switch). To apply this method to background-only trials, in which no targets were presented, we considered background trials to be target trials containing infinitely small target amplitudes. For each background trial, we assigned a response window with equiprobable delay matched to the target conditions and considered only licks within those “target” response windows. Thus, over the course of a session, we randomly sampled lick probabilities in background trials during the same temporal windows as those considered during target trials. Using this scheme, we treated target and background-only trials identically, and estimated hit rates and false alarm rates over time in an unbiased manner.

Each mouse performed three stages in the behavioral task: training, psychometric testing, and offset testing. During the training task, trials consisted of two types, background-only trials or target trials presented with equal probability. To facilitate learning, we selected target SNRs at the highest end of the range described previously: in low contrast training sessions, targets were 16 dB SNR, and in high contrast training sessions, targets were 20 dB SNR. To prevent response bias as a function of target timing, we randomly varied the target delay between 250, 500, 750 and 1000 ms after the contrast change in each trial. During the psychometric testing task, there were 7 trial types consisting of background-only trials and target trials spanning six different SNRs (Supplementary Table 3). Based on behavioral piloting, we presented high SNR trials with a greater probability, to ensure that mice were consistently rewarded during the task. In low and high contrast psychometric sessions, the probability of a background trial was 0.4, the probability of the four lowest target SNRs was 0.05 each, and the probability of the two highest target SNRs was 0.2 each. As in training, target timing was varied randomly between 250, 500, 750 and 1000 ms after the contrast change in each trial. After completing at least three sessions of the psychometric task, stimuli were generated for the offset testing task. This task consisted of 15 unique trial types: 3 target levels (background trials, threshold target trials, and high SNR target trials), and 5 target delays relative to the contrast change (25, 75, 225, 475, 975 ms delay). Threshold target amplitudes were determined individually for each mouse by fitting performance averaged over several sessions with a psychometric function, and extracting the level at which the slope of the psychometric curve was steepest. Based on behavioral piloting, background trials, threshold target trials, and high SNR target trials were presented with probabilities of 0.4, 0.2, and 0.4, respectively. Target delay on each trial was selected with equal probability. In all behavioral stages, trial order was pseudorandomly generated, such that there were no more than three target or background trials in a row.

A subset of mice (n = 2), were presented targets in silence (Figure 4). To generate this stimulus set without changing the basic structure of the task or stimuli, we simply took the spectrograms of all stimuli containing 25 dB SNR targets from the low-to-high contrast stimulus sessions, and set the stimulus power flanking each target to zero. This manipulation was only performed in the target period, and the low contrast adaptation period of the trials remained the same. Thus, the targets and adaptation periods were identical to those presented in the target-in-background task. To vary the difficulty of the task, the level of the target was attenuated using the following values: -75, -60, -45, -30, -15, and 0 dB attenuation relative to the 25 dB SNR target. Mice were previously trained in the target-in-background task prior to performing the target in silence task. Before psychometrically varying the target attenuation, mice were trained in the new task to criterion performance. Mice generalized very rapidly to the new task, reaching 97% and 94% training accuracy on the first day of exposure to targets in silence (mice CA124 and CA125, respectively).

### Chronic muscimol application

A separate cohort of mice (n = 4) were bilaterally implanted with 26 GA guide cannulae (PlasticsOne, C315GMN-SPC mini, cut 5 mm below pedestal) in auditory cortex. The surgery was performed as described above with the following modifications. After the skull was leveled using a stereotax, two small craniotomies were performed -2.6 mm anterior, ±4.3 mm lateral from bregma, over auditory cortex. The guide cannulae and dummy infusion cannulae (PlasticsOne, C315DCMN-SPC mini, cut to fit 5 mm C315GMN with a 0.5 mm projection depth) were sterilized in an autoclave. The dummy cannulae were partially screwed into the guide cannulae and placed in a stereotaxic clamp. After zeroing the tip of the guide cannula to the brain surface, the cannula was lowered to 500 μm below the cortical surface. This depth was chosen because the infusion cannulae (PlasticsOne, C315LIMN-SPC mini) project 500 μm from the end of the guide cannulae when completely inserted, leading to a final depth of 1000 μm – the location of auditory cortex. The dummy cannulae were then fully inserted and this procedure was repeated for the next cortical hemisphere.

Prior to injecting, two injection syringes (Hamilton Syringe, 10μL Gaslight #1701) and tubing (C313CT tubing 023x050 PE50) were backfilled with mineral oil. Sterilized infusion cannulae were then attached to each syringe and ∼500nL of muscimol (diluted with 1x PBS to .25 mg/mL; Sigma Aldrich, M1523) or 0.9% sterile saline was drawn up into the injection cannulae using a dual injector (Harvard Apparatus, Pump 11 Pico Plus Elite). The mouse was then headfixed and the dummy cannulae were removed and sterilized. The loaded infusion cannulae were then screwed all the way into the guide cannulae and 400 nL of muscimol or saline was infused bilaterally at a rate of 250 nL/minute. The infusion cannulae were then replaced with the dummy cannulae and the mouse rested in its home cage for 30-45 minutes before beginning the behavioral session.

### Acute electrophysiological recordings

For acute recordings used to fit the GC-GLM model (Figure 2), neuronal signals were recorded from n = 1 awake, untrained mouse. Prior to the recording session, the mouse was anesthetized and a headpost and ground pin were implanted on the skull (see *Surgery*). On the day of the recording, the mouse was briefly anesthetized with 3% isoflurane and a small craniotomy was performed over auditory cortex using a dental drill or scalpel (∼1 mm x 1 mm craniotomy centered approximately 1.25 mm anterior to the lambdoid suture along caudal end of the squamosal suture). A 32 channel silicon probe (Neuronexus) was then positioned perpendicularly to the cortical surface and lowered at a rate of 1-2 μm/s to a final depth of 800-1200 μm. As the probe was lowered, trains of brief noise bursts were repeated, and if stimulus locked responses to the noise bursts were observed, the probe was determined to be in auditory cortex. The probe was then allowed to settle for up to 30 minutes before starting the recording. Neuronal signals were amplified and digitized with an Intan headstage (RHD 32ch) and recorded by an openEphys acquisition board^86^ at a rate of 30 kHz.

For this experiment, the mouse was presented with 3 s DRCs alternating between low and high contrast (uniform distribution with a mean of 50 dB and a width of ±5 dB in low contrast or ±15 dB in high contrast at a chord rate of 25 ms, as described in *Stimuli*). In order to accurately fit the GLM in an unbiased manner, these stimuli were highly random, composed of 100 unique chord patterns for each contrast (Supplementary Figure 2i,j). For each of the two recording sites, 5 repeats of this stimulus set were played.

### Behavioral electrophysiological recordings

Neuronal signals were acquired from awake, behaving mice as they performed the psychometric and offset testing tasks described previously. Chronically implanted, 16-, 32-, or 64-channel microdrives^86, 87^ were connected to one or two 32 channel Intan amplifier headstages. Amplified signals were recorded at 30 kHz using an openEphys acquisition board via an SPI cable, where the signals were digitized.

For all recordings, broadband signals were filtered between 500 and 6000 Hz, offset corrected, and re-referenced to the median across all active channels. The preprocessed data was then sorted using KiloSort^88^ or KiloSort2 and the resulting clustering was manually corrected in phy2 according to community-developed guidelines. The resulting units were labelled as single units if they exhibited a clear refractory period and did not need to be split. Splitting assessments were made through manual examination of principle component features for the two best channels of a cluster. If two noticeable clusters in feature space were evident in a unit, the unit was either manually split, or classified as a multiunit.

### Generalized linear model

To justify the form of GLM used here, we discuss a how a model neuron could implement gain control in the simplest terms, and then structure our inference model to extract the parameters of this model neuron. We will assume that the activity of the model neuron is driven by three sources: 1) stimulus drive, 2) stimulus contrast, and 3) the multiplicative interaction between the two, which we use to define the gain (for a formal definition of this forward model and the inference model, see *Supplementary Information*).

As discussed previously, the stimulus used in our experiments is composed of many frequencies that change in loudness in discrete time steps:

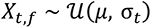

where *X_t,f_* is the stimulus spectrogram that varies as a function of time *t* and frequency *f*. Each time and frequency bin of *X* is sampled from a uniform distribution defined by an average value *μ* and contrast *σ_t_*.

We assume that the hypothetical neuron responds selectively at some frequency and time lag, defined by a filter, or STRF *β_h,f_* with history *h* and frequency *f* components. Given β, we can define the stimulus drive x_t_ as

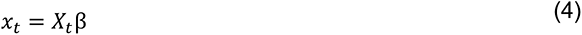

where at each time *t*, *X*_t_ is a row vector of size *F* frequencies times *H* lags (ie. the “unrolled” lagged stimulus spectrogram) and β is the STRF unrolled to a single column vector of the same size.

In the spirit of efficient coding theory, and as shown in previous work, we assume that the gain *g* of the neuron should be inversely proportional to the contrast, such that g(σ) α 1/σ (ie. when contrast is low gain should be high, and vice-versa). We also define “neutral” gain to be the average of the gain of the neuron in low and high contrast. Putting these two features together, we can summarize the gain of the neuron as

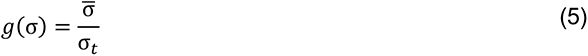

where σ̄ is the harmonic mean of the contrast in the low and high conditions (see *Supplementary Information*). In the case of a 3-fold change in contrast, this function constrains the gain of the neuron between 0.5 and 1.5, with a neutral value of 1. As mentioned previously, we consider gain to be the multiplicative interaction between the stimulus drive and the contrast, such that the contribution of gain control to the response of the neuron is related to 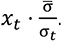.

To summarize, we considered a hypothetical neuron driven by the stimulus according to a STRF β and by the interaction between the stimulus drive and the contrast 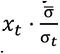. To infer the relative weights of each of these components of the neuronal response, we defined a Poisson GLM with an intercept term and the following predictors:

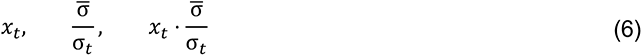

In other words, the model is composed of a stimulus predictor *x_t_*, a contrast predictor *σ̄*/*σ_t_*, and their interaction. Therefore, the GLM models the firing rate λ at time *t* as a Poisson distribution with the following mean:

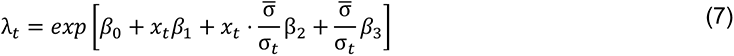

where *β*_0_ … *β*_3_ are the parameters to be inferred. Based on our behavioral data (Figure 3) and the predictions of the efficient coding model (Figure 1), we expected the influence of contrast on neuronal gain to be asymmetric and smooth. To enable the GLM to capture both of these qualities, we first defined the contrast predictors from a set of cubic B-spline temporal basis functions, then defined separate contrast predictors for transitions to low and high contrast. Incorporating these changes, we can redefine equation 4 above as

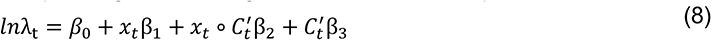

where ° denotes element-by-element “broadcasting” multiplication and *C_t_*’ is a matrix of contrast predictors *σ*/*σ̄_t_* convolved with a set of basis functions and separated by contrast transitions (see *Supplementary Information*). For the sake of clarity, note that in the expression above, *β_0_* is a number, *x* is a column vector of length *T*, β_+_ is a number, C’ is a *T*-by-2*B* matrix, and *β*_2_ and *β*_3_ are column vectors of length 2*B*, where *B* is the number of splines.

So far, we outlined a hypothetical neuron which implements gain control, and a GLM with which we can approximate the behavior of this neuron. Next, we describe how to use the fitted parameters to quantify the gain of the neuron. Conceptually, an increase or decrease in the gain of the neuron is analogous to more or less sensitivity to small changes in the stimulus. Based on this intuition, we focused on how the response of the neuron (as modelled by a fitted GLM) is expected to change between conditions where the gain is expected to contribute (ie. in the presence of gain control) and where it is not (ie. in the absence of gain control, where gain is “neutral”). Following this logic, we derived a definition for gain *w_t_* as the ratio between the sensitivity of the fitted model with changes in contrast, compared to the sensitivity of the same model when the contrast is at a reference value, which we defined previously as *w_t_* = 1:

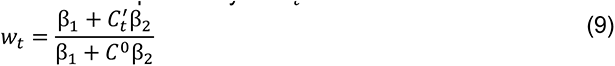

where *w_t_* is the estimated gain at time *t*, and C^0^ is a reference contrast design matrix identical to *C_t_*’ except that all non-zero elements are set to 1 (see *Supplementary Information* for full derivation of *w_t_*).

To fit the model, we implemented a two-step procedure. In the first step, the STRF β of the neuron was estimated according to the model

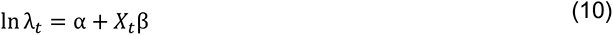

For the second step, we calculated the stimulus drive as described in equation 4, and then fit equation 8 to the data for each neuron using glmfit in MATLAB (v2018a). This entire fitting procedure was 10-fold cross-validated with folds stratified across trials of each contrast and run in parallel on a high performance computing cluster using custom BASH scripts (v3.2.57(1)). In the first step, we fit the STRF β with *F* frequency bins according to the stimulus spectrogram (*F* = 33 or 34, see *Stimuli*) and a history window of 300 ms (*H* = 12). When fitting the full model, we defined the contrast design matrix *C_t_*^’^ to capture 1000 ms of contrast history around each transition (*H*’ = 40), convolved with a set of B-spline temporal basis functions^89^ (here, we used B-splines with a degree of 3 and 3 equally-spaced knots, constrained to go smoothly to zero at the longest lag, which implied that *B* = 4).

To validate the model, we first simulated neurons according to the forward model outlined above (Supplementary Figure 2a) while varying the amount of gain control and the temporal trajectory of gain in different simulation runs. We found that the GLM accurately predicted the STRF shape, spike rates and gain trajectories across a variety of simulation parameters (Supplementary Figure 2c, e-h). For a detailed description and discussion of the simulation results, see *Supplementary Information* and Extended Data Table 4.

### Behavioral and neuronal detection performance

To calculate performance in the target-in-background detection task we adopted commonly used signal detection theory methods^39, 90^ to estimate the ability of an ideal observer to discriminate between two sensory distributions: in our case, a distribution for target trials and a distribution for background trials. When analyzing behavior, we computed the percent correct performance of an ideal observer^91^ as a function of the probability of hits and false alarms:

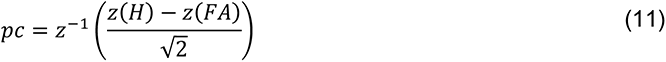

where z^%+^is cumulative probability of the normal distribution (normcdf in MATLAB), z is the inverse of the normal distribution (ie. the z-score, norminv in MATLAB), H is the hit rate, and FA is the false alarm rate. For psychophysical performance, hit rates and false alarm rates near 0 and 1 were adjusted using the log-linear rule^92^, to reduce biases in performance estimation resulting from low numbers of trials.

To calculate neuronal performance in the same reference frame as the behavior, we employed similar ideal observer techniques. First, neuronal responses (either spike rates of single units, or population projection values), were averaged in a 100ms window post target onset (for background trials, this window was randomly chosen on each trial to coincide with target presentation times on target trials). Then, using the distributions of responses during target and background trials, we computed receiver-operating-characteristic curves and took the area under the curve (AUC) as the percent correct of an ideal observer discriminating between the target and background distributions. To determine whether the AUC value for a given set of trial distributions was significantly different from chance, we performed a bootstrap procedure where we sampled from all the trials with replacement 500 times and recomputed AUC for each sample. If the 95% confidence intervals for this bootstrapped distribution did not include chance (.5), we defined that AUC value as significant. For population analyses which generated single-trial predictions, neuronal hit and false alarm rates were transformed to percent correct as described above.

### Population response metrics

On sessions where three or more neurons were simultaneously recorded, we used a population vector technique^27^ to estimate the ability of neuronal populations to discriminate targets from background. First, spike rates in each trial were averaged in a 100ms window post-target onset. Then, using a leave-one-out procedure, we computed a trial averaged population vector for target trials, *v_T_*, and a separate average population vector for background trials, *v_B_*. We then estimated the coding direction in high dimensional neuronal space that best separated the target and background responses: *CD* = *v_T_* – *v_B_* The held out trial was then projected along this dimension, by taking the population response vector on that trial *v_trial_* and projecting it along the estimated coding direction using the dot product: projection value = *v_trial_*\**CD*. This procedure was repeated holding out each trial, and estimating the coding direction from the remaining trials. For psychometric testing sessions, the target responses from the two loudest target levels were used to estimate coding direction, and in offset testing sessions the target responses from the high SNR target trials were used. After computing projections for every trial, the resulting matrix was normalized between 0 and 1.

### Population classifier

Based on previously described methods^28^, we used a criterion-based decision rule to estimate how a hypothetical down-stream neuron may read out the neuronal activity of a population of neurons. As before, trial distributions of neuronal responses to targets or background were created from the average activity in a 100ms window post-target. Then, we sampled 100 criterion values between the minimum and maximum response, and for each criterion estimated the proportion of correct trials under two decision rules: 1) report target present if the response is greater than the criterion, or, 2) report target present if the response is less than the criterion. By assessing these two decision rules, neurons that were suppressed by target presence were treated equally to neurons that were enhanced by target presence. Finally, we chose the criterion and decision rule that yielded the highest proportion of correct trials, and computed neuronal hit rates and false alarm rates for each target level, and background-only trials. These hit rates and false alarm rates were then transformed to percent correct according to Equation 8.

### Linear-nonlinear model

First, we selected only neurons in the dataset which had reliable responses to stimulus repeats. To determine response reliability, we computed a noise ratio (NR) for each neuron, which describes the amount of variability in the response due to noise versus the amount of variability in the response driven by the stimulus^93, 94^. Values approaching 0 indicate increasingly reliable responses to the stimulus, so for the remaining analyses, we included neurons with NR < 100.

The linear nonlinear model was composed of a spectrotemporal receptive field (STRF) and a set of rectifying nonlinearities. The STRF β was fit using normalized reverse correlation

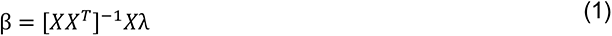

where *X* is the stimulus design matrix *X_t_* defined in equation 4 and λ is the spike count in each 25 ms bin of the DRC stimulus. When defining *X*, we used a history window of 300 ms (*H* = 12) and frequency bins corresponding to the frequencies composing the dynamic random chords (see *Stimuli*). After fitting the STRF, we fit the nonlinearities of the neuron. This two-step fitting procedure was repeated using 10 fold cross-validation, as described below.

For each fold, we selected 90% of the trials for training, leaving the remaining 10% to be held out for testing. Within each trial, we excluded neuronal responses around transitions from silence, or transitions in contrast, to prevent the model from overfitting strong transients in the neuronal response. Additionally, we excluded neuronal responses within a 50ms window after target presentation, to prevent overfitting of target responses. Given these exclusion criteria, we calculated the duration of stimulus sampled in the target period for each trial, and, for each trial, sampled the same duration of stimulus within the adaptation period. This procedure ensured that the model was fit to the same amount of high and low contrast stimulation per trial, to minimize overfitting to one contrast condition. Then, a stimulus design matrix *X* was defined using these stimulus periods, and the STRF was fit using equation 12. We tested whether STRF properties were affected by stimulus contrast, and found STRFs to be largely stable when estimated separately for each contrast (*Supplementary Information* and Supplementary Figure 7). Therefore, we used both periods of contrast to estimate β.

Using the STRF fit to the training data, we computed the linear drive x_t_by convolving the STRF with the lagged spectrogram of the training stimulus (equation 4). For the GC-LN model we separated the linear predictions into low and high contrast periods, while for the static-LN model all matched time points were used. We generated a histogram of the linear prediction values (50 bins), and for each bin, computed the mean spike rate of the neuron when the linear prediction fell within those bin edges (Figure 6d, scatter points). The resulting set of linear prediction values and average spike rates were fit with an exponential function:

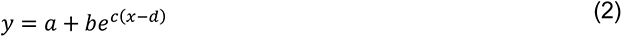

where a determined the minimum firing rate, *b* was a multiplicative scaling factor, c determined the gain of the exponent, and d determined the x-offset, or firing threshold of the neuron. This function was fit to each cell using constrained gradient descent (fmincon in MATLAB), using a 10x10 grid search for parameters *b* and c. The gain for each neuron was defined as c. This entire process was repeated for each cross-validation fold, and the final parameter estimates for the STRF and nonlinearities were taken as the average over the 10 runs.

To determine the relationship between neuronal gain and behavioral performance, we computed the average neuronal gain across all noise responsive neurons (NR < 100) in each session for the adaptation and target periods in the trial. We then compared the session-averaged gain values to the fitted thresholds and slopes of the psychometric curve across sessions using the mixed-effects linear models outlined in the main text.

### Inclusion criteria

Unless otherwise noted, behavioral sessions in which the false alarm rate exceeded 20% were discarded from analysis. One mouse (ID: CA122) had consistently high false alarm rates in the high contrast condition, so we excluded high contrast sessions from this mouse from all analyses. For Figures 5 and 6, we removed neurons with low spike rates (<1Hz) and noise-like or inverted (ie. upward inflected) spike waveforms. To determine waveform quality, we computed the width of each waveform at half of the minimum value (FWHM) and its correlation with the average waveform over all neurons. Neurons whose waveforms had outlier FWHM values (isoutlier; in MATLAB), negative correlations, or were not significantly correlated with the average (Bonferoni corrected *p* > 5.85e-6) were removed from further analysis. For Figure 5f-l, sessions with stable population decoding performance were included (defined as sessions where more than half of the target types elicited significant population AUC values, as determined by the bootstrap procedure described previously). For Figure 6e-h, only neurons with noise ratios less than 100 were included in all analyses.

## Data Availability

The data generated in this study, including sorted spike times from electrophysiological recordings and behavioral data, have been deposited on Dryad: https://doi.org/10.5061/dryad.6djh9w120. Every Figure and Supplementary Figure has associated data with the following exceptions. Figure 1, Supplementary Figure 1, and Supplementary Figure 2 are the results of simulations and thus have no associated raw data. Simulation parameters can be found in the code. There are no restrictions on data availability. A Source Data file has been provided with this manuscript.

## Code Availability

Code for behavioral and neural analysis: https://github.com/geffenlab/contrast_behavior. Code for the GC-GLM model and simulations: https://github.com/geffenlab/contrast_glm.

## Supporting information

Supplemental figures and Statistics table

Source data for figures

## Acknowledgements

This work was supported by NIDCD R01DC015527, R01DC014479, NINDS R01NS113241 to MNG and NIDCD F31DC016524 to CFA. AMW was supported by NIDCD T32DC016903. The authors would also like to thank Dr. Yale Cohen, Dr. Nicole Rust, and Dr. David Brainard for their helpful feedback on this study and Dashiell Ridolfi-Starr, Karmi Oxman, Nitay Caspi, Tyler Ling, and Margaret DenBleyker for experimental assistance.

## Author Contributions Statement

CFA and MNG conceived of the behavioral and neural recording experiments. WM and AMH conceived of and implemented the normative gain control model. EP and CFA conceived of and implemented the GC-GLM model simulations and fitting procedures. AMW performed the acute muscimol recordings (Supplementary Figure 5). CFA, KCW and LG created the behavioral apparatus and performed behavioral experiments. CFA performed the surgeries, electrophysiological behavior experiments and muscimol behavior experiments. KCW performed acute electrophysiology experiments (Figure 2). CFA analyzed the data. CFA and MNG wrote the paper.

