## Supplemental figures and Statistics table for "Dynamics of cortical contrast adaptation predict perception of signals in noise"

**(8 figures, 4 tables, supplementary procedures, supplementary results)**

Supplementary Figure 1: Normative model responses, predictions, and example response distributions.
Supplementary Figure 2: Simulation results to validate the GC-GLM.
Supplementary Figure 3: GC-GLM comparison to a dynamic contrast adaptation model.
Supplementary Figure 4: Behavioral slopes are affected by the target level range.
Supplementary Figure 5: Confirmation of cortical inactivation with muscimol.
Supplementary Figure 6: Histological tracing of tetrode tracks.
Supplementary Figure 7: STRFs are unaffected by contrast during behavioral performance.
Supplementary Figure 8: Relationship between behavior and gain outside of the target period.
Supplementary Table 1: Statistical comparisons.
Supplementary Table 2: Mouse strains and genders.
Supplementary Table 3: Target SNRs used during psychometric testing.
Supplementary Table 4: GLM simulation parameters.
Supplementary Experimental Procedures
Supplementary Results
Graphical Abstract

### Supplementary Figures

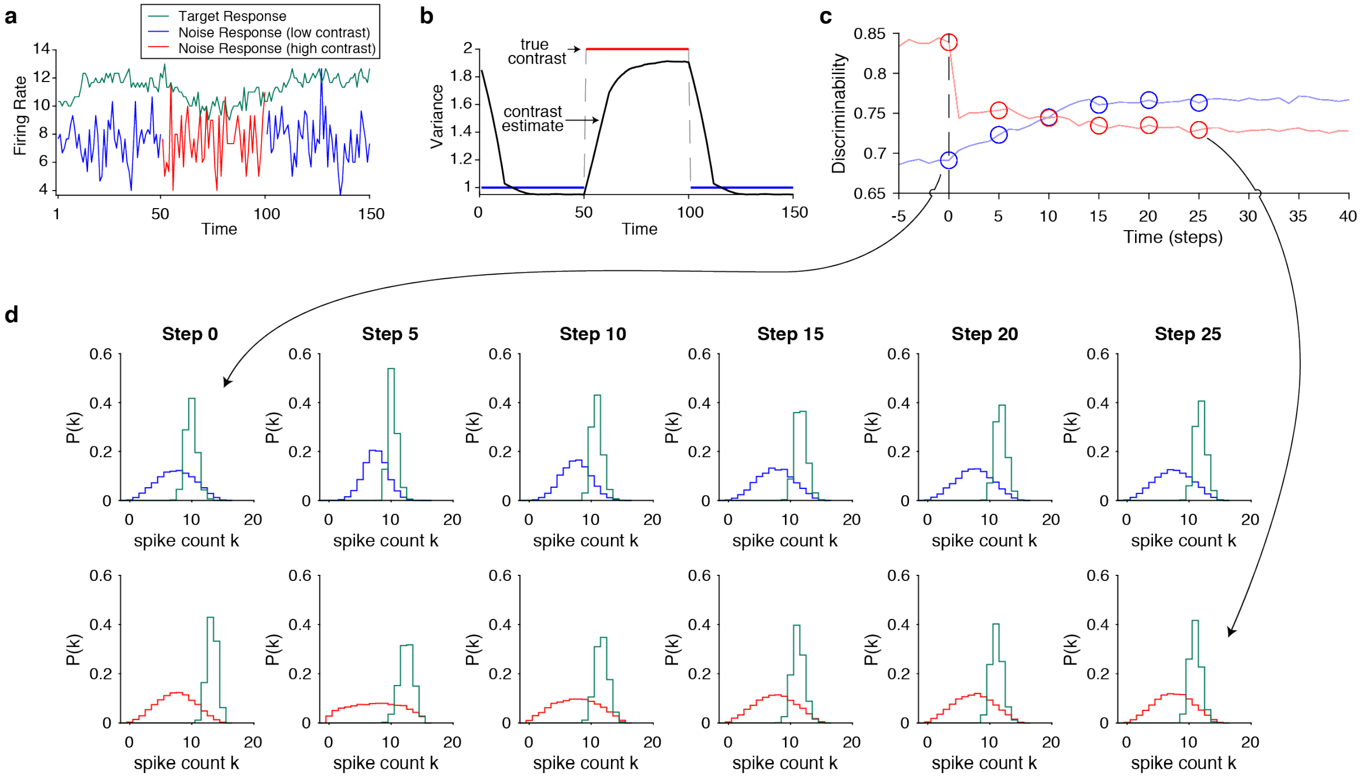


**Supplementary Figure 1 (related to Figure 1). Normative model responses, predictions, and example response distributions.**

**a,** The firing rate of the simulated neuron as a function of time. Traces shaded in blue or red indicate the firing rate to periods of low or high contrast background noise, respectively. The green trace indicates the model response to overlaid targets. **b,** The true contrast (labelled as variance) of the stimulus (blue, red, and dashed gray lines) along with the average model estimate of the contrast (solid black line) over time. **c,** Discriminability as a function of time and contrast, with the trace color indicating the contrast after the switch. The dashed vertical line indicates the time of the contrast switch. Open circles indicate time samples used to plot the distributions in **d**. **d,** Target (green) and background (blue or red) distributions as a function of time and contrast. The top row includes responses to targets and background in low contrast. Each column denotes a different time step relative to the change in contrast, as indicated by the column title. The bottom row is the same, but for high contrast. Arrows between **c** and **d** indicate distributions which yielded the indicated value of discriminability in the trace.

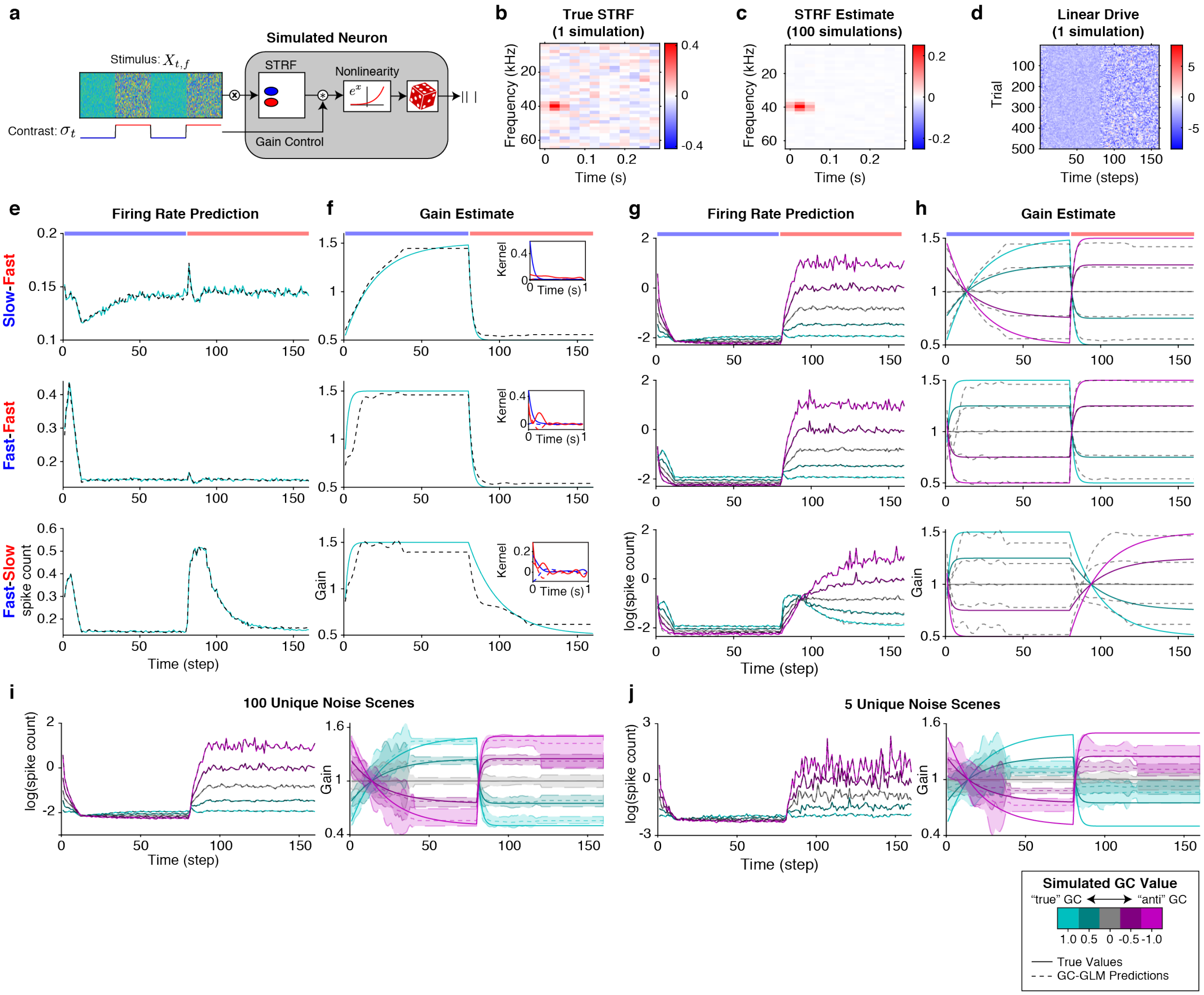


**Supplementary Figure 2 (related to Figure 2). Simulation results to validate the GC-GLM.**

**a,** Schematic of simulated neurons in the forward model. Each neuron received broadband noise inputs which changed contrast every 2s ($X_{t,f}$). A STRF modelled by a 2D-gaussian function with added noise filtered the stimulus to generate a linear response. This filter response was then modulated by a gain control function, which controlled the amount and time-course of gain control based on the stimulus contrast. This gain modulated output was then exponentiated and stochastic spikes were generated using a Poisson process. **b,** Example STRF from one simulated neuron. Colorbar indicates STRF magnitude. **c,** Model estimate of the STRF averaged across 100 simulated neurons. **d,** Example linear drive for one simulated neuron over 500 trials (ie. the filter response of the STRF convolved with the stimulus). **e,** Each panel plots the average firing rates of 100 simulated neurons (solid teal lines) and corresponding GC-GLM fits (dashed black lines) when simulating perfect gain control (GC = 1.0). Each row corresponds to 100 simulations of different gain time courses, with the top row depicting a slow transition to low contrast, with a fast transition to high contrast. The middle row plots simulations were both transitions were fast. The bottom row plots simulations where the transition to low contrast was fast, with a slow transition to high contrast. The corresponding rows of panels **f**, **g**, and **h**, are the results of simulations with the same gain time courses. **f,** Average gain time-course of the simulated neurons (solid teal lines) and the corresponding GC-GLM estimate of the gain, $w$, averaged over 100 simulations (black dashed lines). Insets of each panel depict the contrast kernels (dashed lines) and gain kernels (solid lines) estimated for each contrast. Blue lines indicate kernels after a switch to low contrast and red lines indicate kernels after a switch to high contrast. **g,** Average log firing rate for simulations with different gain time-courses and different degrees of gain control (GC value; the legend in the lower right indicates the color-GC value mapping). Each plotted line indicates the average firing rate/prediction for 100 simulations. **h,** Average gain time-course of all simulations (solid colored lines) and the average estimates of $w$ (dashed gray lines). **i,** Simulations with 100 unique stimulus scenes, repeated 5 times each. Left panel plots the average firing rates and model fits. Right panel plots the true gain time-course (solid lines) and the average model gain estimate, $w$ (dashed lines). The shaded areas indicate 2.5 and 97.5 percentiles of the gain estimates. **j,** Simulations with 5 unique stimulus scenes, repeated 100 times each. Formatting as in **i**. For panels **e-j**, the GC value colors and line formatting are indicated in the legend on the bottom right.

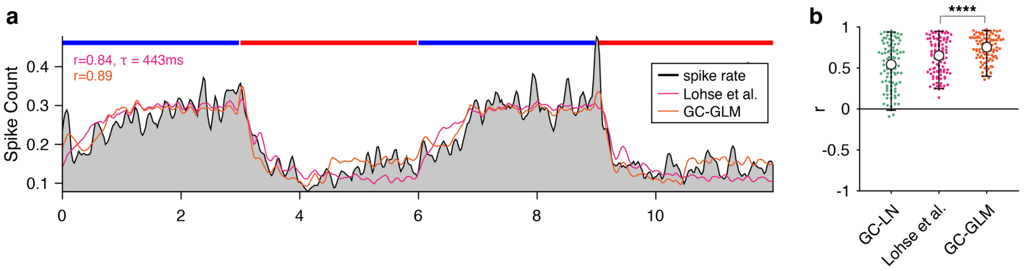


**Supplementary Figure 3 (related to Figure 2). GC-GLM comparison to a dynamic contrast adaptation model.**

**a,** Example model fits to the neuron plotted in Figure 2d. Grey volume indicates the smoothed spike count, while overlaid pink and orange lines indicate fits of a previously published contrast adaptation model[19] and the GC-GLM described in this study, respectively. Inset text indicates the correlation between the model prediction and the firing rate, and the adaptation time constant for the model from Lohse et al. **b,** Model performance for each neuron quantified by cross-validated correlation between the trial averaged firing rate and the trial-averaged model prediction. Colors as in a, with green dots representing the GC-LN model correlations, for comparison. A two-way sign-rank test between the Lohse et al. model and the GC-GLM indicated that the GC-GLM was a better fit to the data (n = 97 neurons; *median* ± *IQR difference in r*: 0.065 ± 0.12, $Z$ = 7.66, $p$ = 1.93e-14).

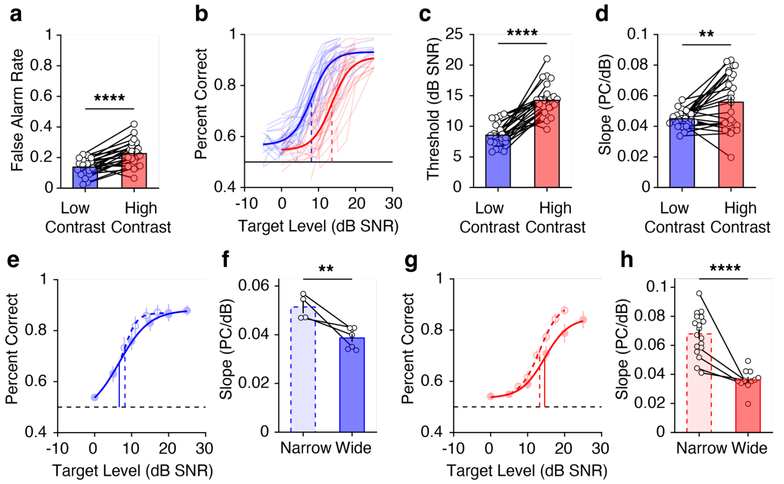


**Supplementary Figure 4 (related to Figure 3). Behavioral slopes are affected by the target level range.**

**a,** The effect of contrast on the false alarm rates in psychometric sessions (n = 25 mice). Each dot and line represent a mouse, the blue and red bars indicate the mean false alarm rate for low and high contrast ±SEM. Results of a two-way paired t-test ($t(23) = -6.16$, $p = 2.75e-6$) across contrast revealed a significantly higher false alarm rate in high contrast (Mean ($M$) = 0.22, standard deviation ($std$) = 0.080) compared to low contrast ($M$ = 0.13, $std$ = 0.054). **b,** Psychometric curves for all mice (n = 25). Thin traces are curves of individual mice, overlaid with psychometric fits to the entire dataset. **c,** Comparison of psychometric thresholds for the curves plotted in b, formatted as in a. Asterisks indicate results of a two-way paired t-test ($t(23) = -8.19$, $p = 2.88e-8$). **d,** Slope of the psychometric curves plotted in b, formatted as in a. Asterisks are results of a two-way paired t-test ($t(23) = -3.42$, $p = 0.0024$). **e,** Average psychometric curves and percent correct for mice presented with a narrow range of targets (n = 4 mice, range = 15 dB SNR; Supplementary Table 3, row 4; dashed lines and open dots), and those presented with a wide range of targets (n = 7 mice, range = 25 dB SNR; Supplementary Table 3, row 1; solid lines and filled dots) in low contrast. Error bars indicate ±SEM. **f,** Psychometric slopes for mice in e. Each bar indicates the mean for each condition ±SEM. A two-way unpaired t-test ($t(9) = 4.48$, $p = 0.0015$) indicated significantly steeper slopes in response to narrow target distributions ($M$ = 0.051, $std$ = 0.0051) compared to wide target distributions ($M$ = 0.039, $std$ = 0.0042). **g**, Average psychometric curves and percent correct for mice presented with a narrow range of targets (n = 18 mice, average of ranges = 12 and 15 dB SNR; Supplementary Table 3, rows 4 and 5; dashed lines and open dots) or wide range of targets (n = 12 mice, range = 25 dB SNR; Supplementary Table 3, row 1; solid lines and filled dots) in high contrast. Error bars indicate ±SEM. **h,** Psychometric slope for each mouse in g. Formatting as in f. Error bars indicate ±SEM. A two-way unpaired t-test ($t(28) = 7.00$, $p = 1.30e-7$) indicated significantly steeper slopes in response to narrow target distributions ($M$ = 0.068, $std$ = 0.015) compared to wide target distributions ($M$ = 0.036, $std$ = 0.0070) in high contrast. In all plots: $^{ns}p>0.1$; $^{\dagger}p<0.1$, $^{*}p<0.05$, $^{**}p<0.01$, $^{***}p<0.001$, $^{****}p<0.0001$.

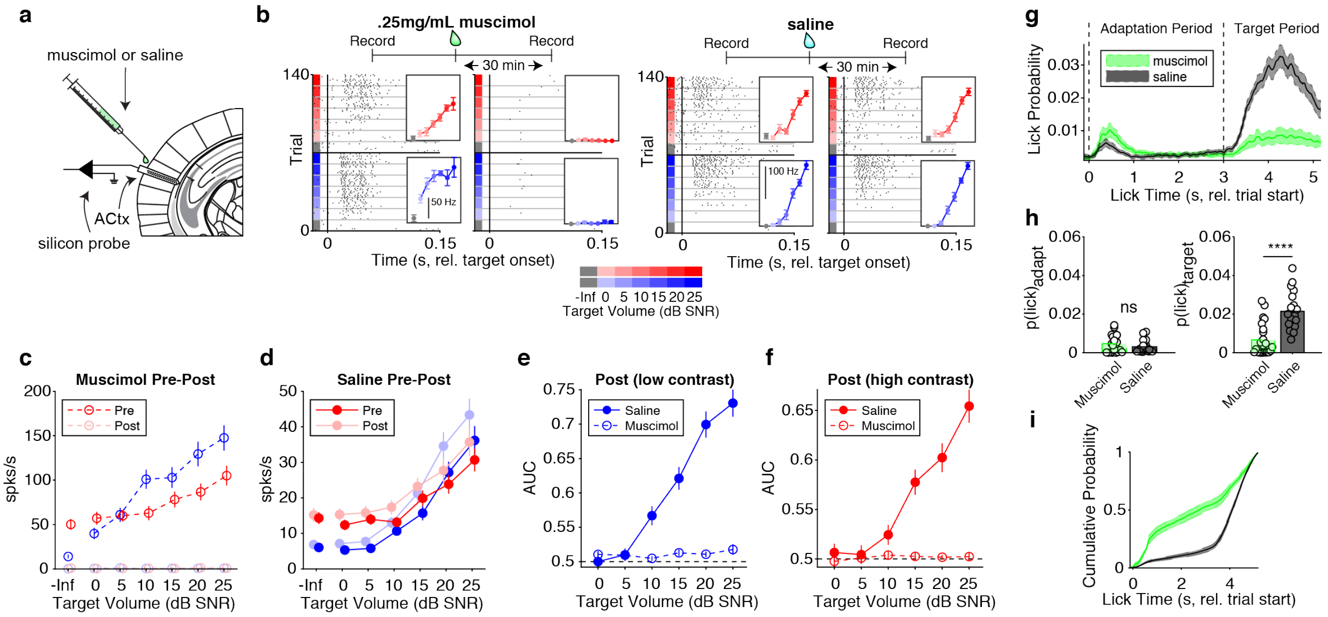


**Supplementary Figure 5 (related to Figure 4). Confirmation of cortical inactivation with muscimol.**

**a,** Setup schematic for acute muscimol recordings in ACtx. **b,** Example spike rasters from two different neurons pre- and post-muscimol or saline application. On top of the raster is the timeline for each recording. Rasters are sorted by contrast and target level, with color indicating low or high contrast backgrounds, color shade indicating target level, and gray indicating background-only trials (-Inf). *Left panel:* spike raster of a representative neuron recorded prior to muscimol application, followed by the raster for the same neuron 30 minutes after muscimol application. *Insets:* Mean firing rate for each condition. Shade indicates target level and the scale bar indicates the firing rate. Error bars are ±SEM across trials. *Right panel:* Example neuron before and after application of saline. Formatting as in left panels. **c,** Firing rates before and after muscimol application as a function of target level and contrast. Dark dashed lines indicate spike rates recorded pre-muscimol application and light dashed lines indicate the responses post-application. Error bars indicate ±SEM across neurons (n=104). **d,** Firing rates before and after saline application. As in c, dark lines are responses recorded prior to saline application and light lines indicate responses recorded after saline application. In c and d, blue and red plots indicate responses during low contrast and high contrast, respectively, and the circles not connected by a line and labelled "-Inf" are responses to background alone. Error bars indicate ±SEM across neurons (n=42). **e,** Area under the ROC curve (AUC) averaged across neurons after drug application in muscimol and saline recording sessions in low contrast. Filled circles and solid lines are responses after saline was applied while open circles and dashed lines are responses after muscimol was applied. Error bars indicate ±SEM across neurons (muscimol, n=104; saline, n=42). **f,** Same as e, but for high contrast. **g,** Lick probability over time during muscimol or saline sessions (n=23 muscimol, n=21 saline sessions for g, h, i). Dashed vertical lines indicate trial onset (0 s) and the contrast switch (3 s). Green traces are muscimol sessions and black traces are saline sessions. The shading around each trace indicates ±SEM across sessions. **h,** *Left:* comparison of lick probability during the adaptation period (two-way rank-sum test: $p$ = 0.81). *Right:* comparison of lick probability during the target period (two-way rank-sum test: $p$ = 2.34e-5). Each circle indicates a session and color is as in g. Error bars indicate ±SEM across sessions. **i,** Cumulative probability of licking throughout the trial, normalized within muscimol or saline conditions to sum to 1. Colors as in g, h. Shading indicates ±SEM across sessions. In all plots: $^{ns}p>0.1$; $^{\dagger}p<0.1$, $^{*}p<0.05$, $^{**}p<0.01$, $^{***}p<0.001$, $^{****}p<0.0001$.

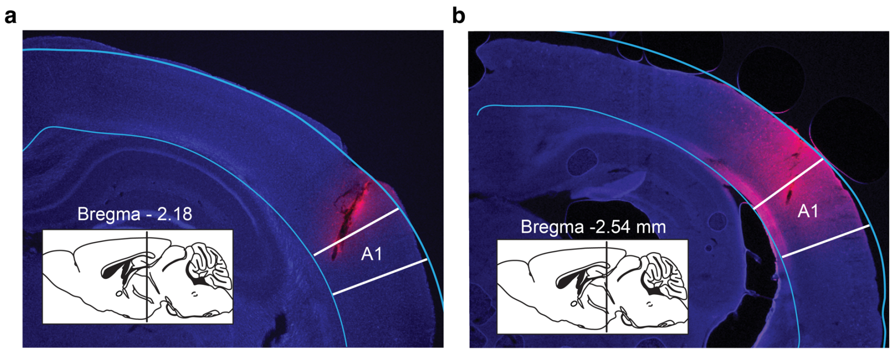


**Supplementary Figure 6 (related to Figure 5). Histological tracing of tetrode tracks.**

**a,b** Tetrode placement targeting A1 in two mice. The location of primary auditory cortex is indicated with white lines, and labelled A1, assessed by alignment of the slice features to a mouse atlas. Inset: the slice position in the atlas and displacement from bregma.

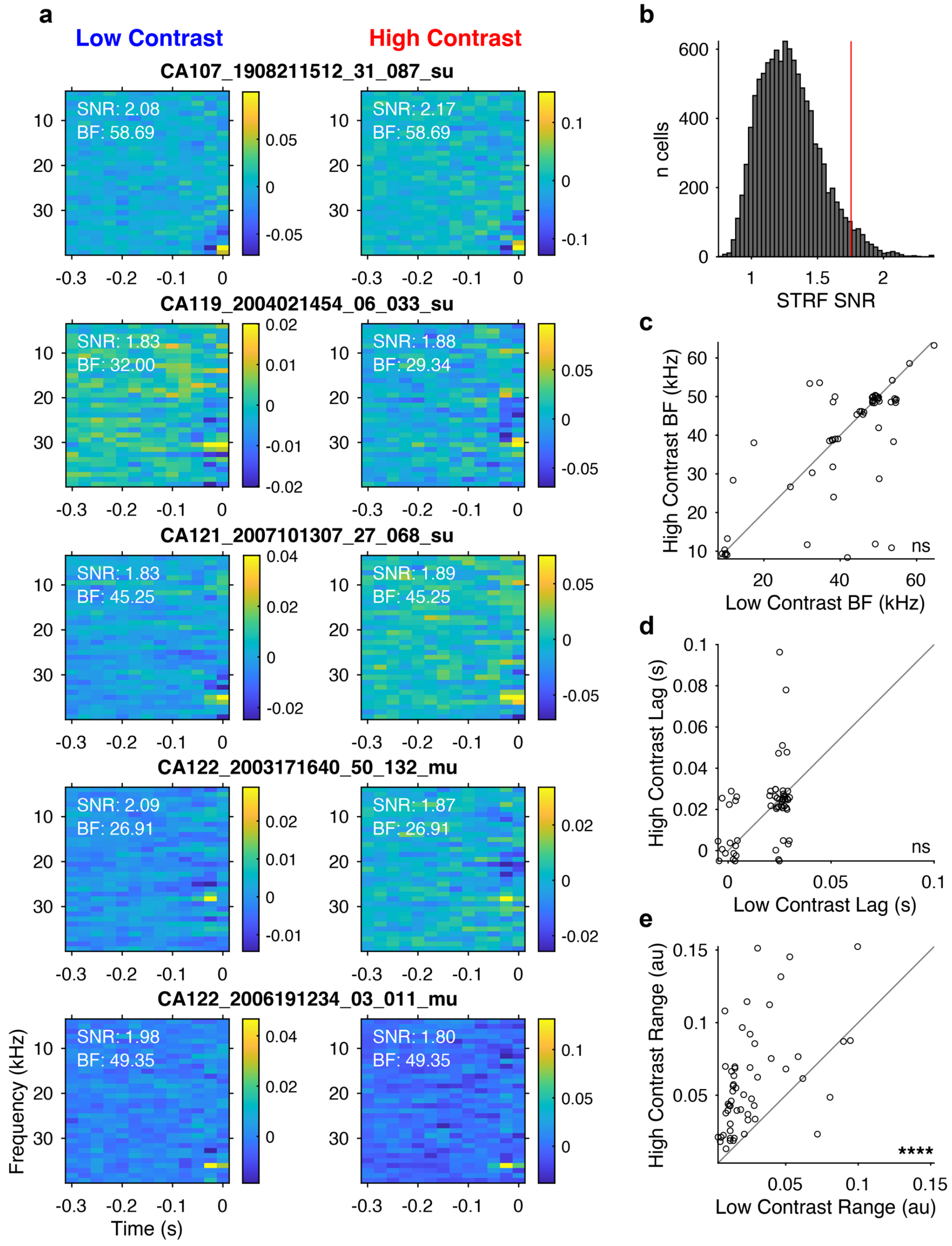


**Supplementary Figure 7 (related to Figure 6). STRFs are unaffected by contrast during behavioral performance.**

**a,** Example STRFs estimated in low and high contrast during behavioral performance (rows are individual neurons, columns are STRFs estimated per contrast, n = 55 neurons). Inset text indicates the SNR of the STRF (*Online Methods*), and the best frequency (BF). **b,** Histogram of SNRs across all neurons and contrasts. We restricted analyses to well-defined STRFs, including only cells with STRFs which were greater than 1.72 SNR (~2std of the SNR distribution) in both low and high contrast. The SNR cutoff is indicated by the red line. **c,** Best frequency of high SNR STRFs in low and high contrast was not affected by contrast (two-way sign-rank test: $p = 0.18$). **d,** The temporal lag of the peak STRF response in low and high contrast was not affected by contrast (two-way sign-rank test: $p = 0.20$). **e,** The range between the minimum and maximum STRF coefficients was affected by contrast (two-way sign-rank test: $p = 8.19\text{e-}9$).

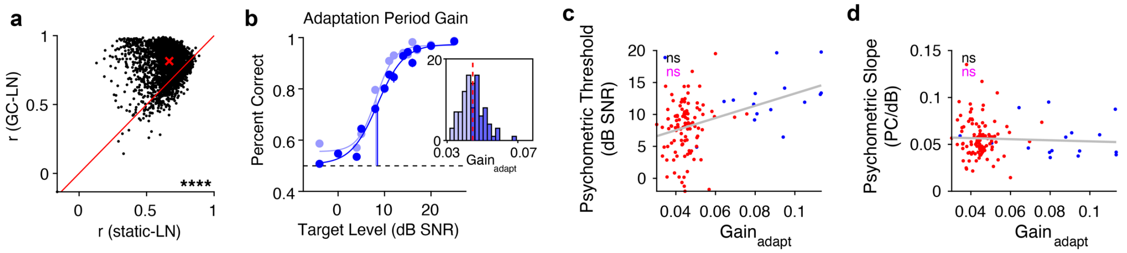


**Supplementary Figure 8 (related to Figure 6). Relationship between behavior and gain outside of the target period.**

**a,** Correlation coefficients between the prediction of a linear-nonlinear model using STRFs estimated from the model without gain control (static-LN) versus a model with gain control (GC-LN). Each dot indicates a neuron. The red solid line indicates unity. The red "x" indicates the median correlation in each contrast. Asterisks indicate the significance of a two-way sign-rank test ($p$ = 1.88e-295). **b,** Psychometric performance in low contrast, averaged based on a median split of average cortical gain during the *adaptation* period of the trial. Light dots and lines indicate the session average and psychometric fit to sessions in the bottom 50$^{th}$ percentile of gain, while dark dots and lines indicate the same values for sessions in the top 50$^{th}$ percentile of gain. Error bars indicate ±SEM across sessions (n=107). *Inset*: distribution of average gain in each session estimated from the adaptation period. The red dashed line indicates the median of the distribution, and the histogram bars are shaded according to whether they fall above (dark blue) or below (light blue) the median. **c,** Session-wise relationship between average gain in the adaptation period and psychometric threshold. Each dot indicates the gain and threshold for a single session, and its color indicates the contrast of the adaptation period. The gray line is the best linear fit to the data. The text in the lower right indicates the results of Likelihood Ratio Tests for models including gain as a predictor (in gray) or contrast as a predictor (in red). Full statistical results in Supplementary Table 1. Grey and black "ns" indicate that gain in the adaptation period and contrast, respectively, did not significantly predict psychometric slopes. **d,** Same as in **j**, but plotting psychometric slope as a function of gain.

### Supplementary Tables

**Supplementary Table 1:** Statistical comparisons.

| Comparison | Figure | Center | Spread | N | Test | Statistic | Effect Size | p-value |
|---|---|---|---|---|---|---|---|---|
| Behavior percent correct, low contrast: time 1 vs. time 2 | 2g | T1: 0.68<br>T2: 0.70<br>(median) | T1: 0.10<br>T2: 0.15<br>(IQR) | 21 mice | Two-tailed Wilcoxon sign-rank test (FDR corrected[94] for multiple comparisons) | Z = -1.93<br>Rank: 60 | $Z/\sqrt{n}$ = -0.42 | 0.054 |
| Behavior percent correct, low contrast: time 1 vs. time 3 | | T1: 0.68<br>T3: 0.82<br>(median) | T1: 0.10<br>T3: 0.092<br>(IQR) | | | Z = -4.01<br>Rank: 0 | $Z/\sqrt{n}$ = -0.88 | 5.96e-5 |
| Behavior percent correct, low contrast: time 1 vs. time 4 | | T1: 0.68<br>T4: 0.87<br>(median) | T1: 0.10<br>T4: 0.190<br>(IQR) | | | Z = -4.01<br>Rank: 0 | $Z/\sqrt{n}$ = -0.88 | 5.96e-5 |
| Behavior percent correct, low contrast: time 1 vs. time 5 | | T1: 0.68<br>T5: 0.91<br>(median) | T1: 0.10<br>T5: 0.11<br>(IQR) | | | Z = -4.01<br>Rank: 0 | $Z/\sqrt{n}$ = -0.88 | 5.96e-5 |
| Behavior percent correct, high contrast: time 1 vs. time 2 | | T1: 0.82<br>T2: 0.77<br>(median) | T1: 0.083<br>T2: 0.19<br>(IQR) | | | Z = 2.84<br>Rank: 181 | $Z/\sqrt{n}$ = 0.62 | 0.0045 |
| Behavior percent correct, high contrast: time 1 vs. time 3 | | T1: 0.82<br>T3: 0.77<br>(median) | T1: 0.083<br>T3: 0.14<br>(IQR) | | | Z = 2.17<br>Rank: 163 | $Z/\sqrt{n}$ = 0.47 | 0.030 |
| Behavior percent correct, high contrast: time 1 vs. time 4 | | T1: 0.82<br>T4: 0.78<br>(median) | T1: 0.083<br>T4: 0.16<br>(IQR) | | | Z = 3.36<br>Rank: 195 | $Z/\sqrt{n}$ = 0.73 | 7.80e-4 |
| Behavior percent correct, high contrast: time 1 vs. time 5 | | T1: 0.82<br>T5: 0.79<br>(median) | T1: 0.083<br>T5: 0.12<br>(IQR) | | | Z = 1.94<br>Rank: 157 | $Z/\sqrt{n}$ = 0.42 | 0.052 |
| ANOVA for effects of pre-post muscimol application, contrast, and level on firing rate in ACtx | S5c | n/a | n/a | 42 neurons | three-way ANOVA | $F_{pre\text{-}post}(1)$ = 812.54<br>$F_{contrast}(1)$ = 22.64<br>$F_{level}(6)$ = 21.70 | $\eta^2$ = 0.38<br>$\eta^2$ = 0.011<br>$\eta^2$ = 0.061 | 4.48e-136<br>2.19e06<br>2.77e-24 |
| ANOVA for effects of pre-post saline application, contrast, and level on firing rate in ACtx | S5d | n/a | n/a | 104 neurons | three-way ANOVA | $F_{pre\text{-}post}(1)$ = 15.40<br>$F_{contrast}(1)$ = 0.43<br>$F_{level}(6)$ = 76.067 | $\eta^2$ = 0.0046<br>$\eta^2$ = 1.29e-4<br>$\eta^2$ = 0.14 | 8.89-5<br>0.51<br>1.76e-88 |
| Percent correct max dB SNR, low contrast: muscimol vs. saline | 3c | Musc.: 0.10<br>Saline: 0.85<br>(median) | Musc.: 0.67<br>Saline: 0.27<br>(IQR) | 10 musc. sessions, 10 saline sessions (4 mice) | Two-tailed Wilcoxon rank-sum test | Z = -2.76<br>Rank: 68 | $Z/\sqrt{n}$ = -0.62 | 0.0058 |
| Threshold (dB SNR), low contrast: muscimol vs. saline | | Musc.: 14.78<br>Saline: 9.66<br>(median) | Musc.: 18.46<br>Saline: 6.88<br>(IQR) | | | Z = 0.72<br>Rank: 115 | $Z/\sqrt{n}$ = 0.16 | 0.47 |
| FA rate, low contrast: muscimol vs. saline | | Musc.: 0.026<br>Saline: 0.132<br>(median) | Musc.: 0.10<br>Saline: 0.85<br>(IQR) | | | Z = -2.91<br>Rank: 66 | $Z/\sqrt{n}$ = -0.65 | 0.0036 |
| Max slope (PC/dB), low contrast: muscimol vs. saline | | Musc.: 0.026<br>Saline: 0.072<br>(median) | Musc.: 0.056<br>Saline: 0.030<br>(IQR) | | | Z: -2.68<br>Rank: 69 | $Z/\sqrt{n}$ = -0.60 | 0.0073 |
| Percent correct max dB SNR, high contrast: muscimol vs. saline | | Musc.: 0.06<br>Saline: 0.80<br>(median) | Musc.: 0.10<br>Saline: 0.85<br>(IQR) | 13 musc. sessions, 10 saline sessions (4 mice) | | Z = -4.06<br>Rank: 92 | $Z/\sqrt{n}$ = -0.83 | 4.96e-5 |
| Threshold (dB SNR), high contrast: muscimol vs. saline | | Musc.: 16.77<br>Saline: 18.80<br>(median) | Musc.: 21.33<br>Saline: 5.89<br>(IQR) | | | Z = -0.35<br>Rank: 156 | $Z/\sqrt{n}$ = -0.071 | 0.73 |
| FA rate, low contrast: muscimol vs. saline | | Musc.: 0.027<br>Saline: 0.213<br>(median) | Musc.: 0.10<br>Saline: 0.85<br>(IQR) | | | Z = -3.19<br>Rank: 107 | $Z/\sqrt{n}$ = -0.65 | 0.0014 |

| | | | | | | | | |
|---|---|---|---|---|---|---|---|---|
| Max slope (PC/dB), high contrast: muscimol vs. saline | | Musc.: 0.012<br>Saline: 0.058<br>(median) | Musc.: 0.024<br>Saline: 0.018<br>(IQR) | | | Z = -3.77<br>Rank: 97 | $Z/\sqrt{n}$ = -0.77 | 1.66e-4 |
| Percent correct max dB SNR, target in high contrast : muscimol vs. saline | 3f | Musc.: 0.07<br>Saline: 0.82<br>(median) | Musc.: 0.51<br>Saline: 0.095<br>(IQR) | 5 musc. sessions, 5 saline sessions (2 mice) | Two-tailed Wilcoxon rank-sum test | Z = nan<br>Rank: 15 | $Z/\sqrt{n}$ = nan | 0.0079 |
| Percent correct at threshold, target in high contrast: muscimol vs. saline | | Musc.: 0.03<br>Saline: 0.53<br>(median) | Musc.: 0.35<br>Saline: 0.11<br>(IQR) | | | Z = nan<br>Rank: 17 | $Z/\sqrt{n}$ = nan | 0.032 |
| FA rate, target in high contrast : muscimol vs. saline | | Musc.: 0.12<br>Saline: 0.23<br>(median) | Musc.: 0.22<br>Saline: 0.11<br>(IQR) | | | Z = nan<br>Rank: 21 | $Z/\sqrt{n}$ = nan | 0.22 |
| Max slope (PC/dB), target in high contrast : muscimol vs. saline | | Musc.: 0.038<br>Saline: 0.057<br>(median) | Musc.: 0.046<br>Saline: 0.012<br>(IQR) | | | Z = nan<br>Rank: 19 | $Z/\sqrt{n}$ = nan | 0.095 |
| Percent correct max dB SNR, target in silence: muscimol vs. saline | | Musc.: 0.85<br>Saline: 0.92<br>(median) | Musc.: 0.23<br>Saline: 0.15<br>(IQR) | 8 musc. sessions, 8 saline sessions (2 mice) | | Z = nan<br>Rank: 53 | $Z/\sqrt{n}$ = nan | 0.13 |
| Percent correct at threshold, target in silence : muscimol vs. saline | | Musc.: 0.11<br>Saline: 0.22<br>(median) | Musc.: 0.28<br>Saline: 0.22<br>(IQR) | | | Z = nan<br>Rank: 55 | $Z/\sqrt{n}$ = nan | 0.20 |
| FA rate, target in silence : muscimol vs. saline | | Musc.: 0.029<br>Saline: 0.041<br>(median) | Musc.: 0.038<br>Saline: 0.11<br>(IQR) | | | Z = nan<br>Rank: 60 | $Z/\sqrt{n}$ = nan | 0.44 |
| Max slope (PC/dB), target in silence : muscimol vs. saline | | Musc.: 0.028<br>Saline: 0.031<br>(median) | Musc.: 0.015<br>Saline: 0.0048<br>(IQR) | | | Z = nan<br>Rank: 63 | $Z/\sqrt{n}$ = nan | 0.65 |
| Correlation coefficient between behavioral and neuronal percent correct (split by target level):<br><br>Low level | 5h | n/a | n/a | 215 sessions | Pearson correlation | Pearson r = 0.31 | n/a | 3.44e-5 |
| Medium level | | n/a | n/a | 194 sessions | | Pearson r = 0.28 | n/a | 6.18e-5 |
| High level | | n/a | n/a | 203 sessions | | Pearson r = 0.15 | n/a | 0.032 |
| Neuronal percent correct, low contrast: time 1 vs. time 2 | 5l | T1: 0.79<br>T2: 0.83<br>(median) | T1: 0.15<br>T2: 0.22<br>(IQR) | 43 sessions | Two-tailed Wilcoxon sign-rank test (FDR corrected[94] for multiple comparisons) | Z = -1.12<br>Rank: 418 | $Z/\sqrt{n}$ = -0.17 | 0.26 |
| Neuronal percent correct, low contrast: time 1 vs. time 3 | | T1: 0.79<br>T3: 0.85<br>(median) | T1: 0.15<br>T3: 0.15<br>(IQR) | | | Z = -3.61<br>Rank: 198 | $Z/\sqrt{n}$ = -0.56 | 0.00031 |
| Neuronal percent correct, low contrast: time 1 vs. time 4 | | T1: 0.79<br>T4: 0.92<br>(median) | T1: 0.15<br>T4: 0.20<br>(IQR) | | | Z = -4.68<br>Rank: 103 | $Z/\sqrt{n}$ = -0.72 | 2.89e-6 |
| Neuronal percent correct, low contrast: time 1 vs. time 5 | | T1: 0.79<br>T5: 0.91<br>(median) | T1: 0.15<br>T5: 0.16<br>(IQR) | | | Z = -5.34<br>Rank: 31 | $Z/\sqrt{n}$ = -0.82 | 9.44e-8 |
| Neuronal percent correct, high contrast: time 1 vs. time 2 | | T1: 0.78<br>T2: 0.74<br>(median) | T1: 0.15<br>T2: 0.12<br>(IQR) | | | Z = 2.62<br>Rank: 690 | $Z/\sqrt{n}$ = 0.40 | 0.0088 |
| Neuronal percent correct, high contrast: time 1 vs. time 3 | | T1: 0.78<br>T3: 0.76<br>(median) | T1: 0.15<br>T3: 0.13<br>(IQR) | | | Z = 1.45<br>Rank: 593 | $Z/\sqrt{n}$ = 0.22 | 0.15 |

| | | | | | | | | |
|---|---|---|---|---|---|---|---|---|
| Neuronal percent correct, high contrast: time 1 vs. time 4 | | T1: 0.78<br>T4: 0.83<br>(median) | T1: 0.15<br>T4: 0.20<br>(IQR) | | | Z = -0.24<br>Rank: 453 | $Z/\sqrt{n}$ = -0.037 | 0.81 |
| Neuronal percent correct, high contrast: time 1 vs. time 5 | | T1: 0.78<br>T5: 0.83<br>(median) | T1: 0.15<br>T5: 0.16<br>(IQR) | | | Z = -2.00<br>Rank: 307 | $Z/\sqrt{n}$ = -0.31 | 0.045 |
| Mixed-effects model:<br>`threshold ~ gain_target + contrast + (contrast-1\|mouse)` | 6g | **Model Coefficients**<br>Estimate ± standard error<br>[tstat(df), p-value]<br><br>Intercept: 10.97±1.33<br>t(120) = 8.27, p = 2.059e-13<br><br>Target gain: -30.46±12.45<br>t(120) = -2.45, p = 0.016<br><br>Contrast: 3.27±1.55<br>t(120)= 2.10, p = 0.038 | | 123 sessions | Likelihood ratio test against model without gain:<br>`threshold ~ contrast + (contrast-1\|mouse)` | $\chi^2(1)$ = 5.82 | | 0.016 |
| | | | | | Likelihood ratio test against model without contrast:<br>`threshold ~ gain_target + (contrast-1\|mouse)` | $\chi^2(1)$ = 3.71 | | 0.054 |
| Mixed-effects model:<br>`slope ~ gain_target + contrast + (contrast-1\|mouse)` | 6h | Intercept: 0.039±0.0064<br>t(120) = 6.23, p = 7.14e-9<br><br>Target gain: 0.16±0.060<br>t(120) = 2.67, p = 0.0085<br><br>Contrast: 0.0094±0.062<br>t(120)= 1.52, p = 0.13 | | 123 sessions | Likelihood ratio test against model without gain:<br>`slope ~ contrast + (contrast-1\|mouse)` | $\chi^2(1)$ = 6.96 | | 0.0083 |
| | | | | | Likelihood ratio test against model without contrast:<br>`slope ~ gain_target + (contrast-1\|mouse)` | $\chi^2(1)$ = 2.28 | | 0.13 |
| Mixed-effects model:<br>`thresh ~ gain_adapt + contrast + (contrast-1\|mouse)` | S8c | Intercept: 5.33±1.64<br>t(120) = 3.26, p = 0.0015<br><br>Adaptation gain: 56.66±35.62<br>t(120) = 1.59, p = 0.11<br><br>Contrast: 2.77±1.92<br>t(120)= 1.44, p = 0.15 | | 123 sessions | Likelihood ratio test against model without gain:<br>`thresh ~ contrast + (contrast-1\|mouse)` | $\chi^2(1)$ = 2.51 | | 0.11 |
| | | | | | Likelihood ratio test against model without contrast:<br>`thresh ~ gain_adapt + (contrast-1\|mouse)` | $\chi^2(1)$ = 2.020 | | 0.16 |
| Mixed-effects model:<br>`slope ~ gain_adapt + contrast + (contrast-1\|mouse)` | S8d | Intercept: 0.062±0.0078<br>t(120) = 7.98, p = 9.63e-13<br><br>Adaptation gain: -0.14±0.17<br>t(120) = -0.80, p = 0.43<br><br>Contrast: 0.0049±0.0084<br>t(120)= 0.58, p = 0.57 | | 123 sessions | Likelihood ratio test against model without gain:<br>`slope ~ contrast + (contrast-1\|mouse)` | $\chi^2(1)$ = 0.64 | | 0.43 |
| | | | | | Likelihood ratio test against model without contrast:<br>`slope ~ gain_adapt + (contrast-1\|mouse)` | $\chi^2(1)$ = 0.33 | | 0.57 |

**Supplementary Table 2:** Mouse strains and genders.

| Experiment | Figures | Strain | N [female, male] |
|---|---|---|---|
| Acute ACtx recordings | Figure 2 | CDH23 | 1 [M] |
| Behavior (no microdrive) | Figure 3 | C57BL/6 x CamKII-cre | 1 [F], 4 [M] |
| | | C57BL/6 x PV-cre | 1 [F] |
| | | CDH23 x SOM-cre | 1 [F], 1 [M] |
| Behavior (microdrive) | Figure 3, Figure 5, Figure 6 | CDH23 | 4 [F], 4 [M] |
| | | C57BL/6 x PV-cre | 1 [F] |
| | | C57BL/6 x SOM-cre | 1 [F] |
| | | CDH23 x SOM-cre | 1 [F], 2 [M] |
| | | CDH23 x CamKII-cre | 1 [F] |
| Muscimol (behavior) | Figure 4 | CDH23 | 2 [F], 2 [M] |
| Muscimol (acute recording) | Supplemental Figure 4 | CDH23 x CamKII-cre | 1 [M] |
| | | CDH23 | 1 [M] |
| Acute ACtx recordings | Supplemental Figure 5 | CDH23 x SOM-cre | 3 [F], 2 [M] |
| | | CDH23 x PV-cre | 1 [F], 1 [M] |
| | | CDH23 x VGAT | 2 [F] |
| | | **Total:** | 19 [F], 19 [M] |

**Supplementary Table 3:** Target SNRs used during psychometric testing.

| Target Levels [range] | [n]: Mouse IDs | n Sessions (total) | n High-Low Contrast Sessions | n Low-High Contrast Sessions |
|---|---|---|---|---|
| 0, 5, 10, 15, 20, 25 dB SNR [25] | [12]: CA102, CA104, CA106, CA107, CA118, CA119, CA121, CA122, CA123, CA124, CA125, CA126 | 214 | 111 | 103 |
| -5, 0, 5, 10, 15, 20 dB SNR [25] | [8]: CA102, CA104, CA106, CA107, CA118, CA119, CA121, CA122 | 31 | 31 | 0 |
| 0, 4, 8, 12, 16, 20 dB SNR [20] | [1]: CA046 | 1 | 0 | 1 |
| 5, 8, 11, 14, 17, 20 dB SNR [15] | [4]: CA118, CA119, CA121, CA122 | 68 | 52 | 16 |
| 8, 10.4, 12.8, 15.2, 17.6, 20 dB SNR [12] | [15]: CA046, CA047, CA048, CA049, CA051, CA052, CA055, CA061, CA070, CA072, CA073, CA074, CA075, CA104, CA107 | 111 | 0 | 111 |
| -4, 0, 4, 8, 12, 16 dB SNR [20] | [11]: CA051, CA052, CA055, CA061, CA070, CA072, CA073, CA074, CA075, CA102, CA106 | 91 | 91 | 0 |
| -5, -1, 3, 7, 11, 15 dB SNR [20] | [5]: CA046, CA047, CA048, CA049, CA051 | 19 | 19 | 0 |
| -75, -60, -45, -30, -15, 0 dB attenuation rel. 25dB SNR | [2]: CA124, CA125 | 20 | n/a | n/a |

**Supplementary Table 4:** GLM simulation parameters

| Parameter | Value |
|---|---|
| μ | 30 |
| $\sigma_L, \sigma_H$ | $[1,3]$ |
| β centroid $m$ (frequency bin $f$, history bin $h$) | $[20,2]$ |
| β covariance matrix $C$ | $\begin{bmatrix} 0.8 & 0.1 \\ 0.1 & 0.5 \end{bmatrix}$ |
| β dimensions $(F \cdot H)$ | $[33, 12]$ |
| Baseline rate $a$ | 0.1 |
| Stimulus scaling $b$ | 1 |
| Gain operating point $c$ | μ |
| Gain control ξ | $[-1.0, -0.5, 0, 0.5, 1.0]$ |
| Adaptation time constants $[\tau_L \ \tau_H]$ | $\begin{bmatrix} Slow - Fast: 0.05 & 0.5 \\ Fast - Fast: 0.5 & 0.5 \\ Fast - Slow: 0.5 & 0.05 \end{bmatrix}$ |
| Simulated background scenes | $100 \text{ } or \text{ } 5$ |
| Contrast history $H'$ | 40 |
| B-spline degree, knots | $[3, 3]$ |

### Supplementary Experimental Procedures

*Acute electrophysiological recordings with muscimol or saline.*

Neuronal signals were recorded from n = 2 awake, untrained mice. Prior to the recording session, each mouse was anesthetized and a headpost and ground pin were implanted on the skull (see *Surgery* in the main text). On the day of the recording, the mouse was briefly anesthetized with 3% isoflurane and a small craniotomy was performed over auditory cortex using a dental drill or scalpel (~1mm x 1mm craniotomy centered approximately 1.25mm anterior to the lambdoid suture along caudal end of the squamosal suture). A 32-channel silicon probe (Neuronexus) was then positioned perpendicularly to the cortical surface and lowered at a rate of 1-2μm/s to a final depth of 800-1200μm. As the probe was lowered, trains of brief noise bursts were repeated, and if stimulus locked responses to the noise bursts were observed, the probe was determined to be in auditory cortex. The probe was then allowed to settle for up to 30 minutes before starting the recording.

For the muscimol and saline recordings (Supplementary Figure 5), a durotomy was performed over the injection site and baseline neuronal responses to the behavioral stimuli were recorded. Then, 2.5μL of .25mg/mL muscimol or 0.9% sterile saline solution was topically applied to the surface of auditory cortex and allowed 30 minutes to penetrate the tissue. The same stimuli were then recorded again after the elapsed time. In these recordings, the same targets and DRC background presented during behavior were presented. Neuronal signals from n = 2 mice (1 mouse for muscimol application, 1 mouse for saline application) were amplified and digitized using a Cheetah Digital LYNX system (Neuronalynx) at a rate of 32kHz.

### Supplementary Results

*The range of presented target levels shapes psychometric performance curves.*

In the experiments presented here, we utilized several sets of target levels to assess psychometric performance (for a summary of the target levels used, see Supplementary Table 3). When computing psychometric curves across all of the target conditions for each mouse (n = 25; Supplementary Figure 4b), we found that detection thresholds were lower in low contrast (Mean dB SNR ($M$) = 8.56, standard deviation ($std$) = 1.81) compared to high contrast ($M$ = 14.23, $std$ = 2.57; paired t-test: $t(23)$ = -8.19, $p$ = 2.88e-8, Supplementary Figure 4c). From our normative model, we expected psychometric slopes to decrease in high contrast. Instead, when combining sessions with different target levels, we found a significant increase in slope during high contrast ($M$ = 0.056, $std$ = 0.018) when compared to low contrast ($M$ = 0.045, $std$ = 0.0052; paired t-test: $t(23)$ = -3.42, $p$ = 0.0024, Supplementary Figure 4d). We hypothesized that psychometric performance was sensitive to the range of targets presented. To test this, we split the data by the range of target levels used in each session, finding that targets drawn from a narrow range resulted in steeper psychometric slopes than targets drawn from a wide range, regardless of the background contrast (Supplementary Figure 4e-h). Therefore, we concluded that the paradoxical increase in psychometric slope was due to the narrow range of targets selected for many high contrast sessions. To control for these effects and thus isolate the effect of background contrast on psychometric slope, we considered only sessions with identical ranges of targets in low and high contrast and found that slopes did indeed decrease in high contrast (Figure 3).

*Muscimol application disrupts cortical encoding of targets.*

In n = 2 awake, naïve mice, we first recorded baseline responses to the stimuli used in the psychometric task, then topically applied muscimol or saline, waited 30 minutes, and recorded stimulus responses again. After muscimol application, there was a marked decrease in neuronal responses to targets compared to the baseline recordings (Supplementary Figure 5b, left). Notably, in our saline control, we observed little to no change in neuronal responses after saline application (Supplementary Figure 5b, right). We next compared how contrast, level and muscimol or saline application changed the responses during the pre- and post-application periods, finding that muscimol significantly reduced the firing rates between pre- and post-application periods, while saline significantly increased firing rates (Supplementary Figure 5c,d, Supplementary Table 1). We speculate that the small increase in firing rate between pre- and post-saline application was due to changes in recording quality or due to neuronal drift over the ~1 hour recording session, and note that the effect size of saline pre-post application is very small ($\eta^2$ = 0.0046) compared to the effect size of muscimol ($\eta^2$ = 0.38). We then used a three-way ANOVA to compare the effects of muscimol, contrast, and target level on target responses in the saline and muscimol recording sessions. We found a significant main effect of muscimol ($F(1)$ = 322.65, $p$ = 4.88e-67) and level ($F(6)$ = 15.48, $p$ = 1.98e-17), but no main effect of contrast ($F(1)$ = 0.39, $p$ = 0.53), indicating nearly complete suppression of responses to both targets and background in high and low contrast (Supplementary Figure 5e,f). These results confirmed that muscimol effectively disrupts the cortical coding of our behavioral stimuli.

*Muscimol application does not prevent licking.*

An additional alternative effect of muscimol is a general loss of the ability to lick. To assess this, we monitored the lick probability of the mice throughout the trial duration, and found that muscimol specifically reduced licking responses during the period where targets were presented (rank-sum test: $Z = -4.23$, $p = 2.34\text{e-}5$; Supplementary Figure 5g, right panel of Supplementary Figure 5h). Mice also tended to lick immediately after the trial onset (Supplementary Figure 5i, green trace), but we found that the lick rates under muscimol and saline conditions were identical during this period (rank-sum test: $Z = 0.23$, $p = 0.81$; Supplementary Figure 4h, left panel). These results suggest that muscimol does not impair the mouse's ability to lick in general, but results in a specific deficit in licking in response to targets.

*STRF are stable across contrasts.*

To interpret the gain changes observed during the behavioral task, we first needed to eliminate the possibility that STRF structure changed across the behavioral epochs of the task (ie, between low and high contrast periods). To assess STRF stability during these two trial periods, we computed STRFs independently in low and high contrast using a GLM. Next, we selected STRFs with well-defined features by computing the SNR of each STRF. SNR was computed by calculating the ratio between the standard deviation of the STRF coefficients in the first 100ms to the SNR of the coefficients in the remaining STRF. This metric assumes that a STRF that is purely noise would have the same variability in the early and late phase of the STRF, while a STRF with stronger stimulus selectivity would have greater variability during the early phase, due to a strong response to some stimulus feature. A sampling of high SNR STRFs are plotted in Supplementary Figure 7a.

Next we selected only neurons with SNRs greater than 1.75 in both contrasts (approximately 2 standard deviations greater than the mean SNR, n = 55 neurons, Supplementary Figure 7b). Within this subset of well-tuned neurons, we then computed the best frequency (BF) and lag by first computing the STRF response averaged over time and frequency, respectively, and multiplied by the standard deviation of coefficients in each bin. The BF and lag were determined to be the peak of the average frequency and time components, respectively. Additionally, we computed the range of the STRFs. As described previously[17,19], we found no significant change in BF (sign-rank test: $Z = 1.34$, $p = 0.18$, Supplementary Figure 7c) or lag ($Z = -1.29$, $p = 0.20$, Supplementary Figure 7d), but did observe a significant increase in STRF range ($Z = -5.76$, $p = 8.19\text{e-}9$, Supplementary Figure 7e). As noted in previous work, the change in STRF range is consistent with a change in neuronal gain. Taken together, these findings demonstrate that contrast, and thus the different behavioral epochs of the task, did not significantly influence STRF features.

**Generalized linear model of contrast gain control dynamics**

A primary goal of the current study was to estimate the influence of stimulus contrast on neuronal gain dynamics, for instance, after a switch from one contrast to another. To approach this problem, we first define a model neuron with dynamic gain control.

*Forward model*

To best approximate the stimuli used in our experiments, we define the stimulus environment of our model as an $F$-dimensional signal that evolves in discrete time steps:

$$X_{t,f} \sim \mathcal{N}(\mu, \sigma_t),$$

where $X_{t,f}$ is a stimulus spectrogram that varies as a function of time $t$ and frequency $f$. Each time and frequency bin of $X$ is sampled from a normal distribution defined by an average value $\mu$ and contrast $\sigma_t$ at time $t$.

To approximate the behavior of real neurons, we define a model neuron that has a two-dimensional linear filter (representing the STRF of the neuron):

$$\beta_{h,f} = \mathcal{N}(m, C;\ h, f),$$

where stimulus filter $\beta_{h,f}$ is defined as a two-dimensional gaussian distribution evaluated at lag $h$ and frequency $f$. The filter location in frequency-history space is defined by its mean $m$ and covariance matrix $C$. The stimulus drive of the neuron at each time step, $x_t$, is then computed as the convolution of the stimulus matrix and the linear filter:

$$x_t = \mathrm{X_t}\beta \tag{1}$$

where $X_t$ at each time $t$ is a row vector of length $F \cdot H$ (ie. the unrolled stimulus spectrogram lagged by $\mathrm{H}$ lags) and $\beta$ is the filter, unrolled as a column vector of the same length.

The model neuron has a firing rate that depends only on the stimulus drive $x_t$ and the contrast $\sigma_t$ at time $t$. We then assume that the number of spikes $y_t$ emitted by the neuron at each time step follow a Poisson distribution:

$$y_t \sim Poisson(\lambda_t)$$

where $\lambda_t$ is the firing rate at time $t$, given by

$$\lambda_t = exp[a + g(\sigma_t)b(x_t - c)] \tag{2}$$

where $g$ is a gain control function, and $a$, $b$, and $c$ are parameters of the model. The parameter $a$ represents the baseline response of the neuron, $b$ is a scaling factor of the stimulus drive, and $c$ represents the operating point of the gain. We remove the obvious degeneracy in the definition of *g* and *b* (only their product matters) by requiring that *g* be adimensional and such that

$$\frac{1}{2}[\mathrm{g}(\sigma_\mathrm{H}) + g(\sigma_L)] = 1 \tag{3}$$

where $\sigma_H$ and $\sigma_L$ are the high and low contrast values. This constraint forces the neutral value of the gain, $g = 1$ to be the midpoint between gain in the high and low contrast conditions.

*Optimal gain control*

In the spirit of the efficient coding principle, we derived a form for $g(\sigma)$ that will guarantee that, under certain conditions, the dynamic range of the neuron will be approximately conserved under changes in contrast. To do this, we define the dynamic range as

$$R(\sigma) = \lambda(\mu + \sigma) - \lambda(\mu - \sigma) \tag{4}$$

which can be rewritten using equation 2 as

$$R(\sigma) = e^a\left[exp\big(g(\sigma)b(\mu + \sigma - c)\big) - exp\big(g(\sigma)b(\mu - \sigma - c)\big)\right]. \tag{5}$$

If the argument of the exponentials is not too large, we can linearize this expression to obtain

$$R(\sigma) \simeq 2e^a b\sigma \cdot g(\sigma) \tag{6}$$

and that $R$ is approximately independent of $\sigma$ provided that $g(\sigma) \propto 1/\sigma$. So, for our model, we set

$$g(\sigma) = \frac{\overline{\sigma}}{\sigma_t} \tag{7}$$

where $\overline{\sigma}$ is the harmonic mean of $\sigma_H$ and $\sigma_L$:

$$\overline{\sigma} \coloneqq \left[\frac{1}{2}\left(\frac{1}{\sigma_H} + \frac{1}{\sigma_L}\right)\right]^{-1} = 2\frac{\sigma_H\sigma_L}{\sigma_H + \sigma_L} \tag{8}$$

Finally, to validate that our fitting methods are sensitive to real world neurons, which do not necessarily adjust their gain to account for changes in contrast according to the model just described, we consider an interpolation scheme that smoothly transforms a model with positive gain control to a similar model without gain control, or with "anti" gain control. To do this, we redefine $g$ as follows:

$$g(\sigma) \rightarrow \xi g(\sigma) + (1 - \xi), \qquad -1 \leq 0 \leq 1 \tag{9}$$

so that by changing $\xi$ we can control whether gain control is optimal ($\xi = 1$), non-existant ($\xi = 0$), or "anti" ($\xi = -1$).

Putting everything together, the final expression for the firing rate of the forward model is

$$\lambda_t = exp\left[a + b\left(\xi\frac{\bar{\sigma}}{\sigma_t} + (1 - \xi)\right)(x_t - c)\right] \tag{10}$$

*Generalized linear model*

The forward model developed in the previous section provides a simple approximation of the relationship between the stimulus, stimulus contrast and neuronal responses. We also note that the form of the forward model lends itself to estimation using a Poisson GLM, provided that the predictors are chosen appropriately. As such, we define the inference model as a Poisson GLM with an intercept term and the following predictors:

$$(x_t - \mu), \qquad \frac{\bar{\sigma}}{\sigma_t}, \qquad (x_t - \mu)\frac{\bar{\sigma}}{\sigma_t}$$

In other words, the model is composed of a stimulus predictor ($x_t - \mu$), a contrast predictor ($\bar{\sigma}/\sigma_t$), and their interaction. Therefore, the GLM models the data at time $t$ as a Poisson distribution with the following mean:

$$\lambda_t = exp\left[\beta_0 + \beta_1(x_t - \mu) + \beta_2\frac{\bar{\sigma}}{\sigma_t}(x_t - \mu) + \beta_3\frac{\bar{\sigma}}{\sigma_t}\right] \tag{10}$$

where $\beta_0 \dots \beta_3$ are the parameters to be inferred, and, as defined previously, $x_t$ is the stimulus drive of the neuron determined by its STRF.

*Model fitting*

To fit the model, we took a two-step approach. First we found the best-fit filter (STRF) for the neuron. Then, we fit the full GLM to determine how the linear drive determined by the STRF is modulated by contrast. In the first step, the linear drive is obtained by fitting the model

$$\ln \lambda_t = \alpha + X_t\beta \tag{11}$$

where $X_t$ is a design matrix defined as a function of frequency bins $f$ and history lags $h$, and $\beta$ is the fitted STRF. Stimulus drive $x_t$ is then computed as in equation 1**.**

We then define the full model according to equation 10,

$$\ln \lambda_t = \beta_0 + \beta_1 x_t + \sum_{i=1}^{B} \beta_{2i}\, x_t \cdot (b_i * c)(t) + \sum_{i=1}^{B} \beta_{3i}(b_i * c)(t) \tag{12}$$

where $c(t) = \bar{\sigma}/\sigma_t$ and $\{b_i\}_{i=1}^{B}$ is a set of cubic B-spline temporal basis functions. By defining a matrix $C$ as follows

$$C_{ti} = (b_i * c)(t) \tag{13}$$

we can rewrite equation 12 in a more compact form:

$$\ln \lambda = \beta_0 + x\beta_1 + x \circ C\beta_2 + C\beta_3 \tag{14}$$

where $\circ$ denotes element-by-element "broadcasting" multiplication.

To fit asymmetric changes in firing rate after transitions to low or high contrast, we took the simple approach of defining separate sets of contrast predictors for each transition type. This amounted to modifying $C$ by masking transitions to high contrast or transitions to low contrast with zeros, such that the model fit a window $H'$ of 40 time bins around each contrast transition. To do so, we created a new matrix $C'$ by duplicating $C$ column-wise. Then, we define the first $B$ columns as predictors for the transition to low contrast by masking a 1 second period around each transition to high contrast with zeros. This same procedure was repeated for the remaining columns in $C'$, instead masking out the transition to low contrast. Substituting this into equation 14, we obtain

$$\ln \lambda = \beta_0 + x\beta_1 + x \circ C'^{\beta_2} + C'^{\beta_3} \quad (15)$$

For the sake of clarity, note that in the expression above, $\beta_0$ is a number, $x$ is a column vector of length $T$, $\beta_1$ is a number, $C'$ is a $T$-by-$2B$ matrix, and $\beta_2$ and $\beta_3$ are column vectors of length $2B$.

*Defining gain*

We have outlined a forward model for simulating neuronal activity according to efficient coding of stimulus contrast, and described an inference model (a Poisson GLM) for estimating the influence of the stimulus, stimulus contrast, and their interaction. In this section, we describe how to use the fitted parameters to quantify the amount of gain control in the neuron.

Conceptually, an increase or decrease in the gain of a system is analogous to more or less sensitivity to small changes in the stimulus, dependent on what is modulating the gain (in our case, the recent history of the contrast). Based on this intuition, we focus on how the response of the neuron (as modeled by a fitted GLM) is expected to change between conditions where the gain is expected to contribute (i.e. in the presence of gain control) and where it is not (ie. in the absence of gain control, where gain is "neutral").

To do this, we start by considering the gradient of the link function (the log rate) at time $t$ with respect to $X_t$:

$$\begin{aligned} \eta_t = \nabla_{x_t} ln\lambda_t &= \nabla_{x_t}[\beta_0 + x_t\beta_1 + x_t C_t \beta_2 + C_t\beta_3] \\ &= \nabla_{x_t}[\beta_0 + (X_t\beta)\beta_1 + (X_t\beta)C_t\beta_2 + C_t\beta_3] \\ &= \beta(\beta_1 + C_t\beta_2) \end{aligned} \quad (16)$$

We can immediately read equation 16 as "the STRF of the model is modulated by a factor of $\beta_1 + C_t\beta_2$ at time $t$", and define the gain based on this intuition, but we will take a slightly longer and more formal route to get to the same result.

The gradient $\eta$ is a vector with the same dimensionality of $\beta_1$ and $C_t\beta_2$, and it encapsulates all information about the sensitivity of the link function to small changes in $X_t$ at a given time. Because $X_t$ is not a scalar (it has $H \cdot F$ components), these changes can happen along many dimensions, and the sensitivity can be different in different directions. We can define the gain based on the sensitivity to changes in a specific direction $v$ (assuming for concreteness that $||v|| = 1$, although this is not necessary for the derivation below). If $X_t = r \cdot v$, where $r$ is some scalar, then

$$\frac{dln\lambda_t}{dr} = \langle \eta_t, v \rangle \quad (17)$$

by definition of the gradient. We can then define the gain $w$ along direction $v$ as the ratio between the sensitivity of the log rate to changes along $v$ and the sensitivity one would have if the contrast $C_t$ was at some reference value $C^0$ where we define $w = 1$ by construction. If we do so, we obtain

$$\begin{aligned} w_t &= \left(\frac{dln\lambda_t}{dr}\right)\left(\frac{dln\lambda(C = C^0)}{dr}\right)^{-1} \\ &= \frac{\langle \beta, v \rangle (\beta_1 + C_t\beta_2)}{\langle \beta, v \rangle (\beta_1 + C^0\beta_2)} \\ &= \frac{\beta_1 + C_t\beta_2}{\beta_1 + C^0\beta_2} \end{aligned} \quad (18)$$

Note that this definition does not depend on the initial choice of $v$, or even on the specifics of the choice of basis functions used to define $C$. In conclusion, by reasoning about the sensitivity of the response of the fitted GLM, we define a value $w_t$ which captures the relationship between the true gain $g$ and the stimulus contrast $c_t$.

*Simulations*

To validate our inference model, we simulated neuronal activity according to the generative model defined in the *Forward Model* section (Supplementary Figure 2a). We were interested in capturing several dimensions upon which the generative model could vary, namely, the amount of gain control in the simulated neurons $\xi$, and the dynamics of the gain function $g$.

To parametrically control the evolution of gain over time, we simulated different temporal trajectories of gain control, by modifying $g(\sigma_t)$ as follows

$$g(\sigma_c, \tau_c)_t = g(\sigma_{c-1}) + (g(\sigma_c) - g(\sigma_{c-1})) \cdot exp(\tau_c, t) \tag{19}$$

where the gain $g$ after a switch to contrast $\sigma_c$ transitions from the gain in the previous contrast $g(\sigma_{c-1})$ to the gain in the current contrast $g(\sigma_c)$ according to an exponential function with time constant $\tau_c$. Note that $\tau_c$ could vary between the two contrasts to simulate asymmetric dynamics.

For each simulated neuron, we first generated a STRF and linear drive according to equation 1 (Supplementary Figure 2b,d). For different sets of simulated neurons, we parametrically varied the amount of gain control $\xi$ between -1 and 1, and varied the gain time courses to simulate three types of gain adaptation dynamics: 1) Slow transitions to low contrast with fast transitions to high contrast, 2) Fast, symmetric transitions to each contrast, 3) Fast transitions to low contrast and slow transitions to high contrast (Supplementary Figure 2f).

We simulated 100 neurons for each combination of $\xi$ and $\tau$, with other simulation parameters held constant (Supplementary Table 4). Supplementary Figure 2e plots the average firing rates and overlaid model fits for three sets of simulations with optimal gain control ($\xi = 1$) while varying $\tau$. Importantly, the model flexibly captured the gain dynamics in the three simulated adaptation conditions, with the gain estimate $w_t$ following the true gain trajectory (Supplementary Figure 2f). For additional values of $\xi$, the model accurately predicted the firing rates (Supplementary Figure 2g) and gain trajectories (Supplementary Figure 2h). We observed that some combinations of $\xi$ and $\tau$ elicited large firing rate transients, particularly in the cases where simulated gain slowly adapted after a switch to high contrast (bottom panels in Supplementary Figure 2e, f, g, h). This behavior is expected, as gain remains relatively high for a longer period after the switch, causing large fluctuations in firing rate as the stimulus drive during high contrast is increased. These large firing rate transients seemed to reduce the accuracy of gain estimate $w$, but we observed that the predicted time courses still captured the overall asymmetries present in the underlying model.

During our behavioral recordings, we used a limited number of background noise scenes (n = 5) to reduce the overall size of the stimulus set. However, it became clear that our model required a larger sample of stimulus space to accurately estimate gain. To demonstrate this, we plotted the simulation results when neurons were exposed to 100 unique background scenes (Supplementary Figure 2i) compared to simulations where neurons were only exposed to 5 unique background scenes, as in our behavioral recordings (Supplementary Figure 2j). We observed that with 100 scenes, estimates of $w$ were very close to the true gain values, but were consistently underestimated in the case of 5 background scenes, even in the case of perfect gain control. Therefore, when analyzing our behavioral recordings, we used a standard linear-nonlinear model to estimate neuronal gain (Figure 5), as we previously found that gain estimates from the GLM were highly correlated with gain estimated from the LN model (Figure 2i).

### Graphical Abstract

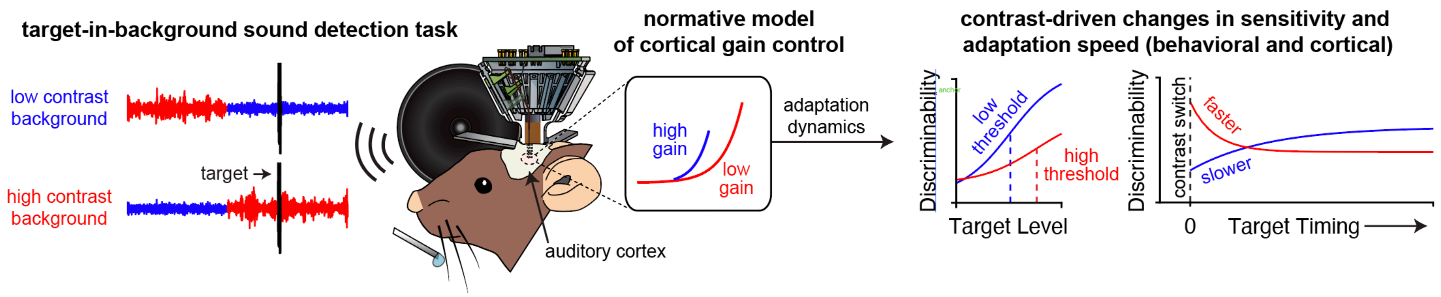
