## Supplementary figures and images for "Dynamics of cortical contrast adaptation predict perception of signals in noise"

### Figure 1

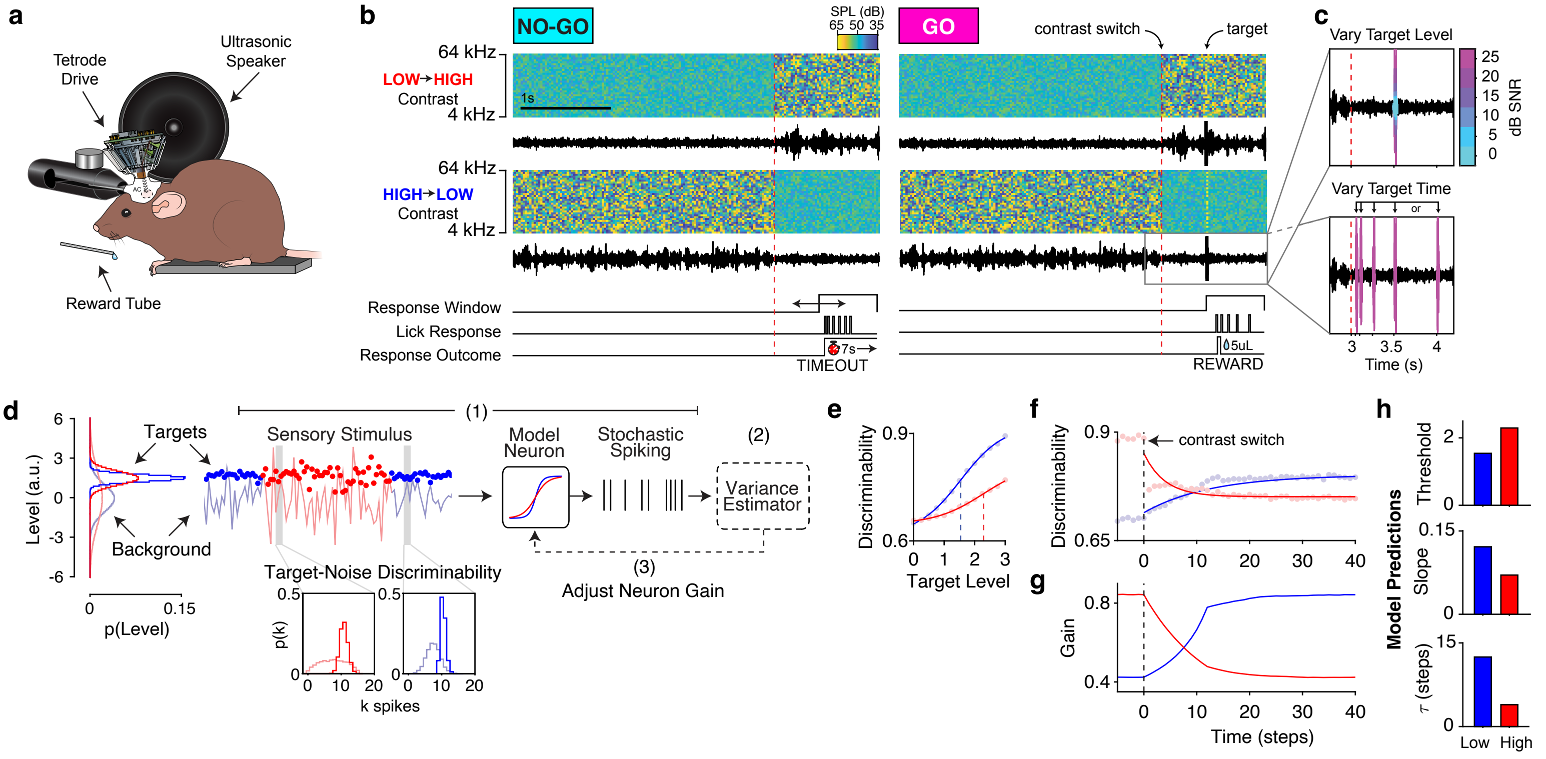

### Figure 2

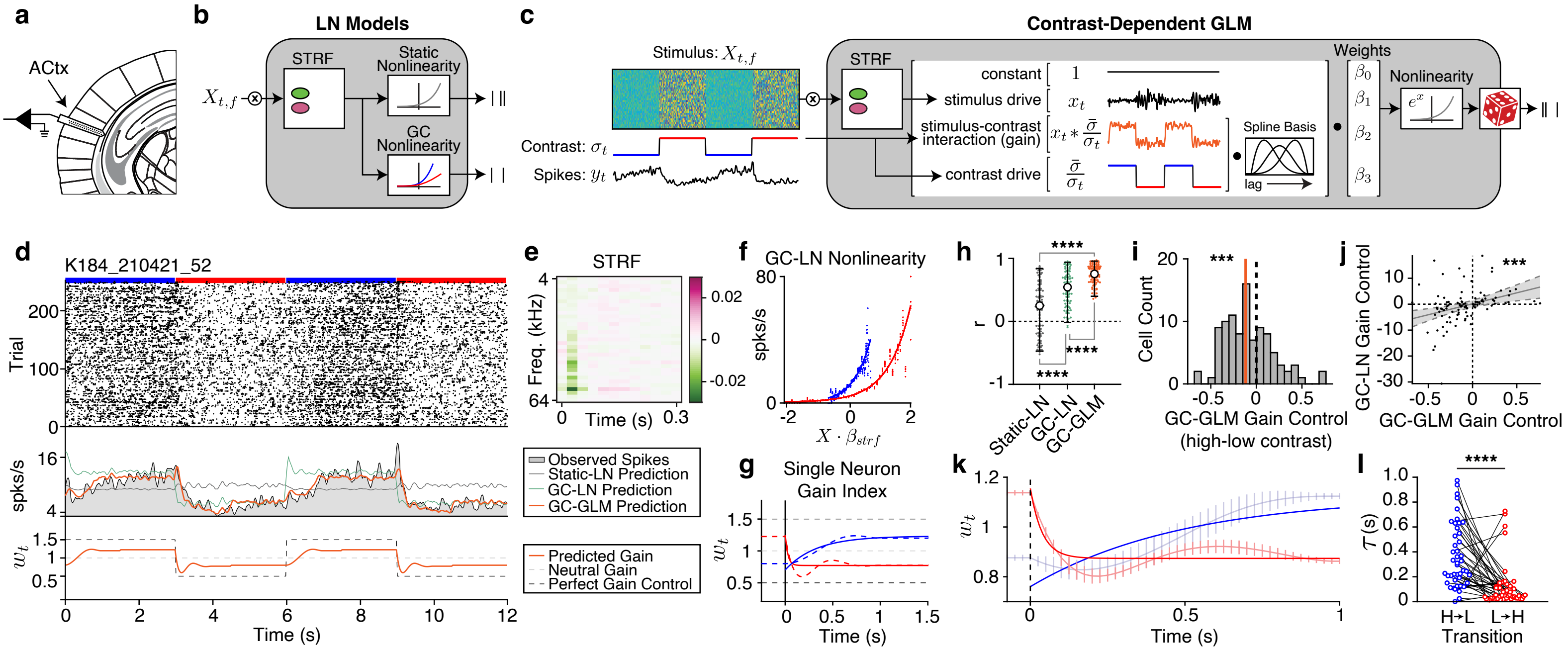

### Figure 3

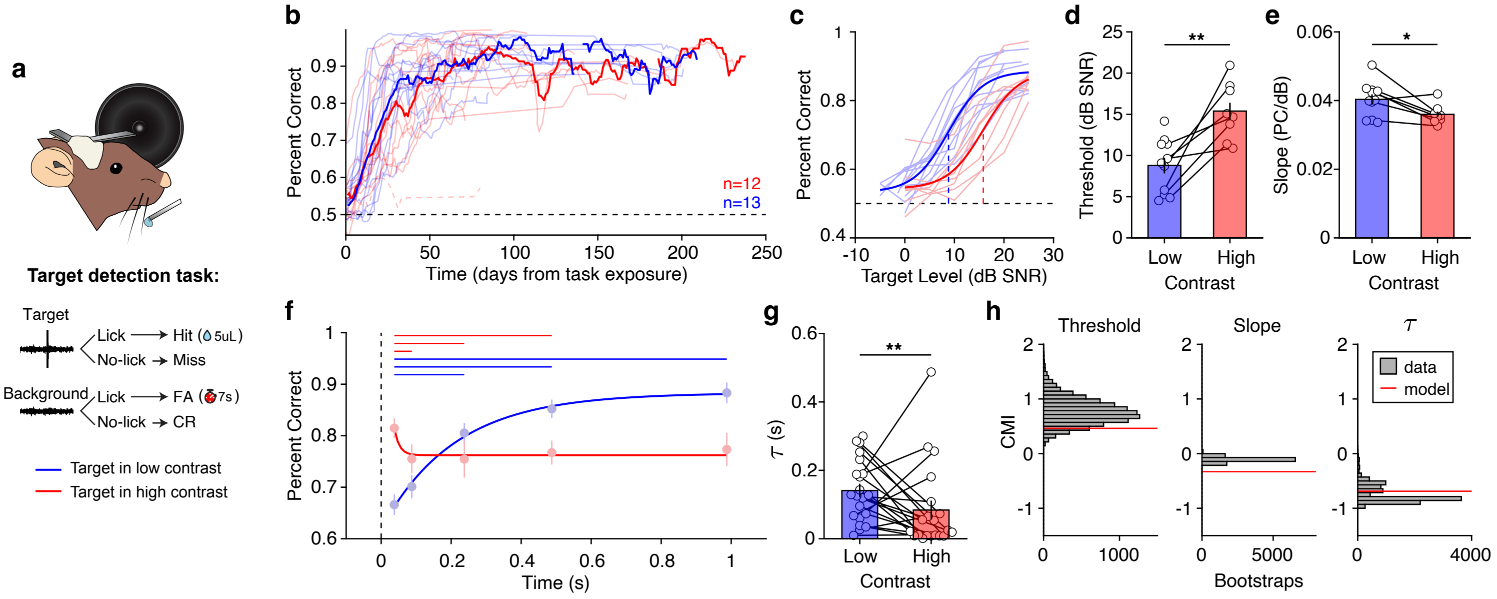

### Figure 4

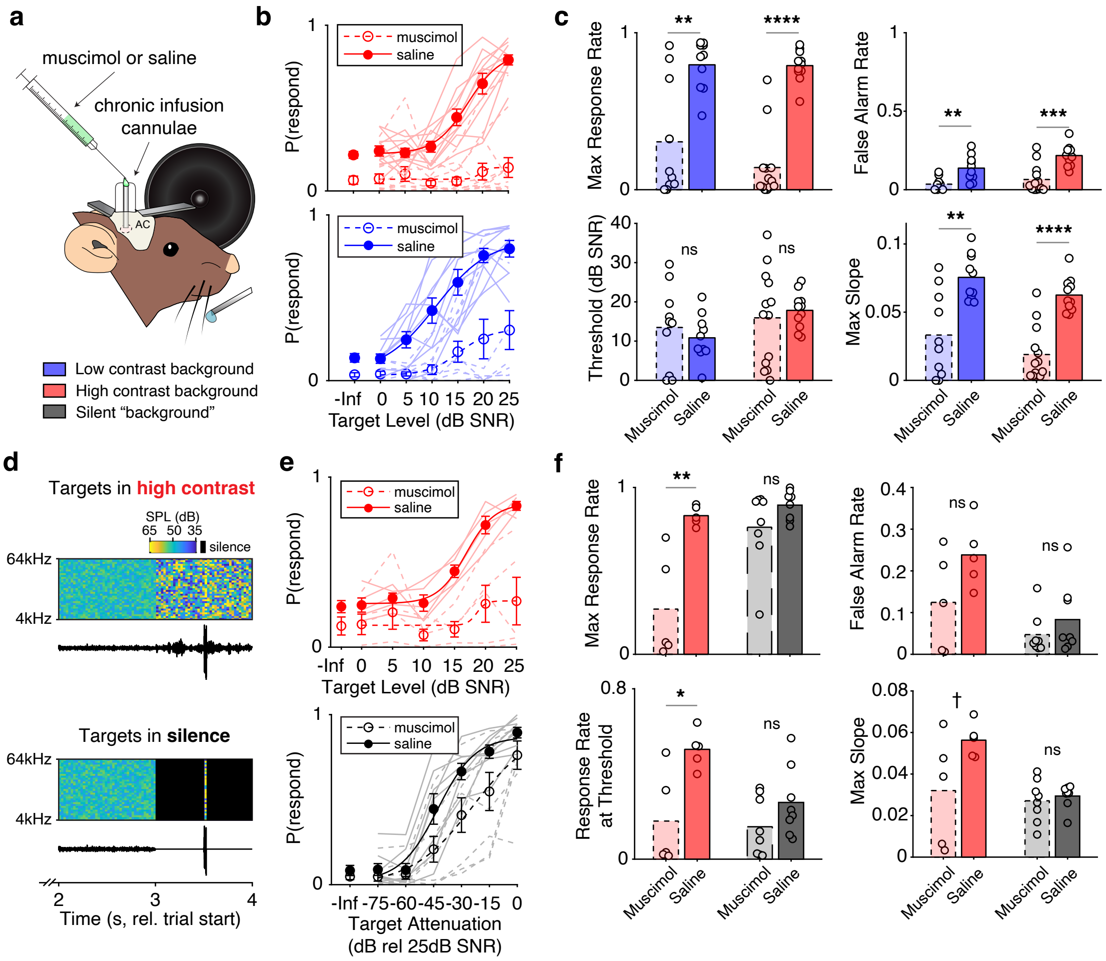

### Figure 5

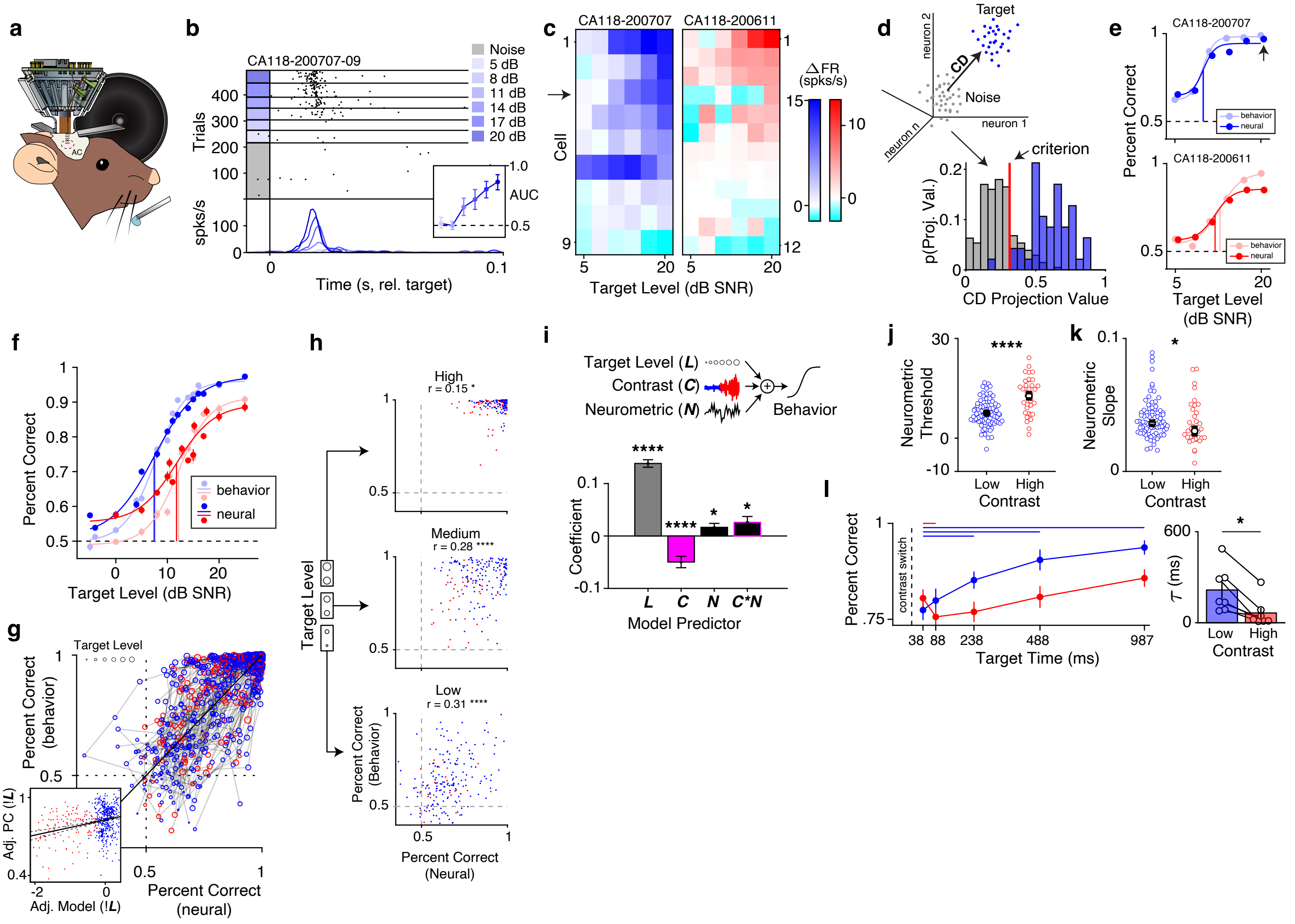

### Figure 6

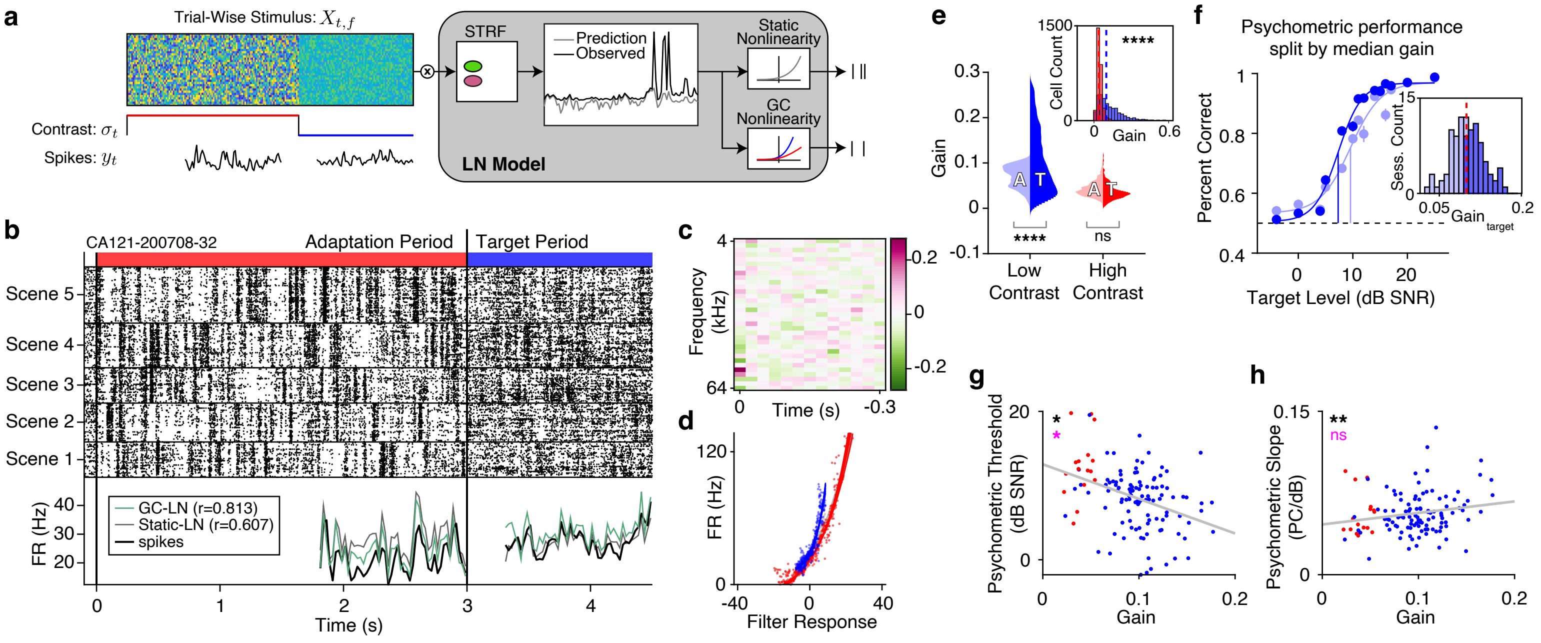

### Figure S1

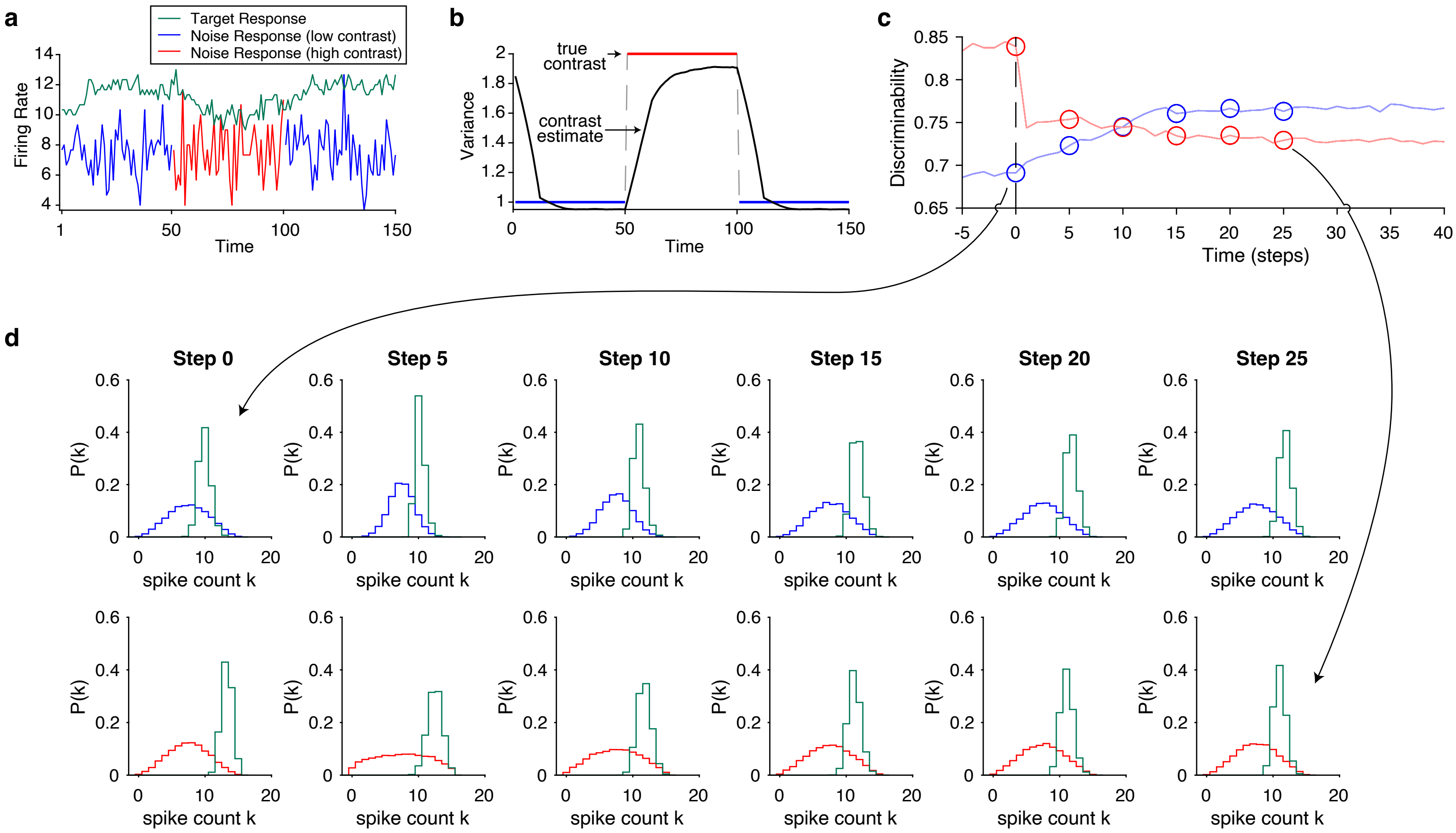

### Figure S2

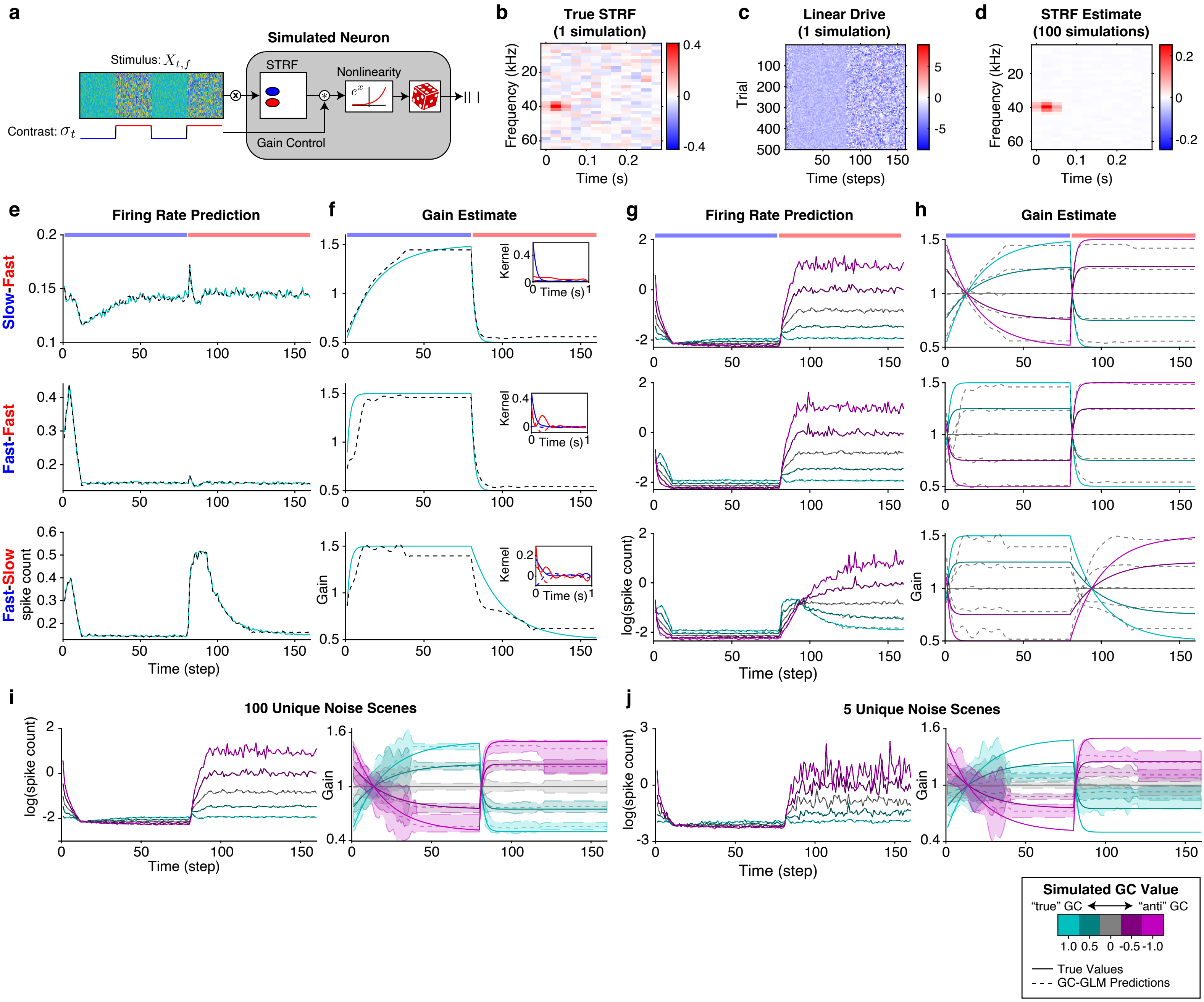

### Figure S3

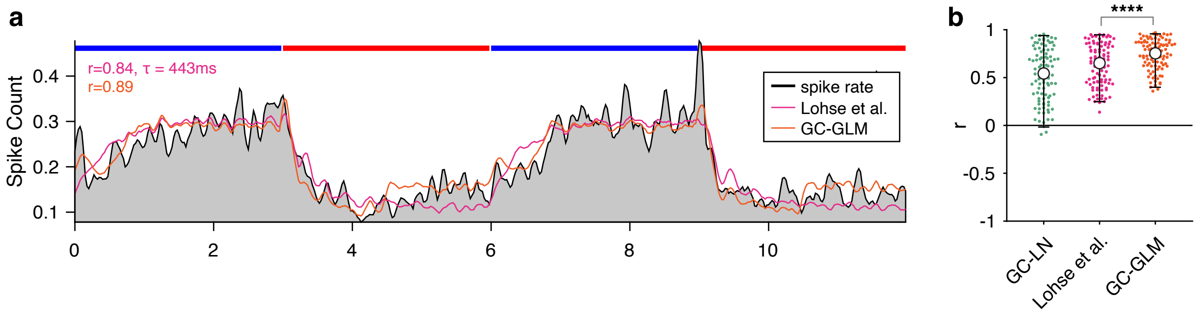

### Figure S4

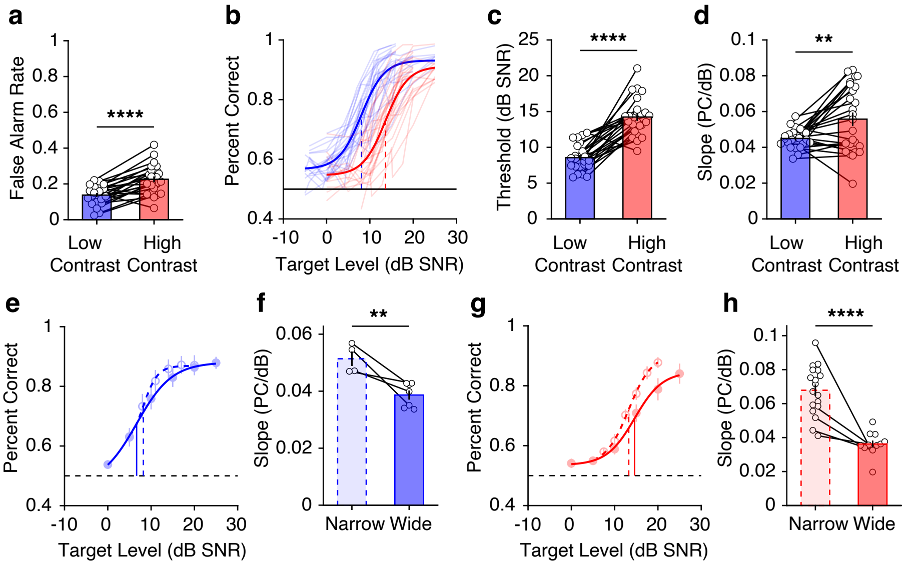

### Figure S5

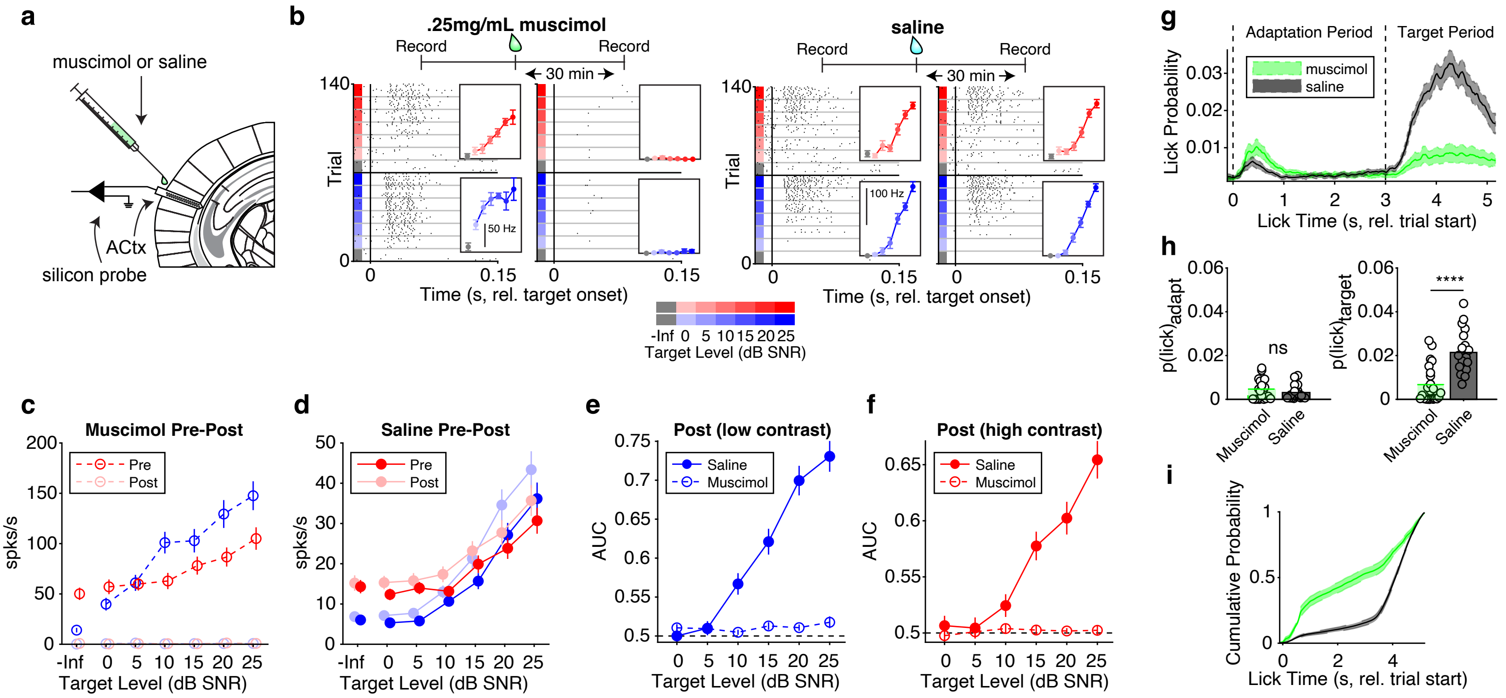

### Figure S6

a

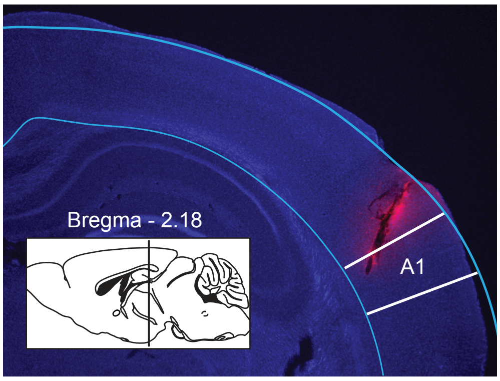


b

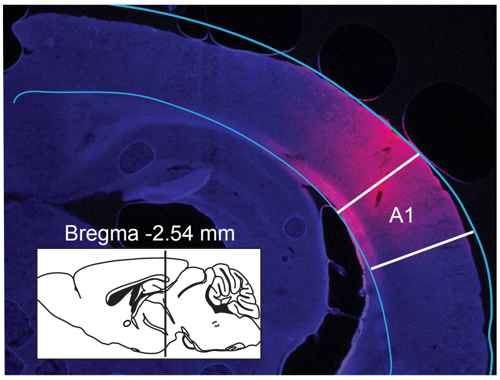

### Figure S7

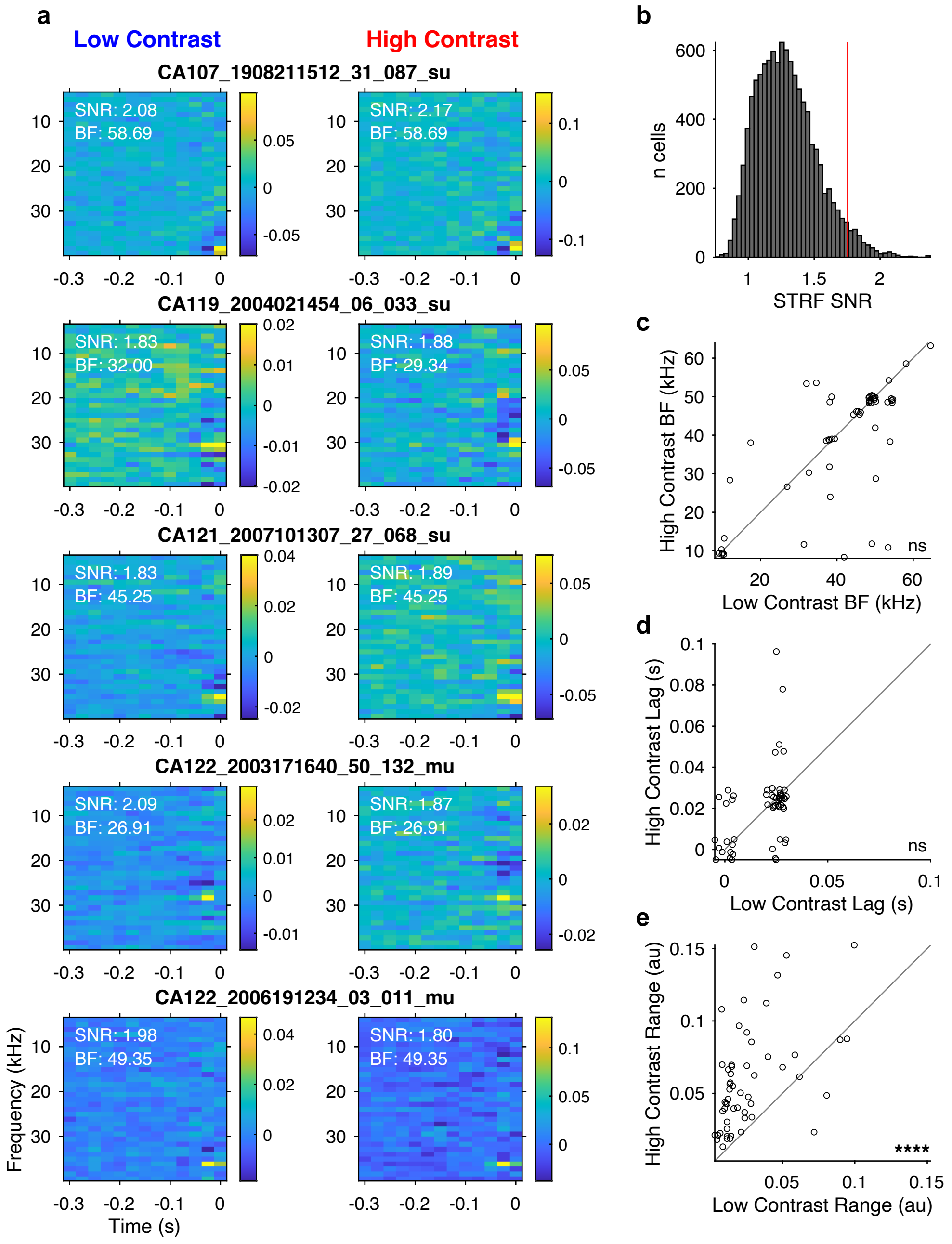

### Figure S8

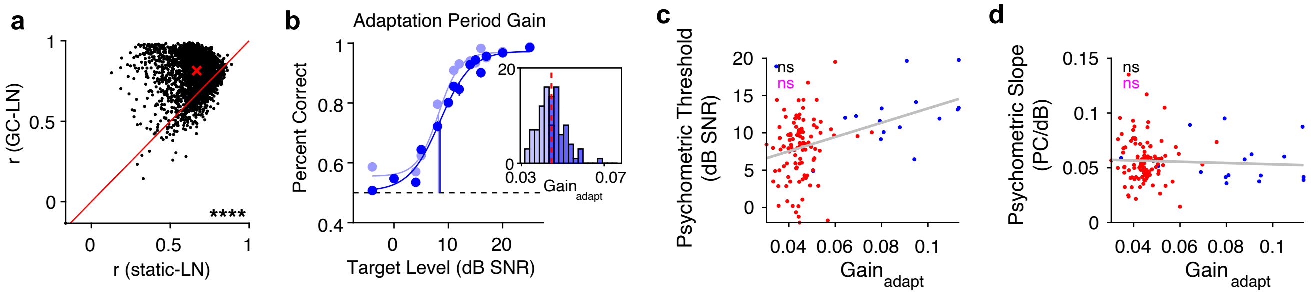
